# Molecular Epidemiology and Mechanisms of Antibiotic Resistance in Gram-positive Bacteria in Africa: A Systematic Review and Meta-Analysis from a One Health Perspective

**DOI:** 10.1101/366807

**Authors:** John Osei Sekyere, JEric Mensah

## Abstract

A systematic review and meta-analysis of antibiotic-resistant Gram-positive bacteria in Africa, showing the molecular epidemiology of resistant species from animal, human and environmental sources, is lacking. Thus, the current burden, type, and sources of Gram-positive bacterial resistance and their dissemination routes from farm to fork is absent. To fill this One Health information gap, we systematically searched PubMed, Web of Science and African Journals Online for English research articles reporting on the resistance mechanisms and clonality of resistant Gram-positive bacteria in Africa within 2007 to 2018. The review and all statistical analysis were undertaken with 130 included articles.

From our analyses, the same resistant Gram-positive bacterial clones, resistance genes, and mobile genetic elements (MGEs) are circulating in humans, animals and the environment. The resistance genes, *mecA, erm(*B), *erm*(C), *tet*(M), *tet*(K), *tet*(L), *vanB, vanA, vanC,* and *tet*(O), were found in isolates from humans, animals and the environment. Commonest clones and mobile genetic elements identified from all three sample sources included *Staphylococcus aureus* ST5 (n=208 isolates), ST 8 (n=116 isolates), ST 80 (n=123 isolates) and ST 88 (n=105 isolates), and IS*16* (n=18 isolates), Tn*916* (n=60 isolates) and SCC*mec* (n=202 isolates). Resistance to penicillin (n=4 224 isolates, 76.2%), erythromycin (n=3 552 isolates, 62.6%), ampicillin (n=1 507 isolates, 54.0%), sulfamethoxazole/trimethoprim (n=2 261 isolates, 46.0%), tetracycline (n=3 054 isolates, 42.1%), vancomycin (n=1 281 isolates, 41.2%), streptomycin (n=1 198 isolates, 37.0%), rifampicin (n=2 645 isolates, 33.1%), ciprofloxacin (n=1 394 isolates, 30.5%), clindamycin (n=1 256 isolates, 29.9%) and gentamicin (n=1 502 isolates, 27.3%) (*p*-value <0.0001) were commonest.

Mean resistance rates of 14.2% to 98.5% were recorded in 20 countries within the study period, which were mediated by clonal, polyclonal and horizontal transmission of resistance genes. A One Health approach to research, surveillance, molecular epidemiology, and antibiotic stewardship to contain ABR should be prioritized.

## HIGHTLIGHTS

- Gram-positive bacteria (GPB) isolated from human, animal and environmental samples were of the same clones and/or shared common resistance genes and mobile genetic (MGEs).
- Multidrug resistant (MDR) clones such as *S. aureus* ST5 and *E. faecium* ST80 were isolated from human, animal and environmental sources.
- *mecA, erm*(B), *erm(C*) *tet(*M/K/L), and *vanA/B/C* were common in GPB, including VRSA.
- Mean drug resistance rates of isolates from humans, animals and the environment were respectively 62.0% (95% CI: 54.7 - 69.3%), 68.2% (95% CI: 58.0 -78.4%) and 84.6% (95% CI: 69.9 – 99.31%) (*P-value* < 0.0001).
- *SCCmec,* IS*16,* and *Tn916* mobilized *mecA, erm*(B) and *tet*(M) respectively across various GPB species isolated from animals, humans, and the environment.
- A One Health approach to studying antibiotic resistance mechanisms and molecular epidemiology of GPB is warranted.

## 1. INTRODUCTION

### Antibiotic resistance, a threat to public health

Limited research and surveillance data in Africa makes it impossible to track and monitor the true burden of antibiotic resistance (ABR) ^1^, particularly the distribution and dissemination of resistance genes between humans, animals and the environment. According to a recent WHO report, the potential for ABR to lead to higher mortalities and morbidities in low- and middle-income countries such as Africa may even be greater as a result of the higher burden of bacterial infections, limited diagnostic capacity and lower access to second-line antibiotics^1,2^. This makes it imperative to have a One Health analysis that describes the burden and epidemiology of resistance genes in bacteria isolated from humans, animals and the environment ^3^.

In a recent review, Gram-positive bacteria (GPB) were responsible for a high proportion of infections among children and showed a high level of resistance to WHO-recommended drugs in Africa ^4^. In some African regions, as many as 80% of *Staphylococcus aureus* infections are methicillin-resistant *S. aureus* (MRSA), which show resistance to most standard licensed drugs including quinolones and peptides ^25^. Although *Enterococcus spp.* are mostly not as virulent as *S. aureus,* their multidrug resistance (MDR) propensities restrict drug options for clinicians ^7^. Patients infected with MRSA are estimated to be 64% more likely to demise than those infected with methicillin-susceptible *S. aureus* (MSSA) ^6^.

Reviews addressing GPB in Africa have reported on increasing rates of ABR from blood-stream infections, pneumonia, urinary tract infections and meningitis caused by *Streptococcus agalactiae, S. aureus, Streptococcus pneumoniae* and *Enterococcus faecium* in both children and adults. Sepsis due to *S. agalactiae* accounts for about 26% of all neonatal deaths and 10% maternal deaths in Sub-Saharan Africa ^8^ However, the potential dissemination of these resistant strains from farm (environment and animals) to fork (humans), are less described.

### Sources and anthropogenic activities driving resistance

High-level ABR has been reported in humans, animals and the environment, with indiscriminate antibiotic use being fingered as a major contributor in Africa. Resistance genes have been detected in surface water fed with runoff effluents from farms utilizing antibiotics, hospitals, and sewage processing plants as well as in ground water ^9-11^. Furthermore, genes mediating resistance to last-resort GPB antibiotics such as vancomycin have been recovered from raw milk and animal products, pigs, wild animals (buffalo, zebra and cattle), waste water, effluents and patients, implicating veterinary and agricultural use of antibiotics as potential sources of resistance genes in humans ^12-14^. These reports suggest that a larger share of the antibiotics that end up polluting the environment and communities emanate from livestock production ^15-17^. This interconnectivity between animals, humans and the environment, explains the need to adopt a One Health research policy.

Several studies have reported high rates of MDR among GPB isolates from humans, animals and the environment in Africa, mainly as a result of overuse, underuse and wrong choice of antibiotics ^18-24^. Different factors have been implicated in the high rate of ABR to the limited drugs in Africa. These include: unrestricted access to antibiotics over-the-counter without prescription such as selling on the streets; inadequate hygienic practices; uncontrolled usage of antibiotics as growth promoters in food animals production; wrong diagnosis and prescription, off-label use and errors in dosage regimens; use of untreated poultry and cattle manure to fertilize agriculture lands; extensive use of broad-spectrum antibiotics in poultry production; and inefficient chlorination of hospital wastewater effluents before discharge into the environment^10,18,22,25-29^. Additionally, inadequate knowledge of animals’ diseases, misdiagnosis and poor antibiotic handling practices in animal production add up to the overall burden of ABR in Africa ^17^.

### Molecular ABR mechanisms

Selective pressures exerted by various antibiotics used in human and veterinary medicine, as well as in agriculture, have resulted in the emergence and dissemination of numerous mechanisms of resistance in GPB in Africa. Commonly reported mechanisms include *blaZ, erm(*B), *mecA, tet*(M), *vanB* and *vanC* ^30-33^. These resistance genes have been found to be associated with mobile genetic elements (MGEs) such as transposons, conjugative plasmids, integrons, and insertion sequences, which are capable of mobilizing resistance genes across a wide spectrum of bacterial species ^34,35^. *SCCmec, Tn916* and IS16 are notable MGEs that carry major ABR determinants in Africa and are transmissible between clones of the same or different bacteria species by a conjugative mechanism. These MGEs have the potential to thus spread resistance genes from environmental and animal bacterial hosts to human pathogens in Africa; they have therefore been analysed herein ^36-38^.

### Purpose of this review

Excellent reviews addressing antimicrobial resistance in some GPB and Gram-Negative ones in Africa have been published ^4,39-44^ However, reviews discussing the molecular epidemiology and mechanisms of ABR in GPB such as *Staphylococcus spp., Streptococcus spp.* and *Enterococcus spp.* in Africa in the context of resistance rates, resistance mechanisms (and MGEs), clonality, and geographical distribution from a One Health perspective are non-existent, to the best of our knowledge. This review sought to fill this gap by analyzing the burden, types, and molecular epidemiology of resistant GPB from a One Health context.

### 1.1 Search strategy and inclusion criteria

English research articles published within the last ten years (01/01/2007 to 07/08/ 2018) and indexed in PubMed, Web of Science and African Journals Online were searched with the following keywords: “Enterococcus”, and “Streptococcus”, “Staphylococcus”, in permutations and combinations with “resistance AND Africa”. Studies which did not identify the underlying ABR mechanisms/genes as well as the clonality of antibiotic-resistant GPB were excluded. Thus, studies that only reported on antibiotic sensitivity testing (AST) results or undertook ABR surveillance studies without further molecular tests to characterize the ABR mechanisms and/or clonality of the isolates were excluded (Figure 1). In all, 248 studies were excluded because they only had MIC data (See Supplementary data 1). All searches were undertaken independently by both authors in triplicates to ensure replication of the results.

Data extracted from the articles included year of study, country, GPB species, clones, sample sources, sample size/number of isolates, number of resistant isolates, resistance genes and MGEs and antibiotics to which the strains were resistant (Tables 1-6; Supplementary data 2). The mean rate of ABR among GPB per country and in Africa was determined to identify countries with the highest or lowest levels of resistance in Africa (Table 5). As well, the antibiotics to which the isolates were most resistant were determined to evaluate their correlation with the detected/reported resistance mechanisms (Table 6).

The resistance mechanisms, as well as MGEs involved in the transmission of resistance genes per species or clone, were determined to assess the means of resistance transfer i.e., horizontal or vertical (through clonal expansion), per specimen sources (animal, human, and environment) (Figures 2a & 2b). The distribution of clones, resistance genes, and MGEs were considered to identify countries with most resistant clones, resistance genes, and their associated MGEs (Figure 3a).

### 1.2 Statistical analysis

The data was analyzed using Microsoft Excel^®^ 2017 and Graph pad prism™ 6 (GraphPad Software, San Diego, CA, USA) (Supplementary data 2). Calculation for the statistical significance of the data was determined using the kolmogorov-smirnov test (with Dallal - wilkinson-Lilliefors p-value) and/or column statistics or one sample t-test, and the confidence intervals determined at 95%. The p-values were two tailed with a Gaussian approximation. A p-value of <0.05 was considered as statistically significant. Only studies that provided the required information were used in the analysis. In all, 130 articles were used for the data analysis (Fig. 1).

## 2. RESULTS AND DISCUSSION

Of the 1,486 articles returned from the systematic literature search from PubMed, Web of Science and African Journals Online, 130 studies representing 20 out of 54 African countries were included in this review and data analysis (Fig. 1). A total of 249 papers were excluded because they only had MIC data. Tunisia (n=33 studies) recorded the highest number of studies followed by South Africa (n=21 studies), Egypt (n=21 studies), Nigeria (n=13 studies) and Algeria (n=7 studies), Angola (n=6 studies), Uganda (n=5 studies), Democratic Republic of the Congo (n=3 studies), Ghana (n=3 studies), Kenya (n=3 studies), São Tomé and Príncipe (n=3 studies), Gabon (n=2 studies), Tanzania (n=2 studies), Cape Verde (n=1 study), Libya (n=1 study), Namibia (n=1 study), Senegal (n=1 study) and Sudan (n=1 study). Majority of the included studies were undertaken in Northern Africa (n=65 studies, 50%), Southern Africa (n=35 studies, 26.9%) and West Africa (n=18 studies, 13.9%). Different rates of resistance to antibiotics were reported in different countries in Africa (Tables 2-5; Supplementary data 1).

A meta-analysis of published literature confirmed the presence of a high mean rate of drug resistance in GPB isolated from humans (62.0%, 95% CI: 54.7 – 69.3%), animals (68.2%, 95% CI: 58.0 -78.4%) and the environment (84.6%, 95% CI: 69.9 – 99.3%) (*P-value*<0.0001) in Africa, albeit many studies that did not address the molecular mechanisms of resistance in GPB were excluded. Obviously, the mean rate of resistance would have been higher had all research articles using only phenotypic methods to describe ABR in GPB been included (Supplementary data 1). Interestingly, although a lesser number of GPB were isolated from environmental sources, they expressed higher ABR than those from humans and animals; hence, the higher mean resistance rate of 84.6%. This also underscores the fact that there is increasing ABR genes in the environment, obviously due to antibiotic pollution from human activity. Evidently, ABR is high among GPB in certain regions in Africa (Figures 3a & 3b) (Table 5) and underpins the need to up the ante against this menace through increased molecular surveillance research, education of clinical microbiologists on ABR, and antibiotic stewardship.

Studies describing detailed molecular mechanisms of GPB resistance and molecular epidemiology in Africa are few, making it difficult to paint a vivid comprehensive picture of ABR in Africa. However, this review shows that *S. aureus* ST5, *E. faecium* ST18, ST80 and ST910, *E. faecalis* and *S. agalactiae* harbouring *mec*A*, tet* and *erm* genes, were commonly found in humans, animals and the environment, particularly in Northern, Western, and Southern Africa. Thus, careful use of β-lactams, tetracyclines, and macrolides is warranted to prevent further selection and dissemination of these resistance genes and resistant clones. Furthermore, it will be prudent for countries within these regions to review their recommended antibiotic regimens, guidelines/protocols for infections caused by these species.

*erm*(B) and *tet*(M) were found in *S. aureus, Enterococcus spp. and Streptococcus spp.,* with *erm(B), tet*(M) and *vanA* genes being mobilized by Tn*916* and IS*16,* indicating horizontal transfer within same clones, different clones and species. The discovery of same clones and resistance genes in specimens from humans, animals and the environment suggest a possible transmission of these clones between humans, animals and the environment, corroborating the need for a One Health approach to infection control and management of antibiotic-resistant infections. Further molecular epidemiological surveillance in the above-mentioned states is crucial to forestall further spread of these resistant pathogenic clones both within their borders and from their borders to other countries.

### Resistance rates per countries and MDR GPB species

High mean resistance rates were reported in Sudan (98.5%), South Africa (82.7%) Nigeria (71.2%), Egypt (70.5%), Angola (66.2%), Tunisia (66.8%), Ghana (65.1%), Algeria (62.2%) etc. (Table 5). Cross-contamination of multi-drug resistant bacteria between patients and the environment accounted for the high rate of resistance in Algeria ^45-49^. The high rate of ABR in Tunisia was attributed to cross contamination between hospital patients and hospital environment, immune deficiency ^50^, overconsumption of antibiotics, heavy consumption of sheep meat, which is a reservoir of MRSA, and high consumptions of antibiotics in animal feed ^51,52^. In Egypt, inappropriate antibiotic prescription practices ^29^, inadequate hygienic handling and processing of food ^12^, and close contact with pet dogs accounted for the high resistance ^53^. The high rate of drug resistance in Nigeria has been attributed to the exchange of resistance genes between farm animals or their products and man ^54,55^, existence of MRSA in clinical and community settings ^56^, uncontrolled usage of antibiotics ^57^ and the presence of efflux pumps in coa
gulase - negative staphylococcus strains ^58^. Expansion of resistant clones ^59^, variability of hospital acquired MRSA clones ^60^, consumption of unpasteurized milk or inefficient thermal processing of milk ^21^, shedding of resistant clones from animals to the environment and heavy consumption of antibiotics to treat TB due to high HIV burden ^61^, were incriminated for the high-level resistance in South Africa.

*Staphylococcus spp. (S. aureus, S. haemolyticus* and *S. saprophyticus); Streptococcus spp. (S. pyogenes* and *S. agalactia),* and *Enterococcus spp. (E. faecium,* E *faecalis, E. hirae, E. durans,* and *E. gallinarum*) were the antibiotic-resistant GPB widely distributed in Northern, Southern, Western and Central Africa. The high number of *tet*(M/L/K), *erm*(A/B/C), *aph(3’)-lll* and *vanA/B/C* in *Staphylococcus spp., Enterococcus spp.,* and *Streptococcus spp.* reported in Tunisia, South Africa, Nigeria, Algeria and Egypt accounted for the high rate of resistance to tetracycline, erythromycin, kanamycin and vancomycin (Figure 3a). Such resistant GPB are known to compromise the safety of invasive medical procedures such as organ transplants, orthopedic surgery, and cancer treatment. In addition, infections such as sepsis, endocarditis, deep wound infections, pneumonia, meningitis and urinary tract infections caused by these resistant pathogens are becoming increasingly fatal due to limited treatment options ^62,63^. The abuse of antibiotics as growth promoters, prophylaxis, and metaphylaxis in food animals in these countries have been implicated in the selection of resistant bacteria that can pass on to humans through food consumption, direct contact with animals and the environment, as well as trade of animals and food products between countries ^64^.

Approximately 26, 385 GPB were isolated from humans (n=83 studies), animals (n=32 studies) and the environment (n=14 studies) (Tables 1-4), with mean rates of ABR varying from 14.2% to 98.5% across the 20 included countries (Tables 2-5). The antibiotics to which the isolates were most resistant to were penicillin (n=4 224 isolates, 76.2%), erythromycin (n=3 552 isolates, 62.6%), ampicillin (n=1 507 isolates, 53.9%), sulfamethoxazole/trimethoprim (n=2 261 isolates, 46.0%), tetracycline (n=3 054 isolates, 42.1%), vancomycin (n=1 281 isolates, 41.2%), streptomycin (n=1 198 isolates, 37.0%), rifampicin (n=2 645 isolates, 33.1%), ciprofloxacin (n=1 394 isolates, 30.5%), clindamycin (n=1 256 isolates, 29.9), and gentamicin (n=1 502 isolates, 27.3%) (*p*-value <0.0001) (Tables 2-4 & 6).Countries with high number of studies such as Tunisia, South Africa, Egypt and Nigeria recorded high number of ABR (Table 5) and high number of *mecA, erm*(B), *tet*(M), *drfG* and *vanB* resistance genes (Figure 3a). Vancomycin resistance was reported in seven studies each for animals and the environment, and 12 studies in Humans. Vancomycin-resistant *Enterococcus spp.* (n=102 isolates) and vancomycin-resistant *Staphylococcus spp.* (n=258 isolates) were reported in humans, animals and the environment (Tables 2-4; Figures 2). Vancomycin-resistant *Staphylococcus aureus* (VRSA) was reported in animals (n=238 isolates), the environment (n=15 isolates) and humans (n=5 isolates). A similar situation occurred with vancomycin-resistant *E. faecium,* which was isolated from the environment (n=306 isolates), animals (n= 671 isolates) and humans (n=26 isolates) (Supplementary data 1).

Antibiotic-resistant *S. aureus* ST5, *E. faecium* (ST18, ST80 and ST910) and *E. faecalis* harbouring *mecA, erm*(B), *erm*(C), *tet*(M), *tet*(K), *tet*(L) and *vanB* were isolated from humans, animals and the environment, albeit in higher proportion in humans and animals than the environment (Tables 2-4). For instance, Farhat et al. (2014) ^46^, van Rensburg *et al.* (20 1 2) ^59^ and De Boeck et al. (2015) ^65^ in Algeria, South Africa and Democratic Republic of Congo respectively, reported on resistant *S. aureus* ST5 in humans whilst Fall et al. (2012) ^66^ reported on the same clone (S. *aureus* ST5) in pigs from Senegal. Further, Mariem et al. (2013) ^24^ isolated the same clone (S. *aureus* ST5) from the environment in Tunisia, suggesting that this clone is widely distributed in Africa in humans, animals and environment. It is currently not clear whether this clone first emerged from humans, animals or the environment, but its presence in all three spheres shows the possibility of resistant species and clones being disseminated between animals, humans and the environment. Notably, *S. aureus* ST5 is among the frequently reported clones in Asia ^67^ and recent evidence suggest that it has spread from hospitals into communities, resulting in community-acquired MRSA ^68^.

Similarly, Lochan et al. (2016) ^30^ in South Africa, Dziri et al. (2016) ^20^ and Elhani et al. (2014) ^69^ in Tunisia isolated resistant *E. faecium* ST80 from humans. For the first time, E. faecium ST80 was isolated from environmental samples in a hospital in Tunisia by Elhani et al. (2013) ^69^ and Dziri et al. (2016) ^70^. Transmission of this resistant clone to animals is possible, although not yet reported. This implies that these resistant species and clones are circulating between humans and the environment, underpinning the broad host range and transmissibility of these strains between humans and the environment.

*mecA* was the predominant resistance gene, which corresponded with the higher penicillin resistance recorded (Figure 2aii). MRSA strains were the most commonly isolated strains (≥ 2,350) ^71-74^. This is consistent with the global report of increasing prevalence of MRSA ^75,76^. MRSA harbours the *mecA* gene, which is carried by the *SCCmec* MGE, and mediates resistance to multiple β-lactam antibiotics^77^. From this review, MRSA showed resistance to eleven different antibiotic classes: aminoglycosides (gentamicin, tobramycin), β-lactams (penicillin, ampicillin, oxacillin, cefoxitin), fluoroquinolones (ciprofloxacin, levofloxacin, ofloxacin), glycopeptides (vancomycin), lincosamide (clindamycin), macrolides (erythromycin), phenicols (chloramphenicol), rifamycins (rifampicin), streptogramins (pristinamycin), sulfonamides (trimethoprim/sulfamethoxazole), and tetracyclines (tetracycline). MRSA is thus a worrying public health threat as some strains have evolved resistance to almost all licensed drugs (26). Vancomycin-resistant Enterococci (VREs) (≥ 594), which were reported in Northern and South Africa, also pose a serious threat to public health as they are resistant to vancomycin, a glycopeptide that is reserved for fatal or life-threatening Gram-positive infections, and other important antibiotics such as ampicillin, erythromycin, fluoroquinolones (ciprofloxacin, levofloxacin), gentamicin, rifampicin, streptomycin, trimethoprim/sulfamethoxazole and tetracycline. In this study, enterococcus isolates had a resistance rate of 60.1% (95%, CI=32.2 -87.9) (p-value = 0.0005) to vancomycin (Table 6). Multidrug resistance in VREs increases VRE-associated mortality rates, which is likely to increase to 75% compared with 45% from susceptible strains ^13,80^. As well, evolution of macrolide resistance (42.0%, 95% CI: 12.02 – 72.1) (p-value = 0.0129) in drug-resistant streptococci is limiting treatment options and resulting in high mortalities ^81-83^. In this study, MRSA, VRE and drug-resistant streptococci remain major public health threats, calling for measures to contain ABR. Novel antibiotics such as linezolid, synercid, and daptomycin should be used empirically whilst awaiting susceptibility results. The empirical therapy can be changed or maintained based on the susceptibility report ^84^.

### Resistance rates of species per animals, humans and the environment

The rates of ABR in isolates recovered from the environment was highest, followed by isolates from animal sources. Among environmental isolates, 91.2% (95%, CI=78.8–103.6) were resistant to penicillin, 82% (95%, CI=40.6–123.4) were resistant to sulfamethoxazole/trimethoprim, 68.5% (95%, CI=24.1–100) were resistant to ampicillin, 60.8% (95%, CI=25.0–96.6) were resistant to vancomycin, 56.9% (95%, CI=-40.7–73.2) were resistant to erythromycin, 54.5% (95%, CI=29.49–79.5) were resistant to ciprofloxacin, and 51.3% (95%, CI=21.3–100) were resistant to clindamycin (Table 6). Among animal isolates, 71.8% (95%, CI=54.9–88.73) were resistant to penicillin, 58.9% (95%, CI=36.1–81.7) were resistant to clindamycin, 58.5% (95%, CI=37.6 –79.4) were resistant to ampicillin, 49.6% (95%,CI=30.1–69.1) were resistant to trimethoprim/sulfamethoxazole, 42.3% (95% CI=17.7–67.0) were resistant to vancomycin, 47.6% (95% CI=34.0–61.2) were resistant to erythromycin, and 38.8% (95% CI=21.3–56.3)(p-value = 0.15) were ciprofloxacin resistant (Table 6; Supplementary file 1).

The rates of resistance were much lower in humans for most of the antibiotics used (Tables 2-4). Among the various species, *Enterococcus spp.* and *Staphylococcus spp.* recorded high rates of resistance for most antibiotics (Figure 3b). *Streptococcus spp.* reported low rates of resistance except for tetracycline to which it recorded a high rate of 55.13% (95%, CI=20.63.18–89.64) (p-value = 0.006). Resistance to vancomycin was not reported in any *Streptococcus spp.* Isolate (Table 6).

*Enterococcus spp,* mainly *E. faecium* and *E. faecalis,* recorded a resistance rate of 98.5% (95%, CI=94.5–102.6)(p-value = 0.0001) to clindamycin, 81.6% (95%, CI=52.1–110)(p-value = 0.0008) to trimethoprim/sulfamethoxazole, 64.0% (95%, CI=50.0–78.1)(p-value=0.0001) to erythromycin, 60.1% (95%, CI=32.2–87.9)(p-value = 0.0005) to vancomycin, 57.3% (95%, CI=24 -90.7)(p-value=0.0057) to penicillin, 51.7% (95%, CI=35.8–67.6)(p-value=0.0001) to tetracycline, 49.9% (95% CI=31.3–68.5)(p-value=0.0001) to ciprofloxacin, 48.9% (95% CI=20.6–77.2)(p-value=0.004) to kanamycin, 47.1% (95% CI=26.7–67.7)(p-value=0.0006) to ampicillin, 40.8% (95% CI=24.3–57.4)(p-value=0.0001) to streptomycin and 34.0% (95% CI=19.7–48.4)(p-value=0.0002) to gentamicin (Table 6).

*S. aureus* showed high resistance (79.6%) to penicillin (95% CI=69.7–89.5)(p-value = 0.0001), 67.8% to erythromycin (95% CI=11.5–147.0)(p-value = 0.0917), 55.5% to ampicillin (95% CI=44.50–88.5)(p-value = 0.0001), 39.3% to trimethoprim/sulfamethoxazole (95% CI=39.3–47.8)(p-value = 0.0001), 36.9% to tetracycline (95% CI=29.3–44.5(p-value = 0.0001), 35.8 to streptomycin (95% CI=14.7–57.0)(p-value = 0.004), 33.6% to rifampicin (95% CI=20.1–47.03)(p-value = 0.0001), 24.0% to clindamycin (95% CI=14.9–33.1)(p-value = 0.0001), 23.9% to ciprofloxacin (95% CI=17.6-30.2)(p-value= 0.0001), 22.7% to vancomycin (95% CI=4.3–41.2)(p-value = 0.0212) and 22.2% to vancomycin (95% CI=15.7–28.3)(p-value = 0.0001) (Table 6).

### Resistance mechanisms, clones, and MGEs

Few studies identified the clones and MGEs in the resistant isolates. Of the 130 included studies, 32 identified the clones whilst 22 described the MGEs, which were used in the statistical analysis. The most dominant gene detected in Africa, which was widespread and responsible for resistance in GPB, was *mecA* (n=3 547), followed by *erm*(B) (n=1 268), *van*C1/2/3 (n=971), *tet*(M) (n=720), *blaZ* (≥565), *dfrG* (n=422), *vanB* ≥451), *aph(3’)-IIIa* (≥170) and *aac(6’)-aph(2’*)(≥268) (p-value = 0.0011) (Fig. 2a).

Figure 2b represents MGEs per clone. *S. aureus* clones ST5, ST8, ST 80 and ST88 were highly associated with *mecA.* Resistant *S. aureus, E. faecium* and *E. faecalis* clones such as *S. aureus* ST5, and *E. faecium* clones ST18, ST80, and ST16 were widely distributed in humans, animals and the environment. Similarly, *mecA,* erm(B), erm(C), tet(M), tet(K), tet(L), *vanB, vanA, vanC* and tet(O) were reported in isolates from humans, animals and the environment (Table 1).

IS*16* and Tn916 were found with the resistance genes *erm*(B) and *tet*(M) in *E. faecium* (ST18, ST80 and ST910), *S. agalactiae* (ST612, ST616 and ST617), *E. faecalis* and *S. pyogenes (emm18, emm42, emm76* and *emm118*) isolated from humans, animals and the environment (Tables 2-4; Figure 2b). *tet*(M) was associated with Tn916 transposon in tetracycline-resistant *S. agalactiae* ^85^ and *S. pyogenes* ^81^ in humans in Tunisia. Fischer et al. (2013) also reported the association between Tn916 and tet(M) in tetracycline-resistant *S. agalactiae* in camel in Kenya ^86^. Similarly, IS *16* was found in vancomycin-resistant *E. faecium* (ST80, ST180 and ST910) in humans and the environment in Tunisia ^69,70^. Investigations into the association between MGEs and resistance genes were limited by few studies (n=22 studies) on MGEs.

From Tables 2-4, majority of the resistance genes namely, *mecA, erm*(B), *tet* (M), *vanA* etc. were responsible for drug resistance to antibiotics such as aminoglycosides (gentamicin, streptomycin, kanamycin), β-lactams (penicillins, cephalosporins), fluoroquinolones (ciprofloxacin), macrolide (erythromycin), sulfamethoxazole/trimethoprim, tetracycline and glycopeptides (vancomycin). These resistance genes were widely distributed in Northern Africa (Tunisia, Algeria, Egypt, Morocco, and Libya) and Southern Africa (South Africa and Namibia). All the three different MGEs (Tn*916*, SCC*mec* and IS*16*) were reported in Tunisia, with two being reported in Kenya (SCC*mec* and Tn*916*). IS*16* was only reported in an E. *faecium* infection in Tunisia (Figure 3) whilst *mecA* was mostly associated with SSC*mec. erm*(B) and *tet*(M) were highly associated with Tn*916* and IS*16*.

In Africa, different studies have reported *SCCmec-borne mecA* in *S. aureus* in humans, animals and the environment ^23,47,60,66,87^ besides the discovery of IS*16* and Tn*916* in the environment of *erm(B)* and *tet*(M) genes in Enterococcus and Streptococcus. These reports show that MGEs are mediating the dissemination of these (and possibly other) resistance genes across different GPB clones and species. MGEs-mediated mobilization of various resistance genes in different GPB clones and species in humans, animals and the environment (Tables 1-4; Figure 2b) calls for prompt measures to contain ABR as the situation may worsen if additional resistance genes are acquired by the MGEs. Resistance genes on MGEs can be horizontally transferred to susceptible cells or vertically transferred to daughter clones ^37,88,89^, which can easily spread these resistance genes to susceptible pathogens. The higher number of resistant Gram-positive cocci and mean resistance rate in Tunisia may be due to the presence of these three MGEs in this region ^69,70,81,90^

### Molecular epidemiology of antibiotic-resistant GPB

### North Africa: Algeria, Egypt, Morocco, Tunisia, Libya

#### Algeria

*S. aureus* was recovered from two different studies in Algeria. In assessing the nasal carriage of *S. aureus* in patients with medical conditions including pneumonia, urinary tract infections, osteoarthritis, heart diseases, diabetes and chronic kidney disease, Djoudi *et al.* (2014) isolated MRSA ^46^. They also found nasal carriage of *S. aureus* to be significantly associated with cancer and previous hospitalization of patients with kidney failure due to immunological suppression and hemodialysis. The nine MRSA isolates, i.e. ST80 (n=4), ST5 (n=2), ST22 (n=2) and ST535 (n=1), harboured *mecA* and were resistant to tobramycin (n=6), gentamicin (n=1), trimethoprim/sulfamethoxazole (n=2), tetracycline (n=3) and erythromycin (n=1). MRSA ST80 is a well-known and frequent etiological agent of infections in North Africa and Middle-East countries^91,92^. Typing of 64 MRSA isolated from human pus (n=47), venous catheters (n=7), tracheal aspirates (n=4), punction fluids (n=3), blood (n=2) and urine (n=1) in 64 Algerian patients revealed that 50 were hospital acquired (HA-MRSA) and 14 community acquired (CA-MRSA), which were all resistant to cefoxitin and oxacillin ^47^. *mecA,* mobilized by *SCCmec,* was the only detected mechanism of resistance.

#### Egypt

MRSA have been respectively isolated in five animal-based and two human-based studies in Egypt between 2011 to 2017. Hashem et.al (2013) isolated 94 *S. aureus* strains from blood and wounds in which 45 were MRSA while 25 were fluoroquinolone-resistant ^29^. Mutations such as C**2402**T, T**2409**C, T**2460**G, T**1497**C, and A**1578**G in gyrase enzymes, which leads to fluoroquinolones’ target-site alterations, were implicated in resistance to fluoroquinolones (ciprofloxacin, levofloxacin, ofloxacin). The high rate of fluoroquinolone resistance (55.56%) among MRSA infections is rather concerning as patients unable to tolerate vancomycin are treated with other antibiotics such as fluoroquinolones. Vancomycin is often reserved as a last-resort therapy for MRSA infections due to their high resistance to several antibiotics.

Multidrug resistance to drugs such as gentamicin, ampicillin, amoxicillin, cefepime, tetracycline and chloramphenicol in MRSA is mediated by diverse resistance mechanisms including impermeability effects and efflux pumps. Unrestricted access to antibiotics and inappropriate prescriptions were responsible for the high rates of drug resistance in this study ^29^. In a similar study, MRSA was isolated from patients suffering from surgical wound infections, diabetic foot, abscess and burns. Although *mecA* was the only mechanism of resistance, the isolates were multiple-resistant to several antibiotics belonging to the β-lactams, aminoglycosides, fluoroquinolones, macrolides, lincosamides, tetracyclines and glycopeptides, indicating other mechanisms of resistance ^93^. It therefore implies that administration of such antibiotics will not relieve patients from *S. aureus* infections. The high rate of *S. aureus* isolation confirms it to be the most prevalent Gram-positive pathogen isolated from soft tissue and wound infections.

Al-Ashmawy *et. al.* detected a high rate of MRSA (53%) in milk and dairy products believed to originate from human contamination rather than contamination from animals. Besides being resistant to β-lactams and other antibiotics, thirty-six of the isolates were resistant to vancomycin known to be effective in treating MRSA infections ^12^, making milk and dairy products a significant source of multidrug-resistant and toxigenic *S. aureus* infections. The occurrence of MRSA in pets such as dogs admitted in a veterinary clinic ^53^ may confirm a possible route in the community transmission of this pathogen, which is emerging as a veterinary pathogen of public health importance.

In 2017, Osman and colleagues detected *Staphylococcus spp.* in imported beef meat. Sixteen of these isolates were MDR and showed resistance to different groups of antibiotics due to resistance mechanisms such as *mecA,* and mutations in *gyrA* and *gyrB.* Indeed, MRSA has made methicillin and other β-lactams antibiotics clinically useless as a result of their high MDR ^94^. Imported meat acts as a transmission vector for MRSA and is worrisome as *Staphylococcus spp.* are among the most common foodborne pathogens causing food poisoning outbreaks worldwide. Of 133 *S. aureus* recovered from animal origin, more than 70% were MDR and 30 were MRSA, exhibiting high resistance to clindamycin, co-trimoxazole, tetracycline, oxacillin, cefoxitin, ceftriaxone and erythromycin; four of the isolates were resistant to vancomycin ^23^. The isolates showed the maximum sensitivity to imipenem, chloramphenicol and rifamycin, which is consistent with similar reports in China and Pakistan ^95,96^, indicating their effectiveness in treating *S. aureus* infections.

MRSA was isolated from chicken products mainly due to poor hygienic handling processes, posing a risk to public health in 2016. The mean *S. aureus* count in the chicken products were beyond the permissible limits of the Egyptian organization for Standardization and Quality Control (EOSQC 2005), coupled with resistance to different antibiotics classes; thus, retail chicken products could constitute a high health risk to human consumers ^28^

#### Morocco

In a study to assess *S. aureus* carriage among end-stage renal diseases patients undergoing hemodialysis, 42.9% *were* carriers, of which only one was MRSA. The methicillin-susceptible *S. aureus* (MSSA) was resistant to many of the local antibiotics, thus limiting the successful treatment of MSSA infections. Moreover 81.8% of the MSSA were penicillin-resistant. The male gender and age 30 or below were identified as risk factors of *S. aureus* nasal carriage *(P-value* < 0.001) ^27^. Periodic monitoring of patients with hemodialysis is crucial as they are at increased risk of *S. aureus* infection due to periodic hospitalization, immunosuppression and high invasive vascular interventions.

#### Tunisia

Resistant *S. aureus* was isolated from the environment, animals and humans between 2011 to 2017. Ben Said, et al. recovered 12 MSSA from wastewater samples that were resistant to penicillin (n=12 isolates), erythromycin (n=7 isolates), tetracycline (n=1 isolate) and clindamycin (n=1 isolate) due to the presence of *blaZ* (n=7), *msr*(A) (n= 7) and *tet*(K)(n=1). These resistant strains were of ST3245(n=7) and ST15(n=1) ^18^, which have been also reported in animals and humans. In an investigation to evaluate the prevalence of coagulase-negative Staphylococcus (CoNS) in the hospital environment, MDR *S. haemolyticus* and *S. saprophyticus* were the most dominant. Methicillin resistance was detected in *S. haemolyticus*, *S. epidermidis and S. saprophyticus*. These isolates were resistant to erythromycin, tetracycline, gentamicin, kanamycin, tobramycin and streptomycin due to the presence of *msrA* (32), *erm*(C) (8), *tet*(K) and *tet*(M), *aac(6’)-Ie-aph(2”)-Ia (16),), aph(3’)-IIIa(19), ant(4’)-Ia (n=14)* and *ant(6 ‘)-Ia (3)* ^97^. The high prevalence of MDR *Staphyloccoci spp.* isolates may result from transmission between the staff, patients and the environment. Strict infection controls are needed as infections caused by CoNS are common cause of death, particularly in low-birth-weight children, and are opportunistic infections in immunocompromised patients ^98^.

Moreover, nasal swab from sheep detected five MRSA (*mecA*=5), which were all of ST153 and carried *blaZ, ant(6)-Ia, aph(30)-IIIa, erm*(C), *tet*(K), and *fusB* genes that respectively encoded resistance to penicillin, streptomycin, kanamycin, erythromycin, tetracycline and fusidic acid. This study shows that the nares of healthy sheep could act as reservoirs of MRSA ^99^.

Between 2011 to 2012, 99 MRSA strains were detected from nasal swabs, blood, catheter, wounds, pleural puncture and abscess, among which 39 were tetracycline resistant. These isolates were resistant to aminoglycosides, fluoroquinolones, macrolides and lincosamide, with mechanisms of resistance including *mecA* (n=24) *tet(*K) (n=6) *tet*(L) (n=1) and/or *tet*(M) (n=18), *erm*(A)(n=14), *aph(2’)-acc(6’)* (n=13). Identified drug-resistant strains included ST247 (n=12), ST239 (n=6), ST728 (n=2), ST241 (n=1), ST398 (n=1), ST5 (n=1) and ST641 (n=1) ^50^. For the first time, clonal lineage ST398, which has been reported in pigs from several studies in USA, South America, Asia and Canada ^100-103^, was found in human MRSA isolates in Africa in a nasal swab of a 74-year old patient.

Additionally, 69 MRSA strains were isolated from hospital-acquired and community-acquired infections. Although *mecA* (n=59) was the only mechanism of resistance identified, the isolates were resistant to aminoglycosides, tetracycline, fluoroquinolones, macrolides and rifampicin. The resistant clones were ST1 (n=2), ST5 (n=5), ST22 (n=1), ST80 (n=41), ST97 (n=2), ST153 (n=2), ST239 (n=4), ST241 (n=3), ST247 (n=3), ST256 (n=1), ST1819 (n=3) and ST1440 (n=1) ^24^.

Mezghani Maalej and colleagues (2012) isolated five pristinamycin-resistant *S. aureus* strains from patients with skin infections. These isolates were MDR (Table 2), being the first detection of resistance to streptogramins due to *vat*(B) and *vga*(B) resistance genes ^104^, which emerged due to selective pressure from the use of pristinamycin. Thirty-six methicillin-resistant *S. haemolyticus* (MRSHae) were isolated from neutropenic patients (suffering from febrile neutropenia) with hematological cancer between 2002 and 2004. These MDR isolates carried SCC*mec*-borne *mecA* (Table 2) ^105^, which agrees with a report on *S. haemolyticus*’MDR capacity, particularly in immunocompromised patients^106,107,^

#### Libya

Due to the high risk of MRSA colonization developing into infections in children, nasal samples were collected from children inpatients, their mothers, healthcare workers and outpatients’ workers, which yielded a MRSA nasal carriage rate of 8.3%, 11%, 12.3% and 2.2% respectively in Libya ^108^. Thus, nasal carriage of MRSA is common in inpatients children, their mothers and health workers in Libya and could be a source of MRSA infections.

### West Africa: Ghana, Nigeria, Senegal

#### Ghana

Among 308 staphylococcus isolates collected across Northern, Central and Southern Ghana in 2013, low prevalence of antibiotic resistance was reported except for penicillin (97%), tetracycline (42%) and erythromycin (6%) ^109^. Moreover, *mecA* was detected in only nine isolates, implying the presence of other β-lactam resistance mechanisms. The MRSA clones included ST8 (n=1), ST72 (n=1), ST88 (n=2), ST239 (n=1), ST250 (n=2), ST789 (n=1), and ST2021 (n=1). In a similar study that characterized 30 MRSA isolates resistant to tetracycline, fluoroquinolones and macrolides, *tet*(M) (n=13), *tet*(K) (n=10), *aphA3* (n=7), *aacA-aphD* (n=5) *and erm*(C) (n=4) were detected. Similar and different resistant clones, viz. ST88 (n=8), ST8 (n=5), and ST247 (n=4) were detected ^110^, indicating high MRSA clonal diversity in Ghana. These studies show a high rate of resistance to non-β lactams that further complicate MRSA treatment. Furthermore, the isolation of USA300 and other epidemic multidrug-resistant MRSA clones calls for MRSA surveillance and adequate control measures.

#### Nigeria

Five different studies reported drug-resistant *S. aureus* from several human anatomical sites such as throat swabs, soft skin and tissue infection, urinary tract and respiratory infections, wound, vagina, otitis, conjunctivitis, septicemia and bronchitis. Of a total ≥602 isolates, ≥433 were resistant to several antibiotic classes (Table 1). Of note, 429 of the ≥433 drug-resistant isolates were all resistant to cotrimoxazole or trimethoprim/sulfamethoxazole (SXT). Mechanisms of resistance included *mecA* (≥54), *blaZ* (n=284), *dfrA* (≥5) and *dfrG* (≥152). *S. aureus-*resistant clones ST8, ST14, ST37, ST39, ST88, ST152, ST241, and ST772 were present. Colonized persons, including immune-compromised individuals, facilitated the spread of *S. aureus* and MRSA ST8 identified as ubiquitous in various geographic areas of Nigeria. High utilization of co-trimoxazole or SXT because of low cost and easy obtainability through lenient medication regulations were implicated for the high resistance ^56^. Besides *S. aureus, S. haemolyticus* was the major species isolated, and is considered as the second most detected and clinically important *Staphylococci spp.,* particularly in immunocompromised patients ^111^. All the *S. haemolyticus* isolates detected were resistant to at least three antibiotics classes (Tables 2-4) ^112^.

Moreover, O. Ayepola *et al.* (2015) reported a higher rate of 20.8% *S. aureus* from UTIs than the reported ranges in Africa (6.3-13.9%), and far exceed the rate reported from Europe and Brazil (1.1%) ^113^. None of the isolates exhibited resistance to vancomycin, linezolid, daptomycin and mupirocin; indicating their usefulness in treating *S. aureus* infections. Co-trimoxazole, which was previously clinically valuable in treating MRSA infections, demonstrated the highest level of resistance, hence it’s not recommendable^56,57,90,112^. In a study to examine the genetic mechanism(s) of resistance in CoNS in faecal samples, all the 53 islolated CoNS were Penicillin V-resistant and between three to 19 exhibited multidrug resistance (Table 2); *mecA* (n=15), *erm*(C), *tet*(M) (n=4) and *tet*(K) (n=6) were identified ^112^. CoNS isolates from faeces carrying tetracycline, macrolides and aminoglycosides resistance genes may transfer them inter- and intra-species, disseminating MDR in Staphylococcus.

#### Senegal

A low prevalence of MRSA (10.5%) was reported in Senegalese pigs compared to those reported in developed countries. This might be due to a lesser veterinary antibiotic use as growth promoters and/or for therapy. However, all the isolates were resistant to penicillin, 27 were resistant to co-trimoxazole and 16 were resistant to tetracycline ^66^. Five of the MRSA were of ST5 ^66^, evincing the spread of this clone in animals, humans ^46,59^, and the environment ^24^; the importance of this clone as a cause of human infections is well-established ^68^.

#### Cape verde

In Cape Verde, a low prevalence of 5.6% (6/107) MRSA nasal carriage was documented in 2015. The predominant MRSA clones was ST5 (n=3), ST8 (n=1) and ST88 (n=2). These isolates showed significant level of resistance to erythromycin (ERY), sulphamethoxazole-trimethoprim (SXT) and penicillin G (PEN) ^114^.

### Central Africa: Gabon, D.R. Congo

#### Gabon

In Gabon, *S. aureus* isolated from colonized persons, blood, as well as soft and skin tissue infections resulted in 49% (104/212) resistance to trimethoprim: *dfrA* (n=1), *dfrG* (n=100), *dfrK+G* (n=1), *dfrB* (n=2), and *mecA* (n=1) were detected in the isolates ^55^. Thus, *dfrG* is obviously the most abundant and common trimethoprim resistance mechanism in Africa, refuting *dfrB* mutation as the main mechanism of resistance to trimethoprim ^115-117^.

#### D.R. Congo (DRC)

A total of 215 (79.3%) drug-resistant *S. aureus* isolates were collected between 2015 to 2017 from nasal swab and bloodstream infections in the D. R. Congo; 70 isolates were MRSA. Other major resistance genes mediating resistance to trimethoprim/sulfamethoxazole, aminoglycoside, macrolides, tetracycline, penicillin, and chloramphenicol were *dfrG* (≥120), *tet*(K) (≥98), and *femA* (≥98). MRSA showed high-level resistance to β-lactams, aminoglycoside, macrolides and tetracycline. The pathogen caused severe infections such as pneumonia, meningitis, complicated urinary tract infections, gynaecological infections and peritonitis. *S. aureus* ST8 (≥47) was the dominant clone, followed by ST152 (≥17), ST5 (≥2) and ST88 (≥2). In DRC, MRSA ST8 outnumbers the African MRSA clone ST88, which is dominant in Africa. The high-level oxacillin resistance in DRC was associated with a mutation in *femA* (Y**195**F) whist high-level trimethoprim resistance was due to the detection of *dfrG*, which is consistent with trimethoprim resistance in Africa and Asia. In Africa, SXT or cotrimoxazole is frequently administered as prophylactic to immuno-suppressed patients such as HIV/AIDS patients to prevent opportunistic infections such as *Pneumocystis carinii* pneumonia, toxoplasmosis and bacterial pneumonia ^118^ Hence, prophylactic use of SXT in HIV patients may impact resistance. Additionally, there was high-level MDR among MRSA, which is a great concern as microbiological laboratories/facilities and second-line antibiotics are rare in DRC. Moreover, the detection of nasal carriage among healthcare workers’ demands strict infection controls and surveillance ^65,119,120^.

### East Africa: Kenya, Tanzania

#### Kenya

In contrast to earlier studies done in Kenya, Omuse and colleagues (2016) detected a wide genetic diversity of MRSA and well-established epidemic MRSA clones among clinical isolates. MRSA clonal complexes 5, 22 and 30, implicated in several outbreaks were described. These clones included ST5 (n=1 isolates), ST8 (n=2 isolates), ST22 (n=4 isolates), ST88 (n=1 isolates), ST241 (n=12 isolates), ST239 (n=2 isolates) and ST789 (n=1 isolates). Approximately 41% of the MRSA in the study were MDR (Table 2), showing resistance to clindamycin, erythromycin and SXT ^87^. Detection of these clones in referral hospitals in Kenya calls for implementation of strict infection control measures to reduce the high morbidities and mortalities associated with HA-MRSA infections.

#### Tanzania

In a study to investigate the molecular epidemiology of trimethoprim resistance in MSSA causing skin and soft tissues infections, *dfrG* was detected in all 32-trimethoprim resistant isolates. Other reported trimethoprim resistance mechanisms such as *dfrA*, *dfrB* and *dfrK* were missing, confirming *dfrG* as the main trimethoprim resistance mechanism in Sub-Sahara Africa ^55^.

#### Uganda

A MRSA carriage of 56.1% (23/41) was detected in milk from pastoral communities in Uganda, exactly 70% of which were tetracycline-resistant. MRSA clones ST97 and ST1 were identified. Furthermore, over 90% of the isolates carried genes encoding enterotoxin that causes food-borne diseases. The weak veterinary delivery system and the high dependency on animals and animal products for food in Uganda was implicated for the high prevalence of MRSA ^121^.

*S. aureus* isolates, including 24 MRSA and 40 MSSA, were isolated from patients with surgical site infections (SSI). The MRSA isolates were MDR (including resistance to oxacillin, gentamicin, ciprofloxacin and chloramphenicol) compared to the MSSA. Inducible clindamycin resistance was found in 17.2% of the isolates, mostly in MRSA. In a multivariate analysis, inducible clindamycin resistance and cancer were identified as independent predictors of MRSA-SSI ^122^.

### Southern Africa: Angola, Malawi, Mozambique, Namibia, South Africa

#### Angola

Conceicão et al (2014) reported a nasal *S. aureus* carriage of 23.7% (n=128 isolates), out of which 58.1% (n=77 isolates) were MRSA. Fifty-seven of the MRSA clones were of ST5, followed by ST88 (n=9), ST8 (n=5) and ST72 (n=3). This study represents the first description of the spread of MRSA ST5 in Africa. All the 77 MRSA strains were resistant to SXT, cefoxitin (FOX) and PEN ^123^. In a study to identify oxacillin-susceptible *mecA*-positive *S. aureus* (OS-MRSA) for the first time in Africa, a prevalence of 17.7% was detected among healthy healthcare workers in Angola and São Tome′ & Principe, making them potential OS-MRSA reservoirs ^124^. OS-MRSA have been reported worldwide in humans, animals and food animals ^125-128^. The OS-MRSA isolates expressed MDR (Table 2) and belonged to ST88 (n=15 isolates) and ST8 (n=9 isolates). In sub-Saharan Africa, the identification of clinically important *S. aureus* is heavily based on phenotypic agar-screening and oxacillin disc-diffusion methods.

#### Mozambique

The prevalence of HA-MRSA and CA-MRSA in Mozambique was found to be 15.1% and 1%, respectively. MRSA showed high-level resistance to penicillin, cefoxitin, gentamicin, ciprofloxacin, erythromycin, SXT, chloramphenicol and tetracycline, compared to MSSA. Additionally, inducible macrolide–lincosamide–streptogramin B (MLSB) resistance was 41.7% and 10.7% in hospital-acquired *S. aureus* (HA-SA) and community-acquired *S. aureus* (CA-SA) isolates respectively ^129^, further limiting therapeutic options for *S. aureus* infections. This study, which is the first to detect the emergence of HA-MRSA within post-operative abdominal wounds and burn wounds in Mozambique, reported that patients with infected burn wounds had a significantly longer hospitalization than patients with post-operated abdominal wounds. Efforts to prevent the transmission of MDR HA-SA, such as education on proper hand-washing techniques, are urgently needed.

#### Namibia

The dominant resistance gene mediating trimethoprim resistance in MRSA and MSSA in Namibia was *dfrG.* This is similar to reports in other Africa countries ^55^. Moreover, *dfrG* was frequently detected in *S. aureus* from SSTIs in travelers returning from other African countries, suggesting that *dfrG* can be transmitted into populations with low antifolate resistance such as North America and Europe ^130,131^.

#### South Africa

Thirty MDR *S. aureus* were recovered between April 2015 to April 2016 from ten beaches in the Eastern Cape Province, South Africa (Table 2). Notably, the isolates harbored *mecA, femA, rpoB, blaZ, erm*(B) and *tet*(M) ^11^, making marine environments and public beaches potential depositaries of MDR *S. aureus* that can be transmitted to animals and humans. Further, the 50% resistance to vancomycin recorded is concerning to global health due to its role as a last-resort antibiotic for treating MRSA infections.

*S. aureus* was detected in raw and pasteurized milk at an isolation rate of 75% and 29% respectively, due to inefficient thermal processing and post-process contamination. A high proportion (60%-100%) of these isolates showed resistance to aminoglycosides, β-lactams, vancomycin, tetracycline and erythromycin, albeit only 19 *mecA* genes were present ^21^. Evidently, raw and pasteurized milk can harbour MDR *S. aureus,* exposing consumers to colonization and/or infections. Again, *Staphylococcus spp.,* including *S.aureus, S. haemolyticus*, *S. xylosus* and *S. capitis* were isolated from healthy pigs and cattle, of which between 75 to 100% were resistant to penicillin G, tetracycline, sulfamethoxazole and nalidixic acids, due to their use as growth promoters; *mecA* and *mphC* were identified. Additionally, 12% of the isolates were resistant to vancomycin and erythromycin, evincing the important role of animals in the dissemination of resistance determinants and the importance of commensals to public health ^61^.

Van Rensburg et al. ^59^ detected 43.4% (1432/3298 isolates) and 3.1% (328/10448 isolates) rifampicin resistance rate among MRSA and MSSA respectively. Similar studies in South Africa have also reported of high rifampicin resistance in MRSA^132, 133^, obviously due to frequent use of rifampicin among tuberculosis patients, who are highly prevalent in South Africa. MRSA ST5 and ST612 were detected while H**481**Y/N and I**527**M mutations in *rpoB* were associated with high-level rifampicin resistance, similar to reports in Italy ^134^. Additionally, novel H**481**N, I**527**M, K**579**R mutations were also detected.

Three studies reported a prevalence of 29.1% ^135^, 45.44% ^60^ and 100% ^136^ MRSA recovered from humans, expressing resistance to macrolides, tetracycline, aminoglycoside, cotrimoxazole and rifampicin. MRSA ST612, ST239, ST36 and ST5 were the dominant strains similar to other findings in Australia and Europe^137^. The study showed that *S. aureus* bacteremia is common and account for high mortality in South Africa. For instance, in a study by Perovic et al., ^135^ 202 patients died from *S. aureus* bacteremia infections, with HIV patients being more likely to acquire HA-MRSA. The isolates were however susceptible to glycopeptides, fluoroquinolones, linezoid, tigecycline, fosfomycin and fusidic acid, confirming their clinical usefulness in treating MRSA infections. In a recent study, a high prevalence and genetic diversity of multi-drug efflux (MDE) resistance genes were found in clinical *S. aureus* isolates, including 81 MRSA and 16 MSSA ^138^. *norA, norB, mepA, tet*(38), *sepA, mdeA, imrs* and *sdrM* were present in at least 86% of the isolates, predicting resistance to broad-spectrum biocides and fluoroquinolones, which is disturbing. Efforts to develop efflux pump inhibitors can mitigate such resistance mechanisms.

#### Sao Tome & Principe

MRSA prevalence of 26.9% ^139^ and 25.5% ^114^ was reported in nasal swabs in 2014 and 2015, respectively, in Sao Tome & Principe. Additionally, a high prevalence of oxacillin-susceptible *mecA*-positive *S. aureus* was reported in the same study in Sao Tome & Principe and Angola ^124^. The most dominant MRSA clone was ST8 (n=25 isolates), followed by ST5 (n=13 isolates) and ST80 (n=13 isolates). High genetic variability was found in the MSSA strains. Both MRSA and MSSA showed different levels of resistance to SXT, ERY, CIP and TET; however, all the MRSA isolates were resistant to cefoxitin.

#### Streptococcus spp. (S. pyogenes, S. pneumoniae and S. agalactiae)

Drug resistant *Streptococcus spp.* including *S. agalactiae* and *S. pyogenes* have been identified in Northern, Eastern and Southern Africa. *S. pyogenes* were reported in only humans whilst *S. agalactiae* was reported in both animals (camels) and humans with a high rate of resistance to tetracycline and erythromycin.

### North Africa: Algeria, Egypt, Morocco, Tunisia, Libya

#### Algeria

A sole study has so far detected 44 tetracycline (100%, 44/44 isolates)- and erythromycin-resistant (43.18%, 19/44 isolates) *S. agalactiae* from vaginal swabs; *tet*(M); and *erm*(B) respectively mediated this resistance. A high diversity of resistant clones viz., ST1, ST19, ST10, ST158, ST166, ST233, ST460, ST521 and ST677 were detected ^45^, which have been reported worldwide for causing life-threatening invasive diseases such a meningitis and sepsis^140,141^.

#### Egypt

Similarly, Shabayek et al. (2014) detected 98% and between 14-17% *S. agalactiae* resistance to tetracycline and macrolides respectively. *tet*(M) was detected in all the 98 tetracycline-resistant isolates whilst *erm*(B) and *erm*(A) mediated erythromycin resistance. Efflux pump genes such as *tet*(K) (n=12 isolates), *tet*(L) (n=1 isolates) and *mefA/E* (n=1 isolates) were also found ^32^, which reflects the increasing reports of *S. agalactiae* resistance to tetracycline and macrolides ^142^. This study also showed that vancomycin and fluoroquinolones are effective replacement for erythromycin and clindamycin, and for patients allergic to penicillin. Although penicillin is the antibiotic of choice for treating *S. agalactiae* infections, reports of penicillin resistance in USA and China calls for increased surveillance in Africa ^142^.

### Tunisia

#### S. agalactiae

From January 2007 to December 2009, 226 *S. agalactiae* were isolated from female genitals and gastric fluid of infected newborns. Of these, 97.35% (220/226 isolates), 40% (90/226 isolates) and 19.1% (43/226 isolates) were resistant to tetracycline, erythromycin and rifampicin respectively. Additionally, seven isolates were resistant to aminoglycoside (gentamycin and streptomycin) and chloramphenicol. *tet*(M) (n=205 isolates), encoding a ribosomal protection protein, which protect the ribosome from the action of tetracycline, was the main tetracycline resistance mechanism, and was significantly associated with *Tn916* (p-value = 0.0002). Other resistance genes including *erm*(B) (n=79 isolates) and *tet*(O) (n=50 isolates) were detected. All isolates were however susceptible to β-lactams and quinupristin-dalfopristin ^85^. Between 2005 and 2007, 160 erythromycin-resistant *S. agalactiae* were isolated from humans, with a high resistance rate of 84.3% (135/160 isolates) to the constitutive macrolides-lincosamides, streptogramines B (MLSB) ^143^.

#### S. pyogenes

Hraoui *et al.,* (2011) reported a low macrolide resistance rate (5%, 5/103) and a high tetracycline resistance rate (70%, 72/103) among human isolates, with *tet*(M), associated with Tn*916*, being responsible for tetracycline resistance ^144^. Increase tetracycline use in food animals was implicated in this instance, leading to selection and dissemination of resistance genes from animals to human. Macrolide resistance was only detected in seven isolates, which is corroborated by the findings of Ksia et al. (2010), who detected low-level macrolides resistance among Children ^145^.

### East Africa: Kenya, Tanzania Kenya

#### S. agalactiae

In the horn of Africa, camel plays a significant role in the survival of humans by providing milk, meat and transportation. In 2013, Fischer et al. detected 36% (37/92) tetracycline resistance in *S. agalactiae* isolates from camels’ wound infections and mastitis that was mainly mediated by a Tn*916*-borne *tet*(M). ST616 (n=22) was the major resistant clone, followed by ST612 and ST617 ^146^. Shifting from tetracycline to other antibiotics is evidently necessary for effective treatment outcomes in camel infections in Kenya.

### Southern Africa: Angola, Malawi, Mozambique, Namibia, South Africa South Africa

#### S. agalactiae

A *S. agalactiae* colonization rate of 30.9% was detected from vaginal and rectal swabs of pregnant women. Similar to other reports in Africa, a high rate of tetracycline (94.5%, 120/128 isolates) and macrolide (21.1%, 27/128) resistance was documented. All the isolates were however sensitive to penicillin, ampicillin, vancomycin and gentamicin. Macrolide and clindamycin resistance were associated with *erm*(B) and *mefA* genes ^147^. The study highlights the need for research on treatment options for patients allergic to penicillin due to high-level resistance in alternative drugs such as macrolides and lincosamides.

#### Enterococcus spp. (E. faecium, E. faecalis, E. hirae, E. durans, E. gallinarum)

### North Africa: Algeria, Egypt, Morocco, Tunisia, Libya Algeria

The first study to molecularly characterize *Enterococcus spp.* from urinary tract and wound infections in Algeria revealed a high rate of resistance to erythromycin (86.4%, 108/125 isolates), tetracycline (82.4, 103/125 isolates), levofloxacin (71.2%, 89/125 isolates) and gentamicin (54.4, 68/125 isolates). Only 3.2% (4/125 isolates) were VRE, confirming glycopeptides as ideal antibiotics for treating Enterococcus infections. A mortality rate of 10% was reported due to infections caused by Enterococcus. E. *faecium, E. faecalis* and *E. gallinarum* were the main Enterococcus isolated. Majority of these isolates were from females (53%). *erm*(B) (≥92) and *vanC1(*≥*4)* were the main mechanisms of resistance. A high genetic diversity among strains was seen in *E. faecium* and *E. faecalis,* with *E. faecium* ST78 being the dominant resistant strain ^148^, which is also prevalent in Asian (Japan, Taiwan, China and Korea) and European (Italy and Germany) countries ^149-151^. A novel ST317 (n=33) clone was predominant among the *E. faecalis* isolates. Rational use of antibiotics, as well as close monitoring of the epidemiology of the strains are crucial.

### Egypt

In a similar study to characterize *E. faecium* and *E. faecalis* from patients, 82% of the isolates were MDR, showing high-level resistance to aminoglycosides, β-lactams and tetracycline. *VanA* was detected in two E. *faecium* isolates, all of which were resistant to all antibiotics tested. Bioinformatic (sequence) analysis revealed that *vanA* was transmitted horizontally to *S. aureus,* showing the importance of horizontal gene transfer in ABR and subsequent management of enterococci infections such as bacteremia, endocarditis and urinary tract infections ^152^.

#### Tunisia

Antimicrobial-resistant Enterococcus was found in faeces of pet and camel, irrigation water from farm environments, food vegetables, hospital environments, animal meat and patients in Tunisia ^19,22,31,51,52,69^. High-level resistance to vancomycin, macrolides, aminoglycosides, β-lactams and tetracycline was detected in the environment, animals and humans with majority of the isolates being *E. faecium,* followed by *E. faecalis. tet*(M), *tet*(L), *erm*(B), *ant (6)-la, vanA* and *aph(3’)-llla* were the major resistance mechanisms, with IS*16* being the main MGE disseminating the resistance genes. *E. faecium* ST80, ST910 and ST16 were the dominant resistant clones in Tunisia. The studies show that meat, animals, pets, hospital environment and wastewater used for farm irrigation play a crucial role in the spread of antibiotic resistant Enterococcus.

### West Africa: Cape Verde, Ghana, Nigeria, Senegal Nigeria

*Enterococcus spp.* isolated from poultry and cattle as well as their manure demonstrated high-level resistance to tetracycline, erythromycin, gentamicin, ampicillin and streptomycin. Sixty isolates were MDR, showing resistance to three or more antimicrobials ^153^. The rate of MDR is a reflection of the substantial use of broad-spectrum antibiotics in Nigeria, raising major public health concerns as practices such as the use of untreated poultry and cattle manure for fertilizing agricultural soils, particularly vegetables, are a common practice in Africa. This could transfer MDR Enterococci to humans, and cause serious nosocomial infections including endocarditis, bacteremia and urinary tract infections that can result in high morbidities and mortalities.

Ngbede et al. (2017) recently characterized 63 ampicillin- and 37 gentamicin-resistant *E. faecium* from vegetables, soil, farms, animal and manure ^25^. Approximately 95% (35/37 isolates) and 8% (5/63 isolates) of the aminoglycoside- and ampicillin-resistant clones were recognized as high-level aminoglycosides- and ampicillin-resistant *E. faecium* respectively. Modifying enzymes’ genes such as *aac(6’)-Ie-aph(2”)- Ia), aph(2’)-1c,aph(3’)-llla,* and *ant(4’)-la* accounted for the aminoglycoside resistance.

### East Africa: Kenya and Tanzania Tanzania

In a study to determine if cattle co-grazing with wild life influence ABR, ABR in wild animals such as buffalo, zebra and wildebeest was higher than in cattle, although wildlife is periodically treated with antibiotics. Ten VRE and ampicillin-resistant Enterococcus were found in the wild animals but not cattle. Additionally, Enterococcus isolates from wildlife were highly resistant to tetracycline, rifampicin, macrolides, aminoglycosides and cotrimoxazole ^14^. *tet(W)* and *sull* were the resistance genes identified in the isolates. The practice of co-grazing possibly resulted in transmission of ABR genes from livestock to wildlife. The high presence of ABR bacteria in wildlife was likely due to contact with more environmental surfaces that have been contaminated with human, birds or animal excreta. Result from this study demonstrates the presence of ABR Enterococci in wild animals without antibiotic pressure.

### Southern Africa: Angola, Malawi, Mozambique, Namibia, South Africa South Africa

Multiple antibiotic-resistant Enterococci were isolated from borehole water, waste water, pigs and humans in South Africa. Notably, a very high-level vancomycin, aminoglycoside, β-lactam, macrolides and fluoroquinolones resistance was detected among the Enterococci isolates compared to other countries. *erm*(B) (≥300 isolates), *vanC* 2/3(162 isolates), *vanB* (≥138 isolates), *vanC* (≥120 isolates), *strA* (≥120 isolates) were the major resistance genes. The vancomycin-resistant isolates were from patients with haematological malignancies, bacteremia, pigs, wastewater and underground water ^9,10,26,30^. Inefficient chlorination to kill bacteria accounted for the high resistance rates in the final effluents’ discharge into the environment. Hospital wastewater is therefore a major source of MDR Enterococcus. Sub-therapeutic antibiotic usage in animal feed also accounted for the emergence of ABR in pigs whilst the construction of boreholes near pit toilets resulted in high enterococcal isolation and resistance rates in South Africa.

#### Experimental procedures used in included studies

The studies included in this review basically used the following experimental procedures. Transport media such as stuart agar, cary-blair medium, and gel transport swabs with charcoal were used to transport the samples to the laboratory ^53,65^. Cotton swabs were used to swab sample specimens, tissues, surfaces, fluids, etc. and cultured on nutrient agar, blood agar, tryptone soya agar, mannitol salt-phenol red agar, brain-heart infusion broth, Slanetz-Bartley mannitol salt agar, and Edwards agar media prior to identifying the 24-hour colonies using Gram-staining and different biochemical tests such as catalase and coagulase tests, latex coagulase test and DNase agar test. Subsequently, antimicrobial susceptibility testing (AST) using disc diffusion (Kirby-Bauer method or E-test) on Mueller Hinton agar plates and a 0.5 McFarland bacterial inoculum was performed. Antibiotics such as ampicillin (AMP), amoxicillin (AMX), amikacin (AMK), ampicillin-Sulbactam (SAM), amoxicillin-clavulanic acid (AMC), azithromycin (AZI), apramycin (APR), chloramphenicol (CHL), cefoxitin (FOX), ceftazidime (CFZ), clarithromycin (CLR), ciprofloxacin (CIP), cefuroxime (CXM), clindamycin (CLI), cephalexin(LEX), cefoperazone (CFP), cefepime (FEP), cefotaxime (CTX), ceftaroline (CPT), cephalothin (CET), cloxacillin (CLX), doxycycline (DOX), erythromycin (ERY), fusidic acid (FUS), fosfomycin (Fof), gatifloxacin (GAT), gentamicin (GEN), imipenem (IPM), kanamycin (KAN), levofloxacin (LVX), linezolid (LZD), lincomycin (LIN), meropenem (MER), mupirocin (MUP), minocycline (MIC), moxifloxacin (MXF), methicillin (MET), metronidazole (MTZ), nitrofurantoin (NIT), norfloxacin (Nor), nalidixic acid (NAL), netilmicin (NEL), oxacillin (OXA), ofloxacin (OFX), perfloxacin (PF), penicillin (PEN), pristinamycin (PRI), rifampicin (RIF), streptomycin (STR), streptogramin B (SB), sulfamethoxazole (SMZ), tetracycline (TET), teicoplanin (TEC), telithromycin (TEL), tobramycin (TOB), trimethoprim-sulfamethoxazole (SXT), and vancomycin (VAN) were mostly used for the AST. Polymerase chain reaction (PCR) was used to detect the antimicrobial resistance genes and clones (i.e. molecular typing) of the isolates.

## 3. CONCLUSION AND STUDY LIMITATIONS

We report of high rate of ABR among GPB in several African countries, mediated largely by *S. aureus* ST5, ST8, and ST80, *Enterococcus faecium* and *Enterococcus faecalis* strains, *SCCmec,* Tn*916* and IS*16* MGEs are a major threat to clinical medicine, the economy and socio-economic development. This calls for national as well as international rules and regulations to contain resistance. Heavy consumption of antibiotics in animal feed, exchange of resistance genes between animals and food animal products to man, uncontrolled and inappropriate antibiotics prescription practices, inadequate hygienic handling and processing of food, close contact with pet dogs, shedding of resistant clones from animals to humans and the environment, as well as high consumption of antibiotics in humans, particularly in HIV patients, account for the high rate of ABR in Africa.

Effective surveillance and monitoring of antimicrobial drug usage and licensing, banning or restricting the prescription of reserved, expired and substandard drugs, periodic monitoring of pharmacies and veterinary shops, and antibiotic stewardship are recommended measures to contain ABR. Improving animal health through hygienic practices on farms, avoiding prophylactic or growth-promoting antibiotic usage in veterinary medicine, integrative efforts between human and veterinary medicine as well as environmental health are urgently needed to contain ABR. Implementation of these policies will decrease the high rate of ABR in Africa, reduce longer hospital stays and the resort to expensive but toxic antibiotic alternatives, with a concomitant reduction in morbidity and mortality rates. Few studies reporting on the molecular determinants of ABR in GPB in Africa limited the study to 130 articles. Among these, only few studies reported on MGEs and resistant clones.

## Role of Funding Source

Not applicable.

## Contributors

JOS conceived, designed and supervised the study, analysed and vetted the results, wrote the paper, edited and formatted it for publication. EM co-conceived and co-designed the study, gathered and analysed the data and drafted the paper. Both authors approved the final version for submission.

## Funding

None

## Declaration of interests

The authors declare no conflict of interest.

## Acknowledgments

None

---

**Table 1.**
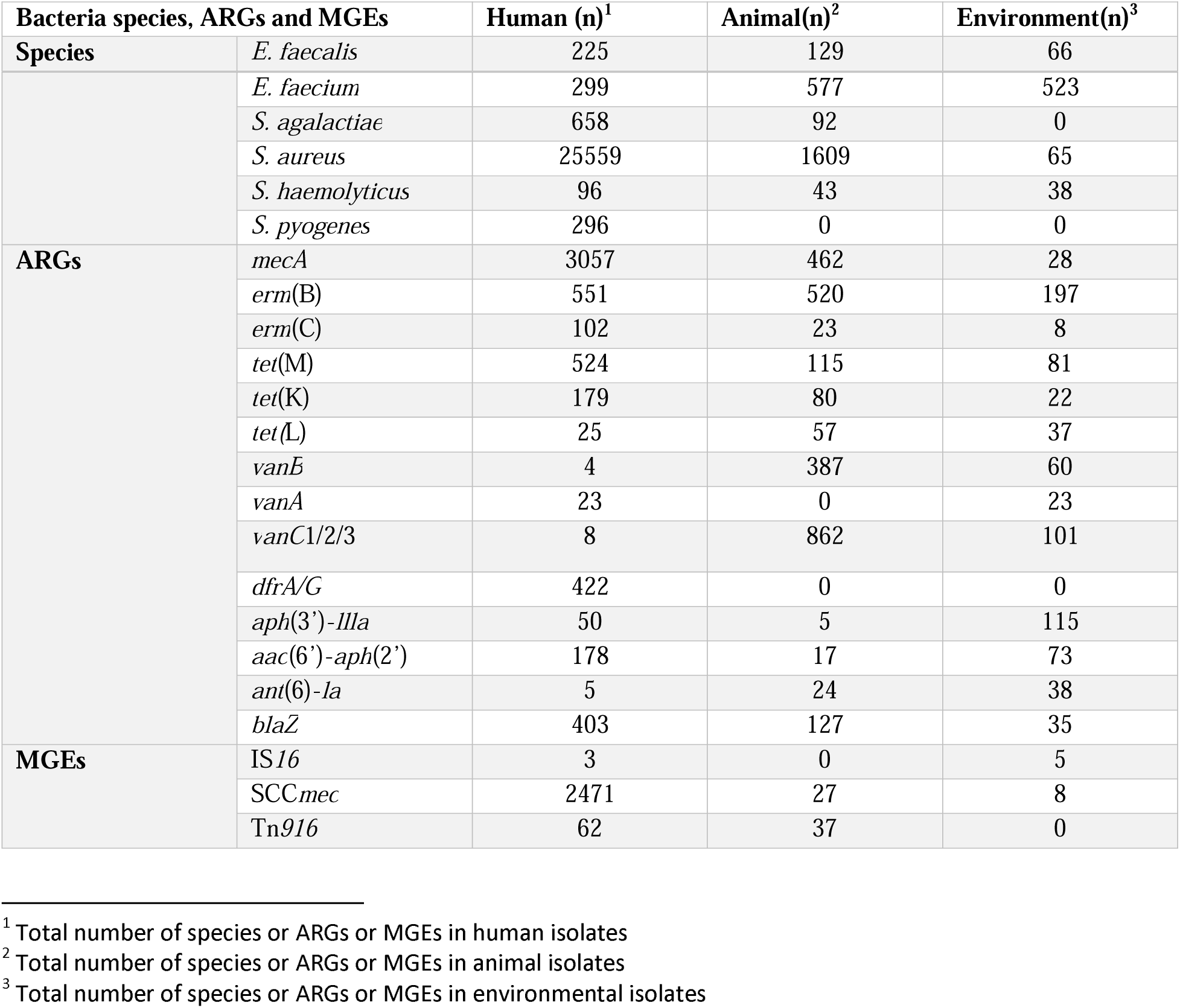
Frequency distribution of Gram-positive bacterial species, resistance genes and MGEs isolated from animals, humans and environmental specimens.

**Table 2.**
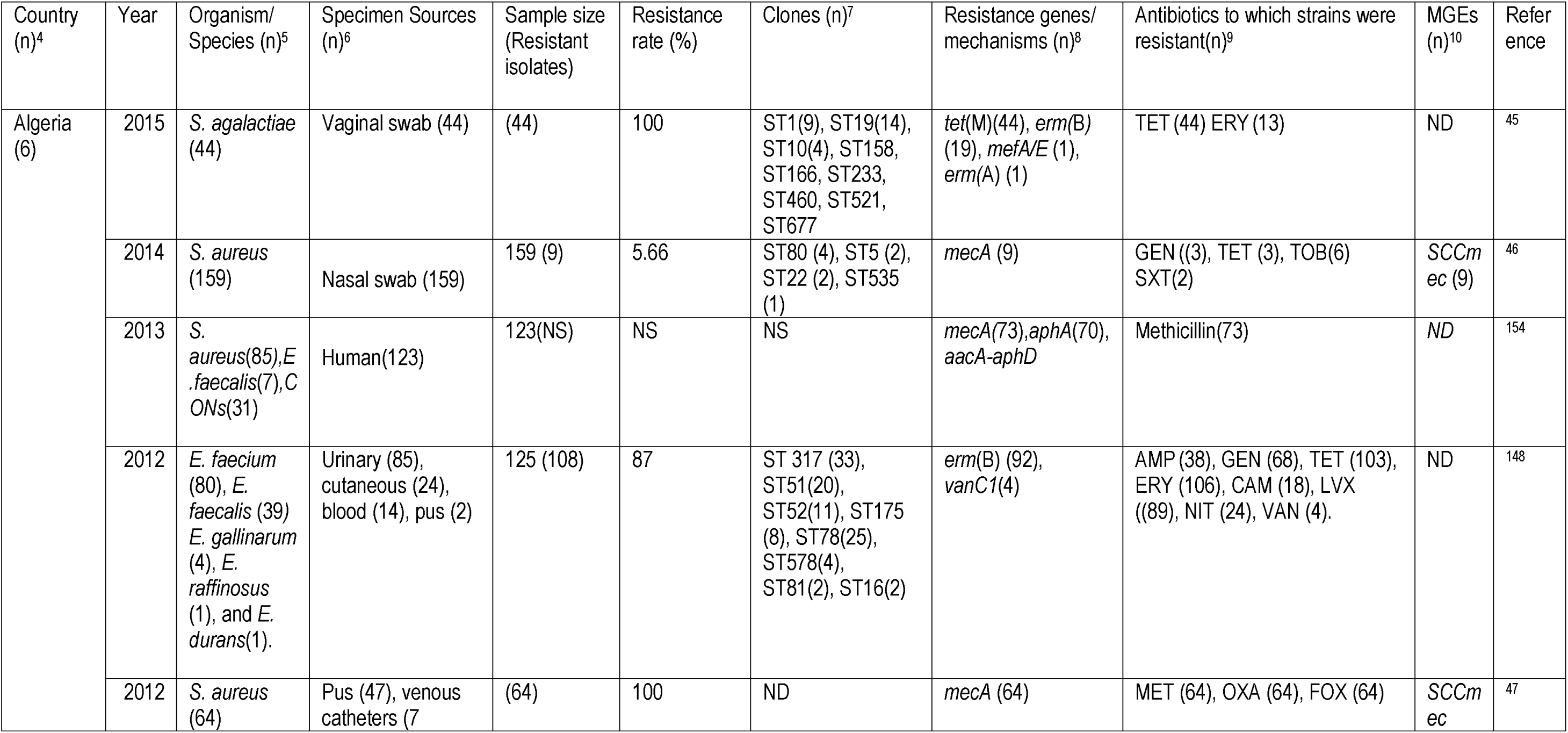
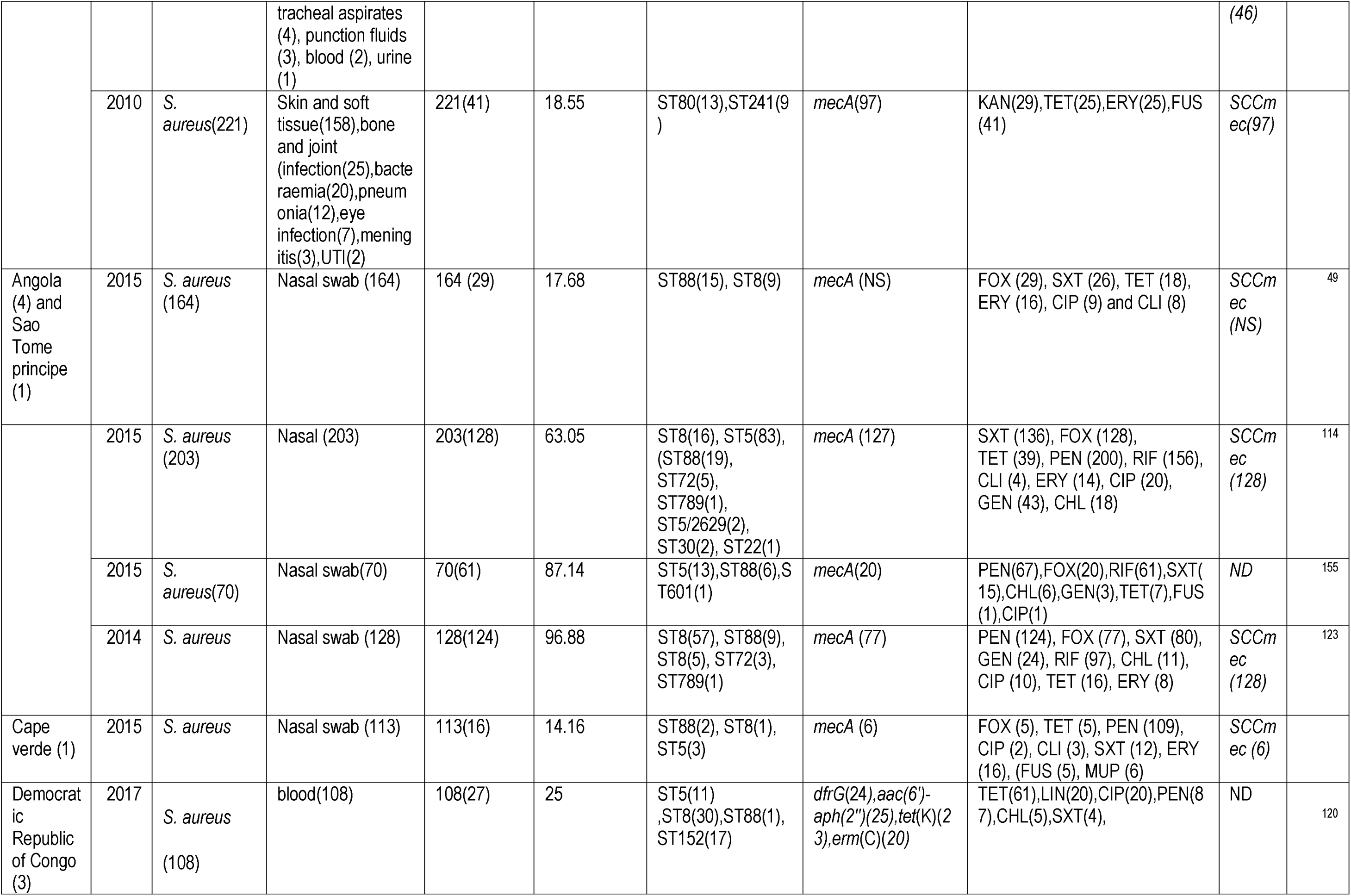
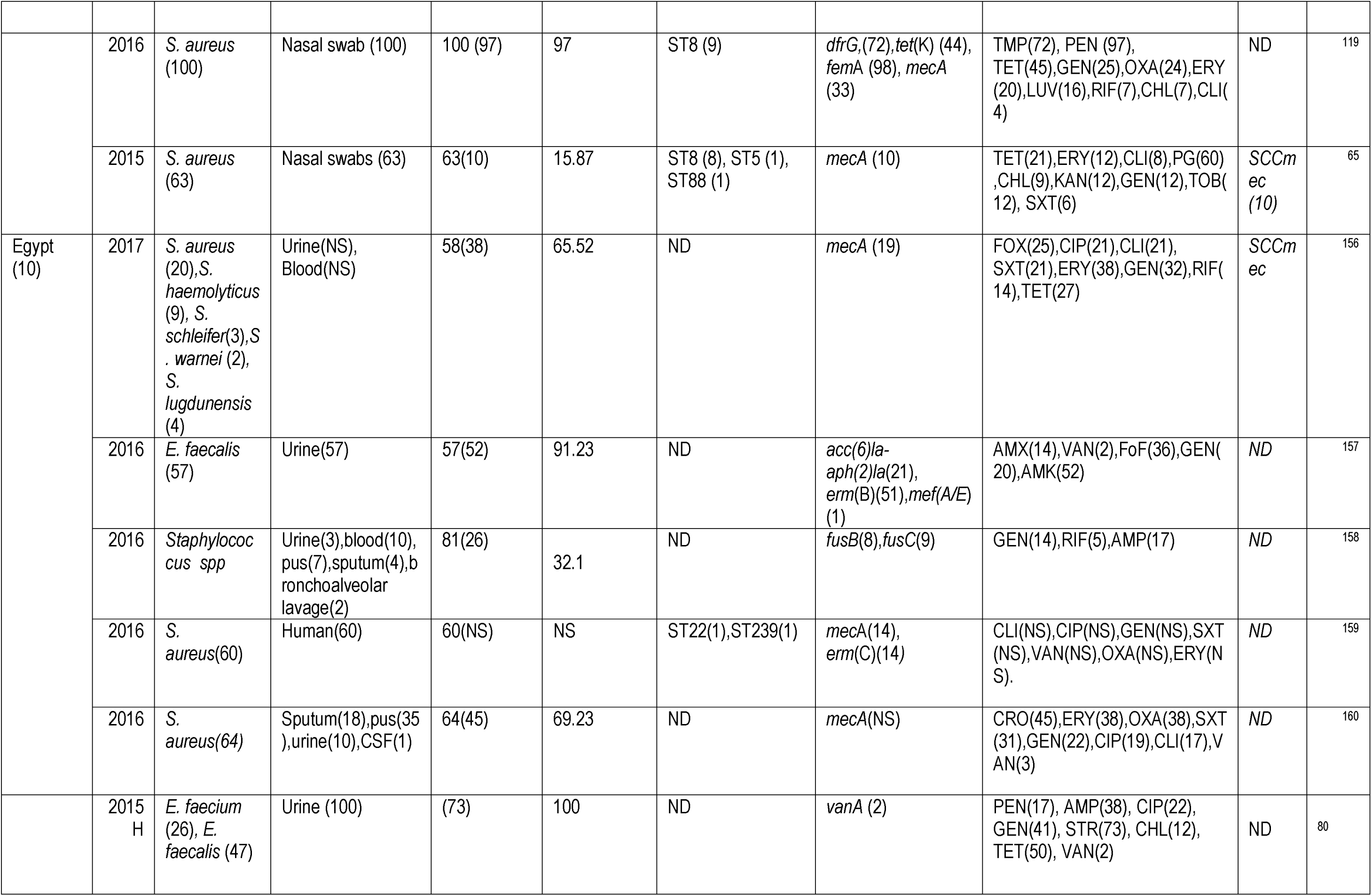
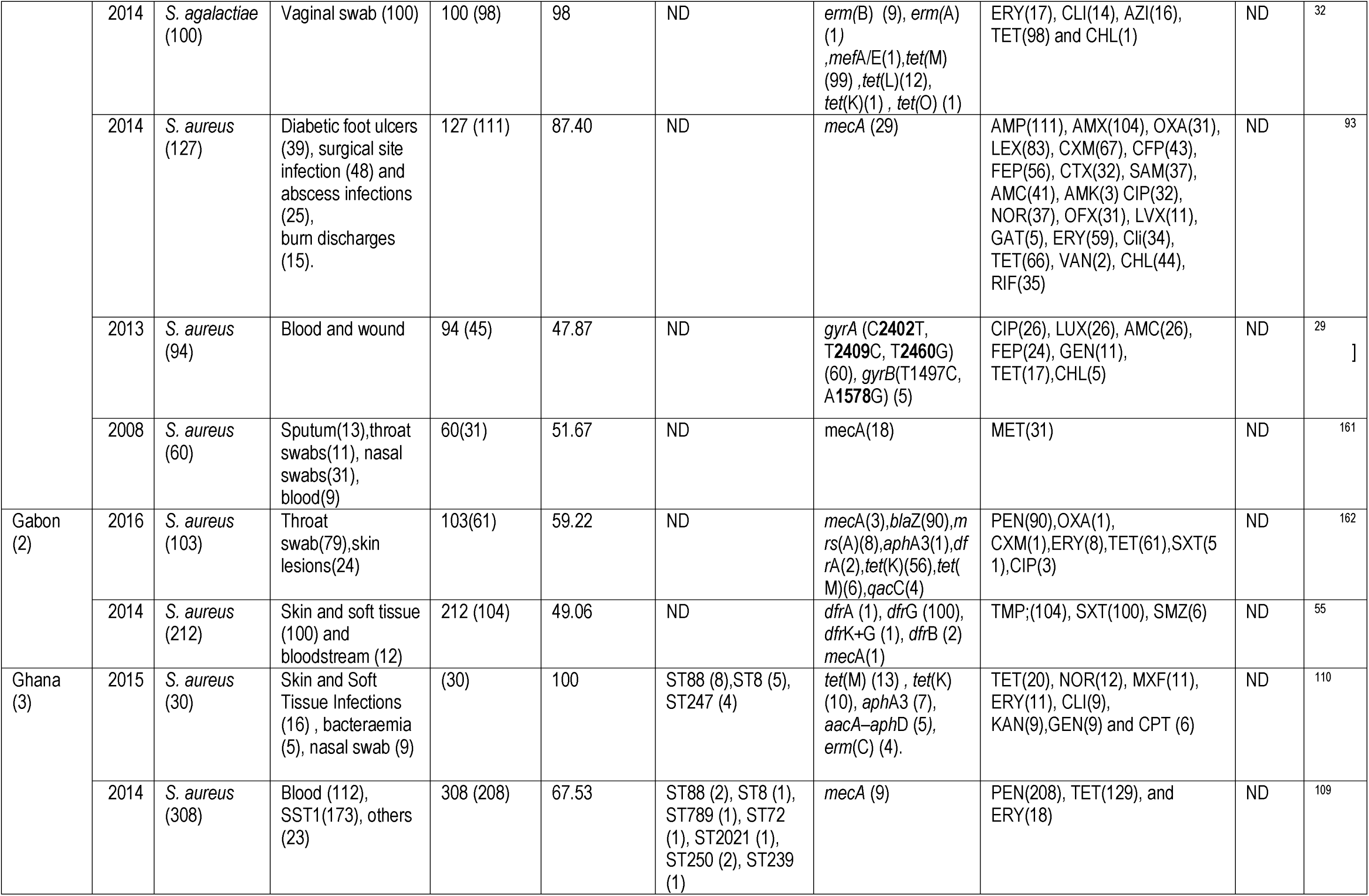
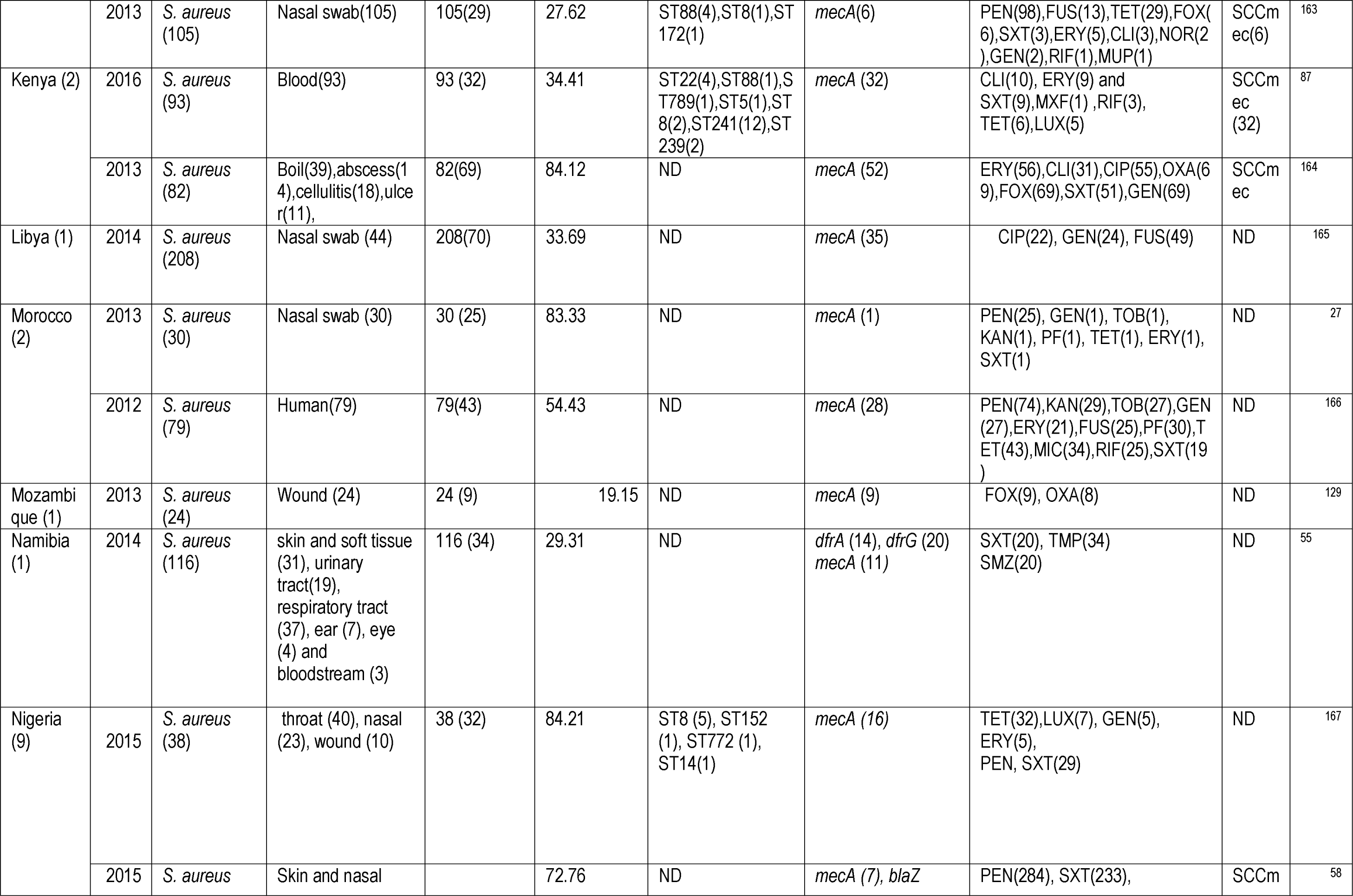
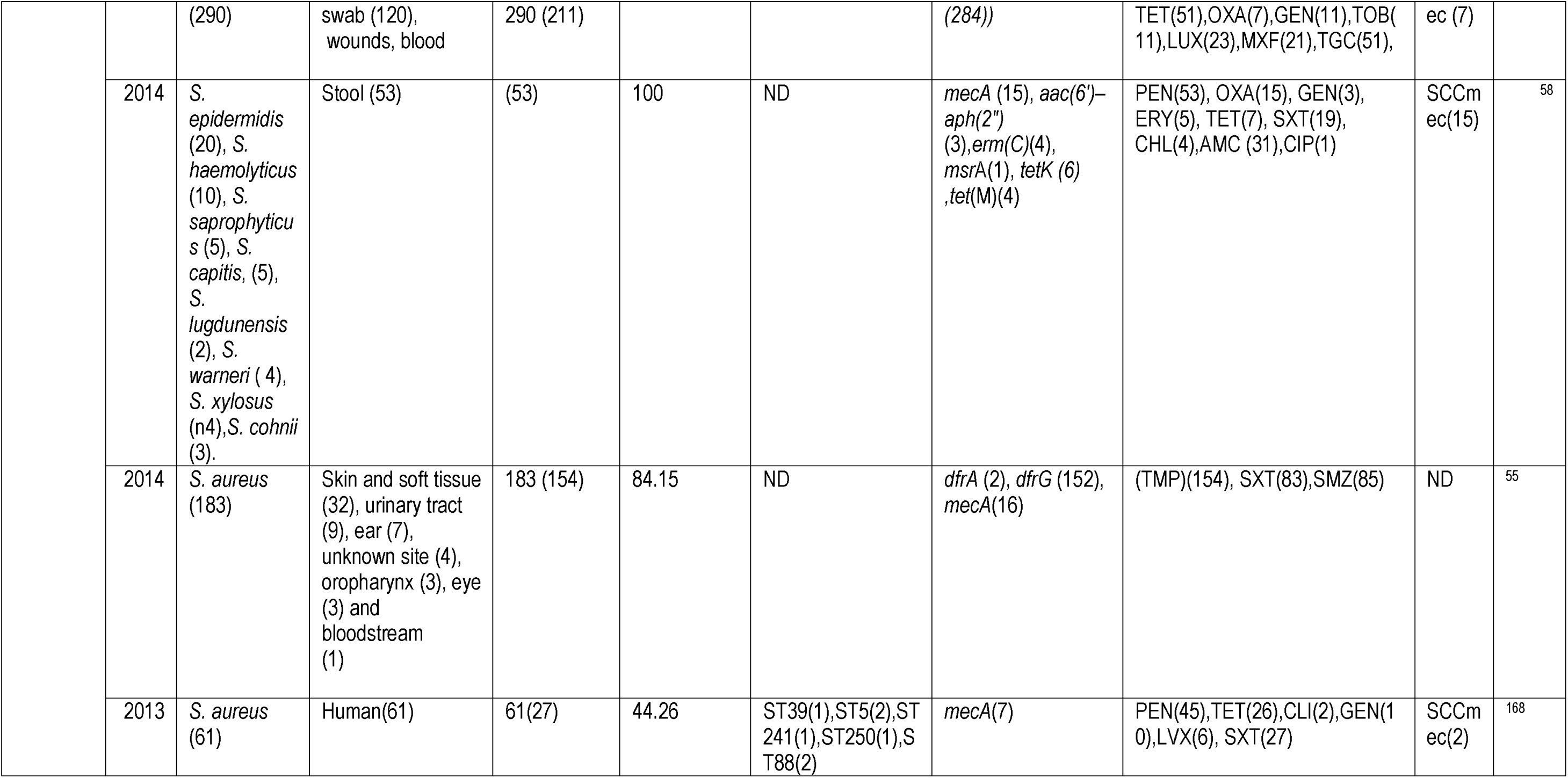
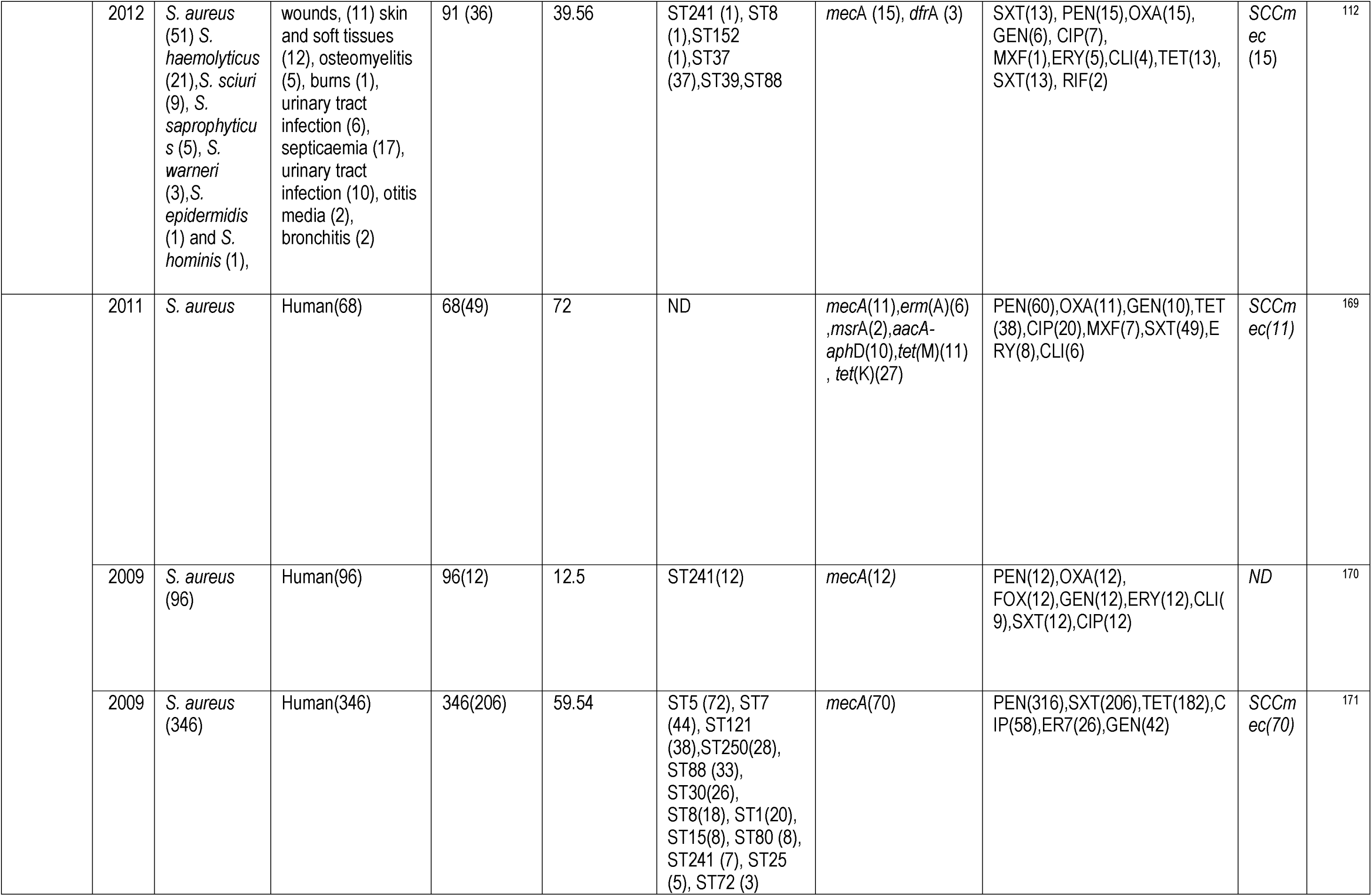
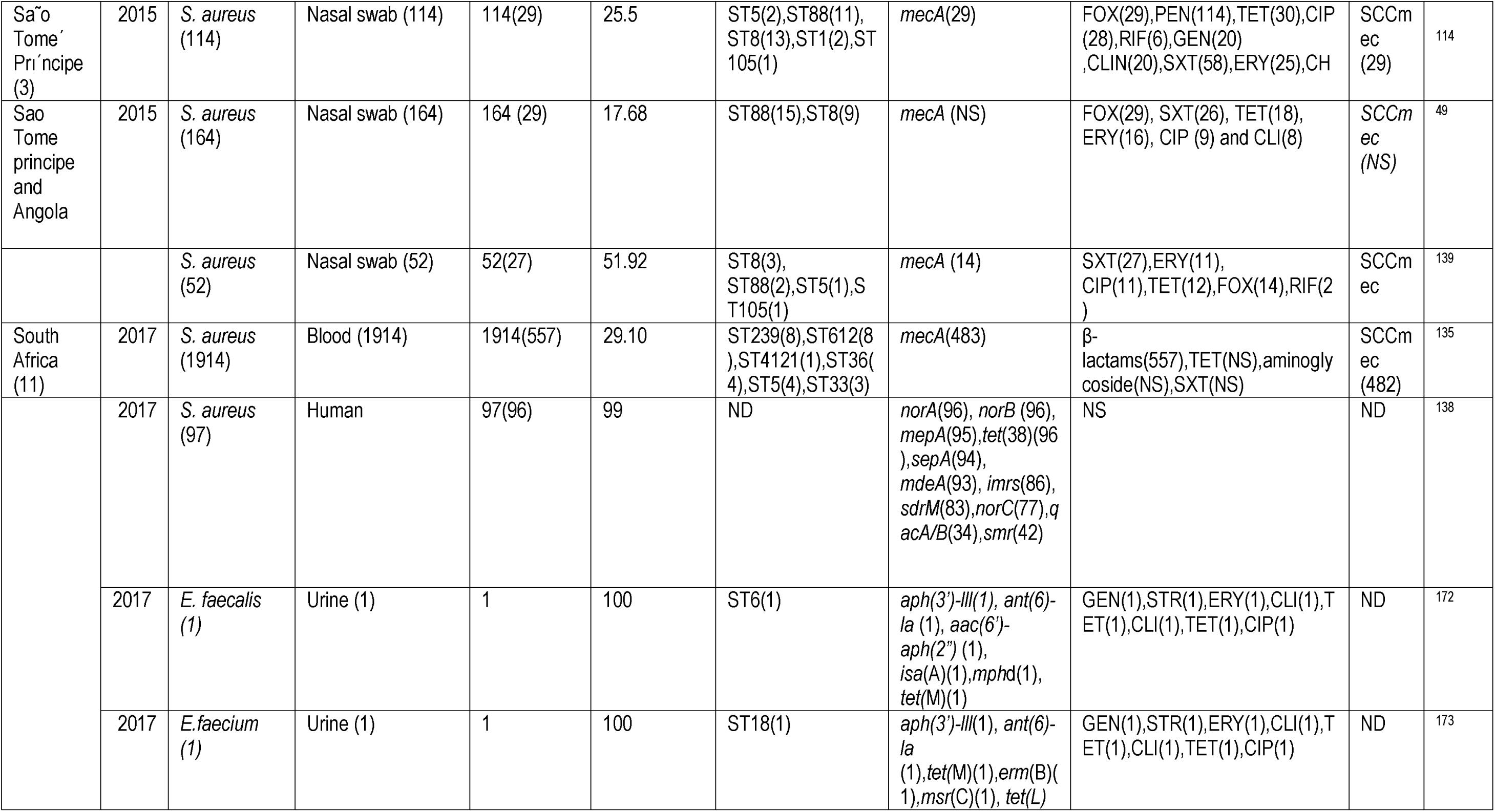
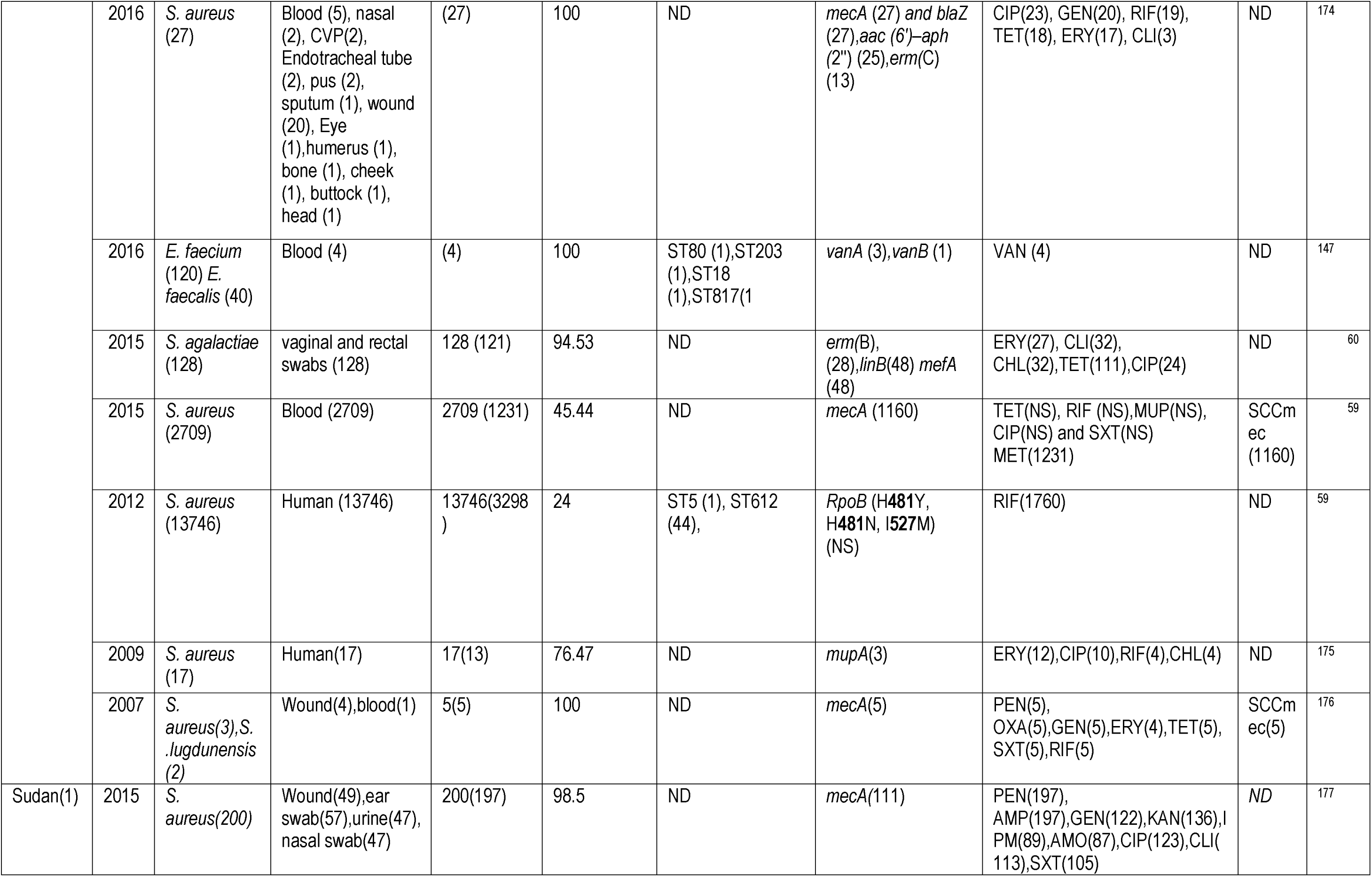
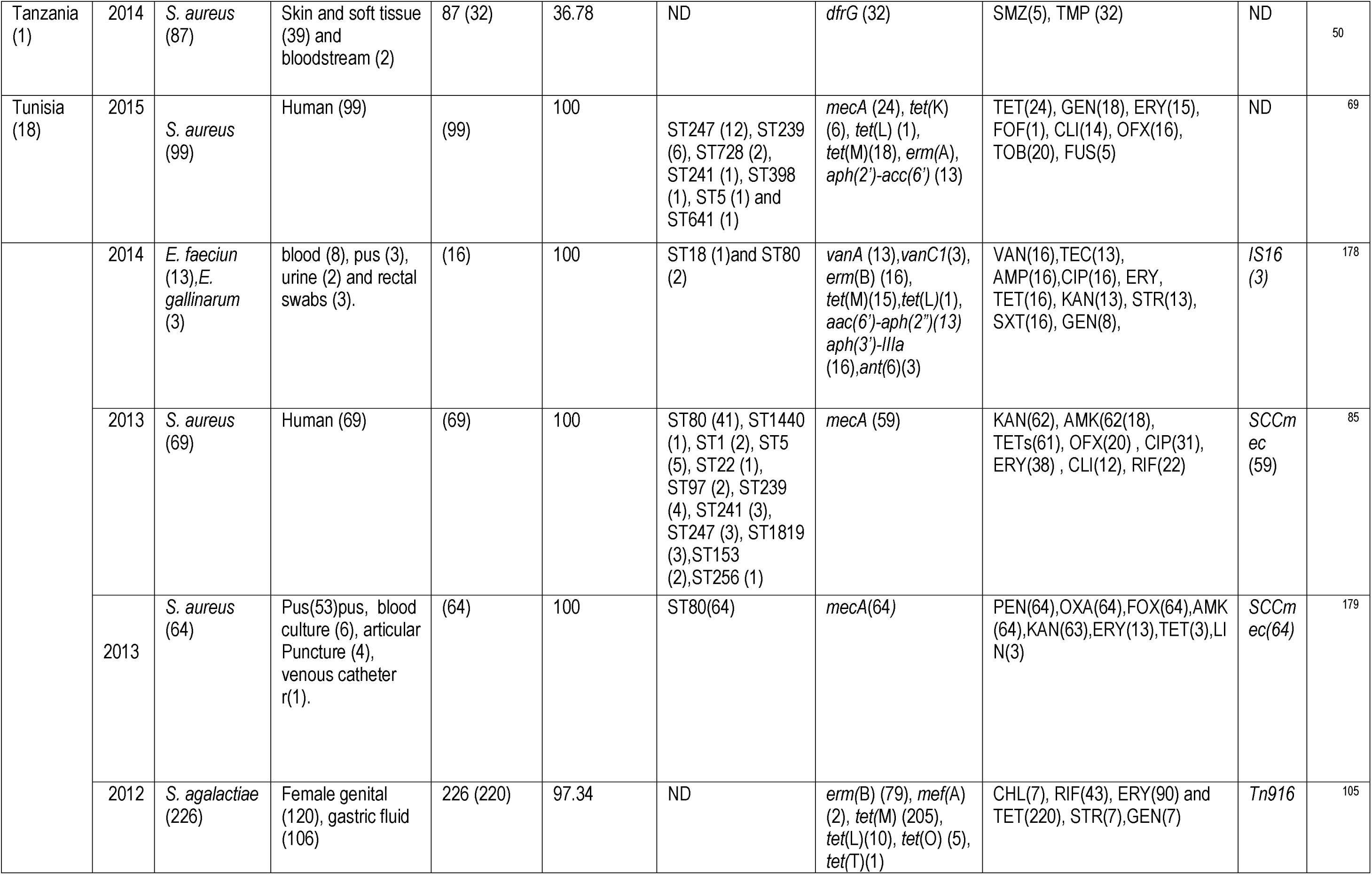
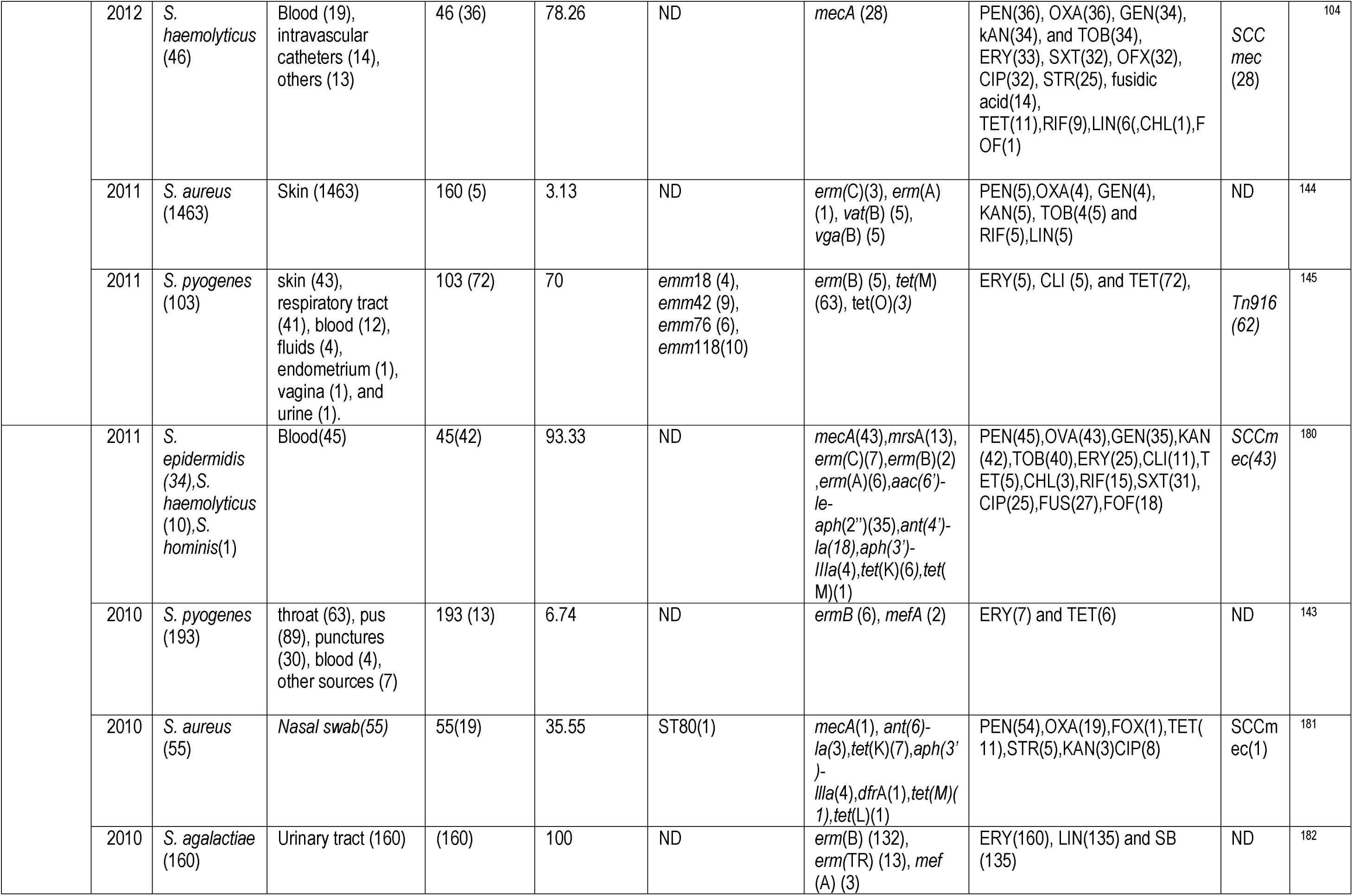
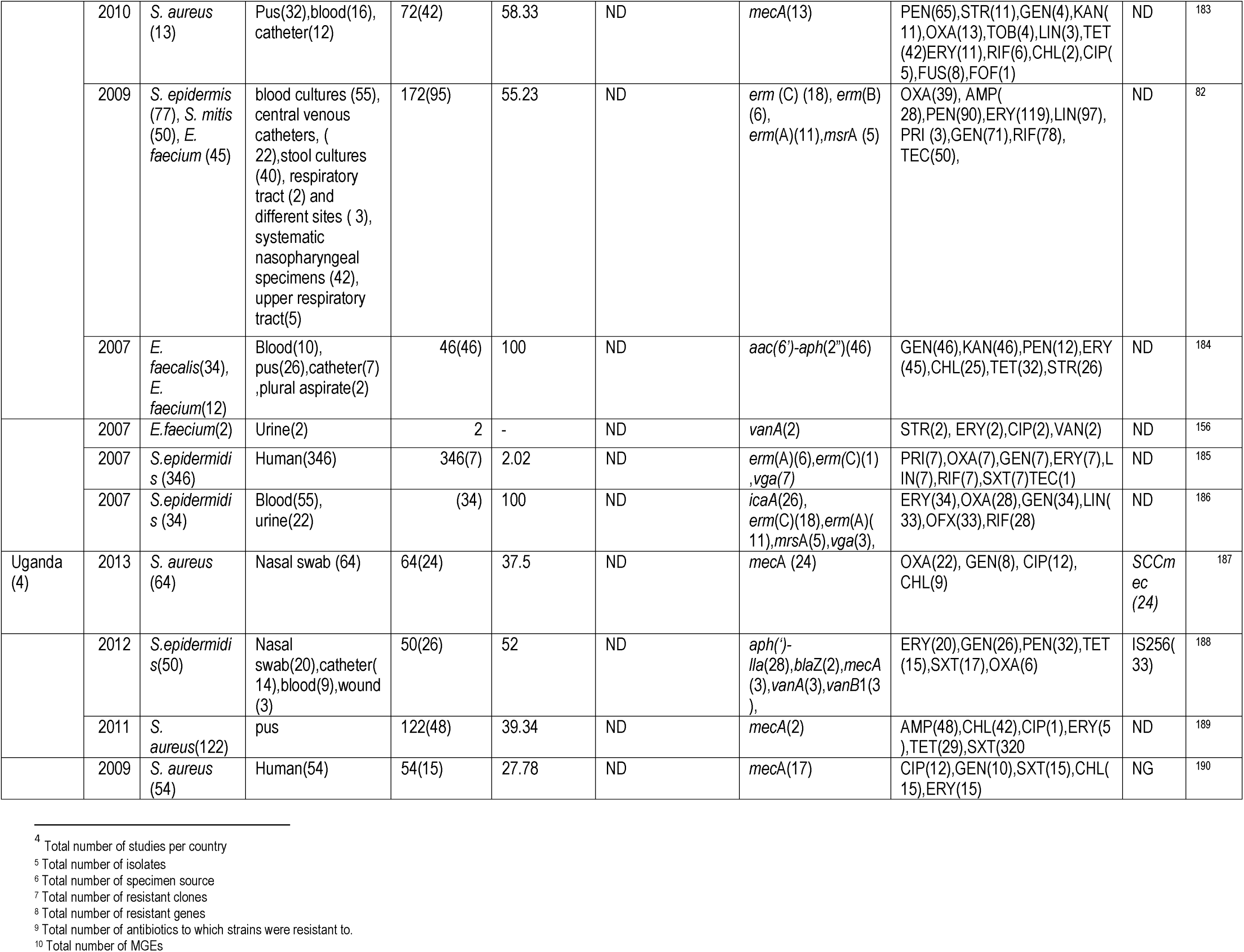
Geographical distribution, species, clones, and resistance mechanisms of antibiotic-resistant Gram-positive bacteria isolated from humans in Africa from 2007-2018.

**Table 3.**
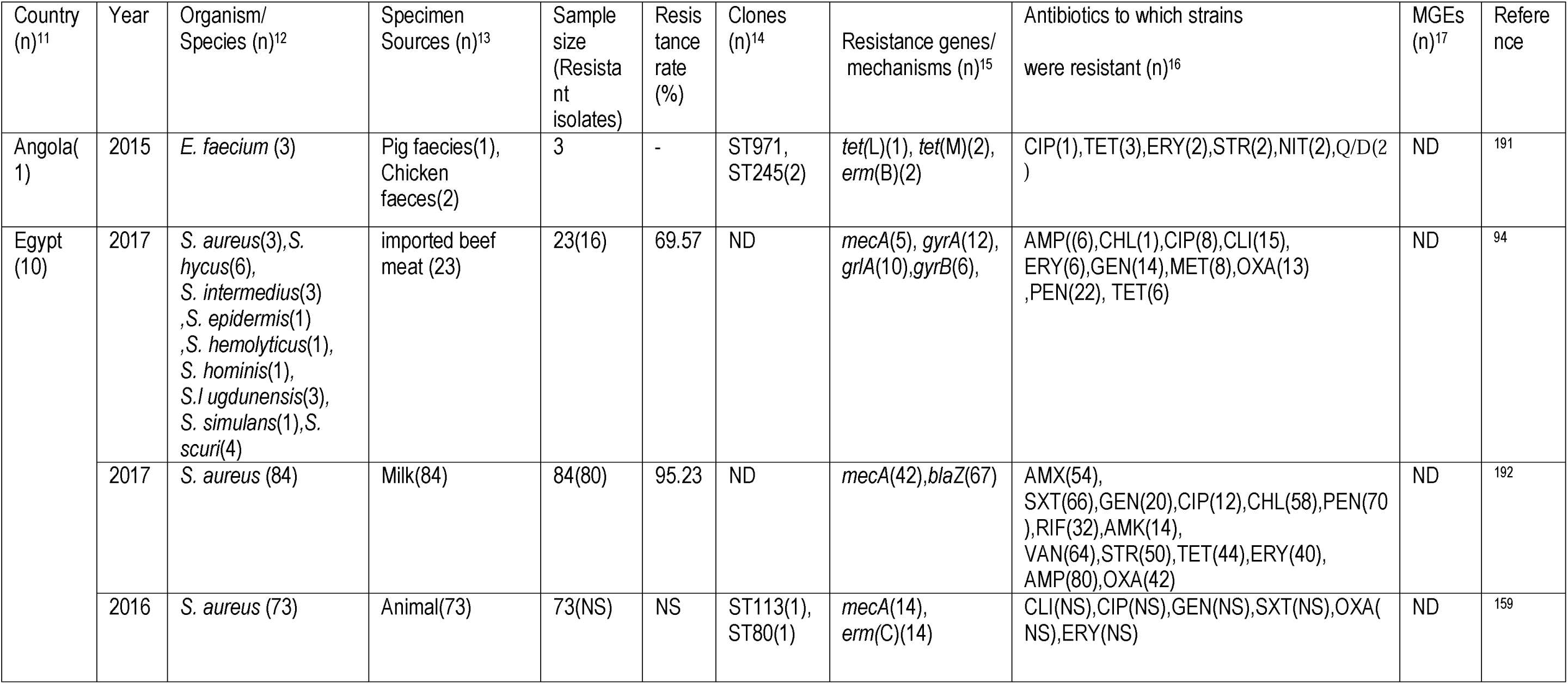
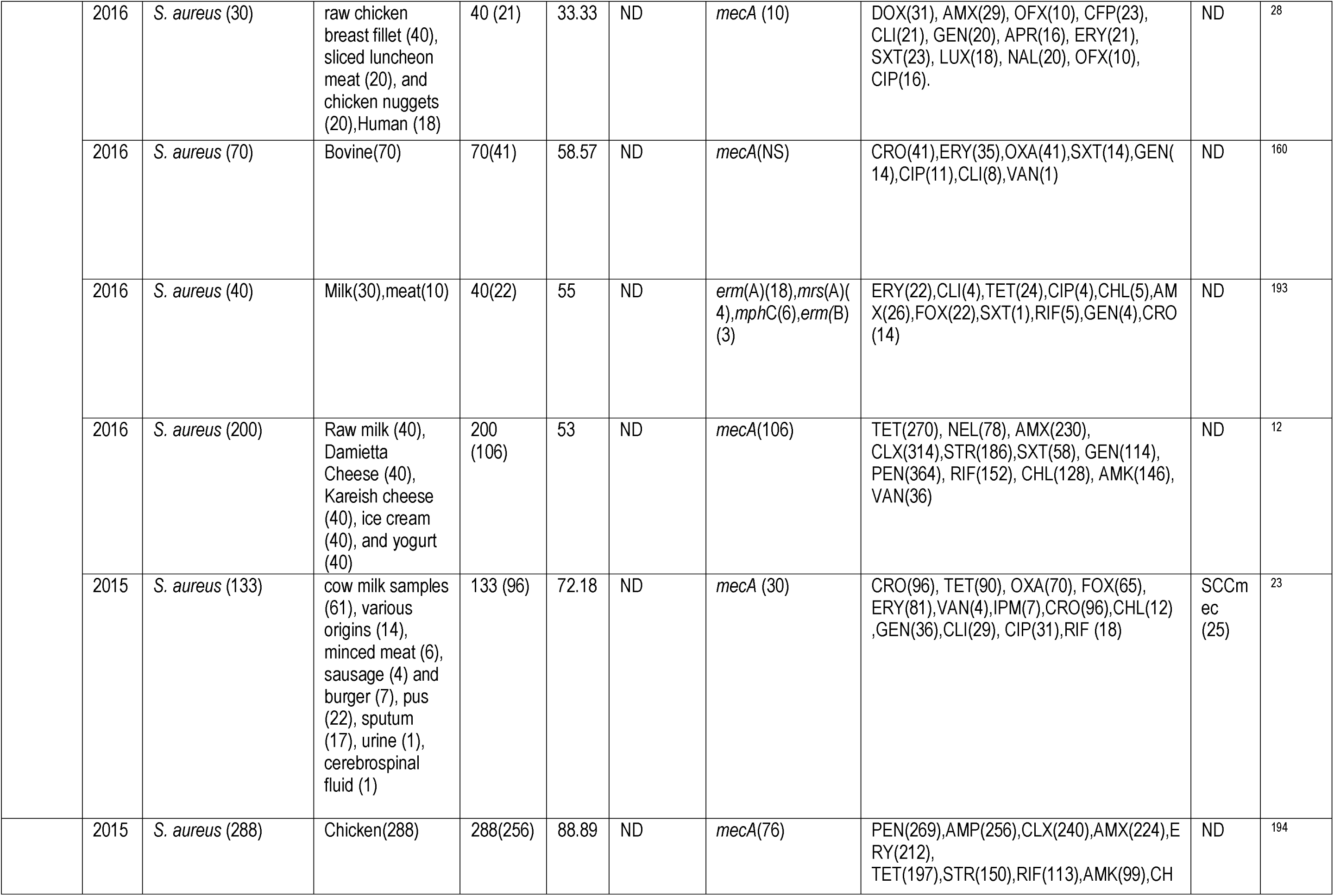
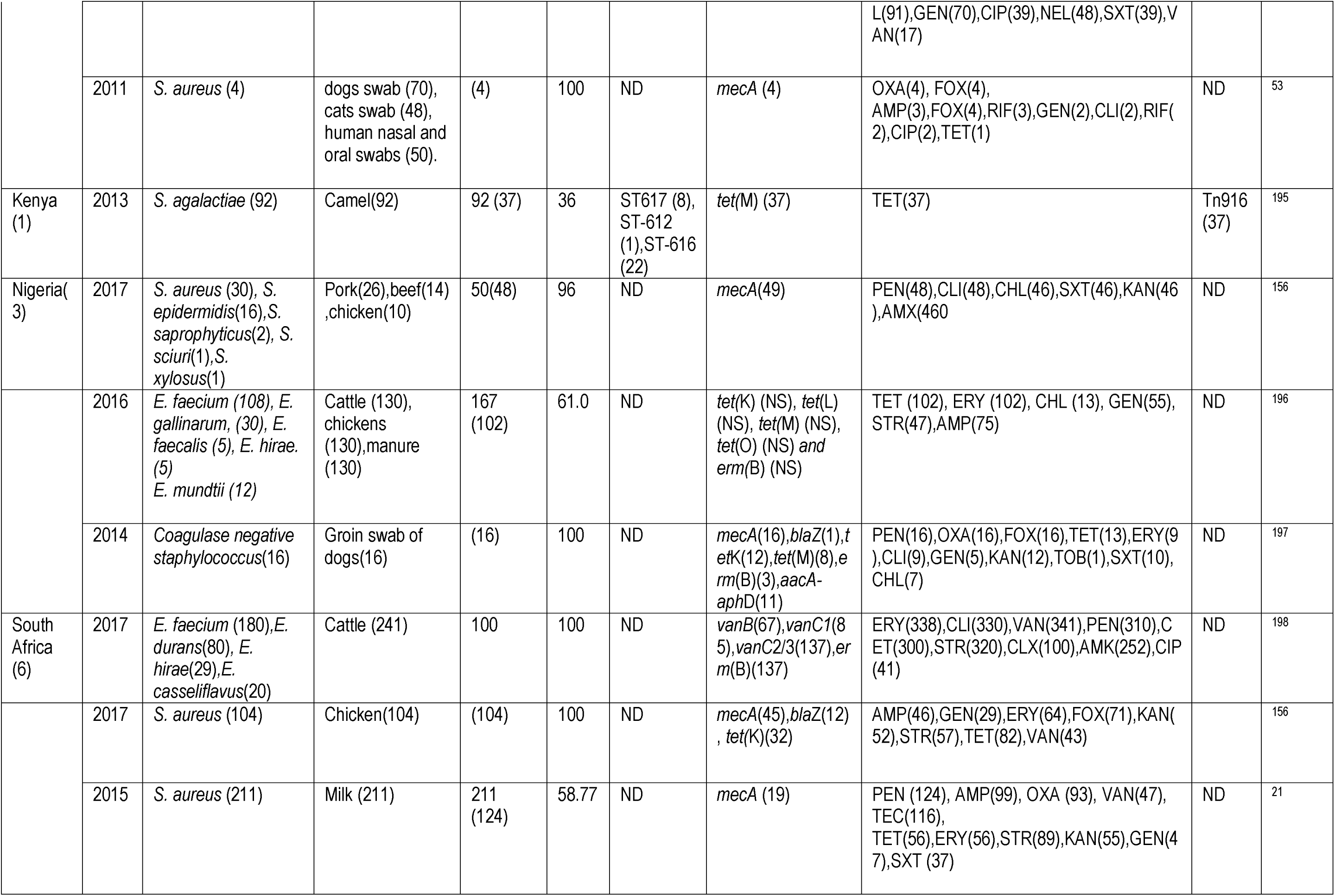
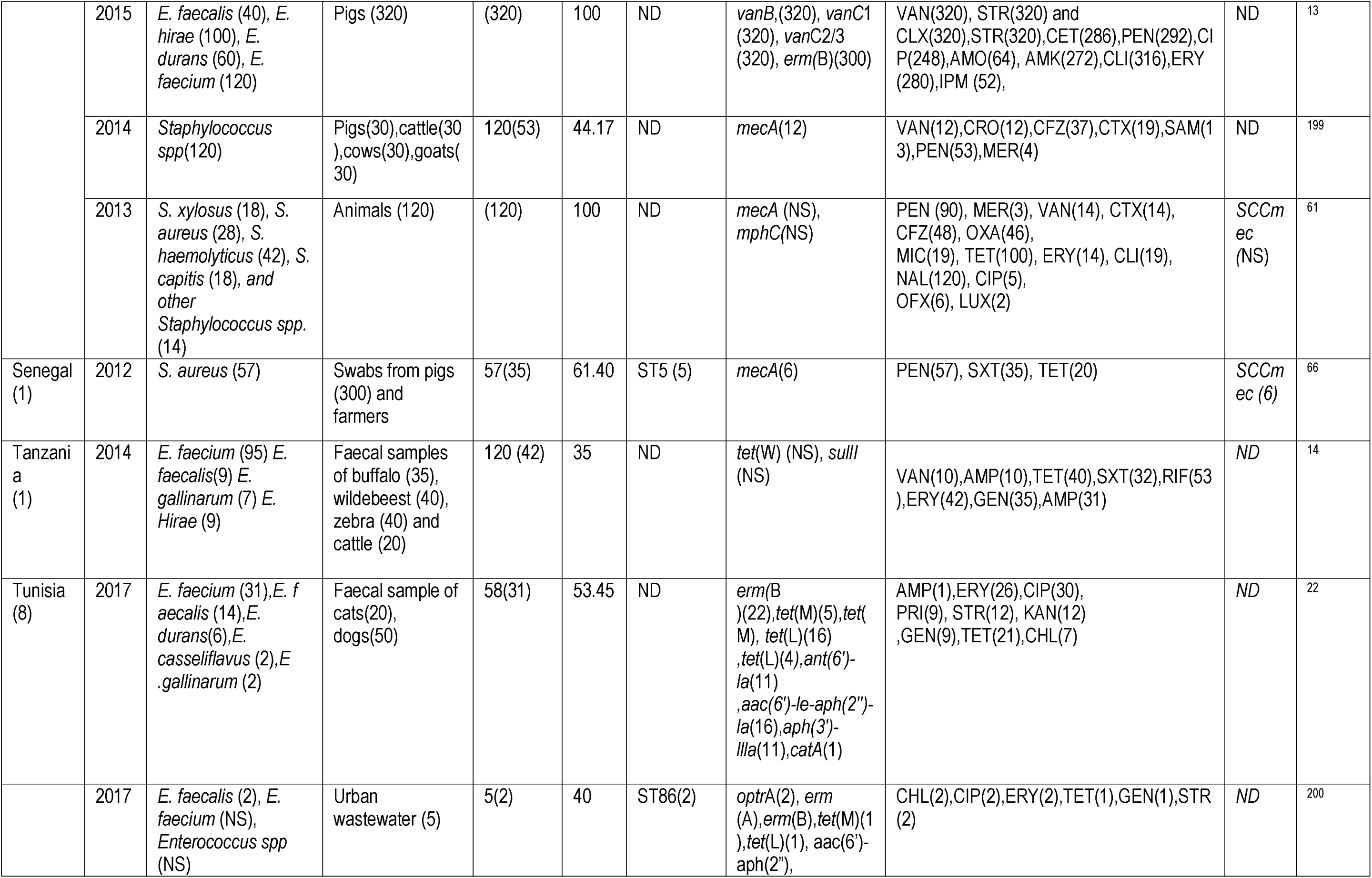
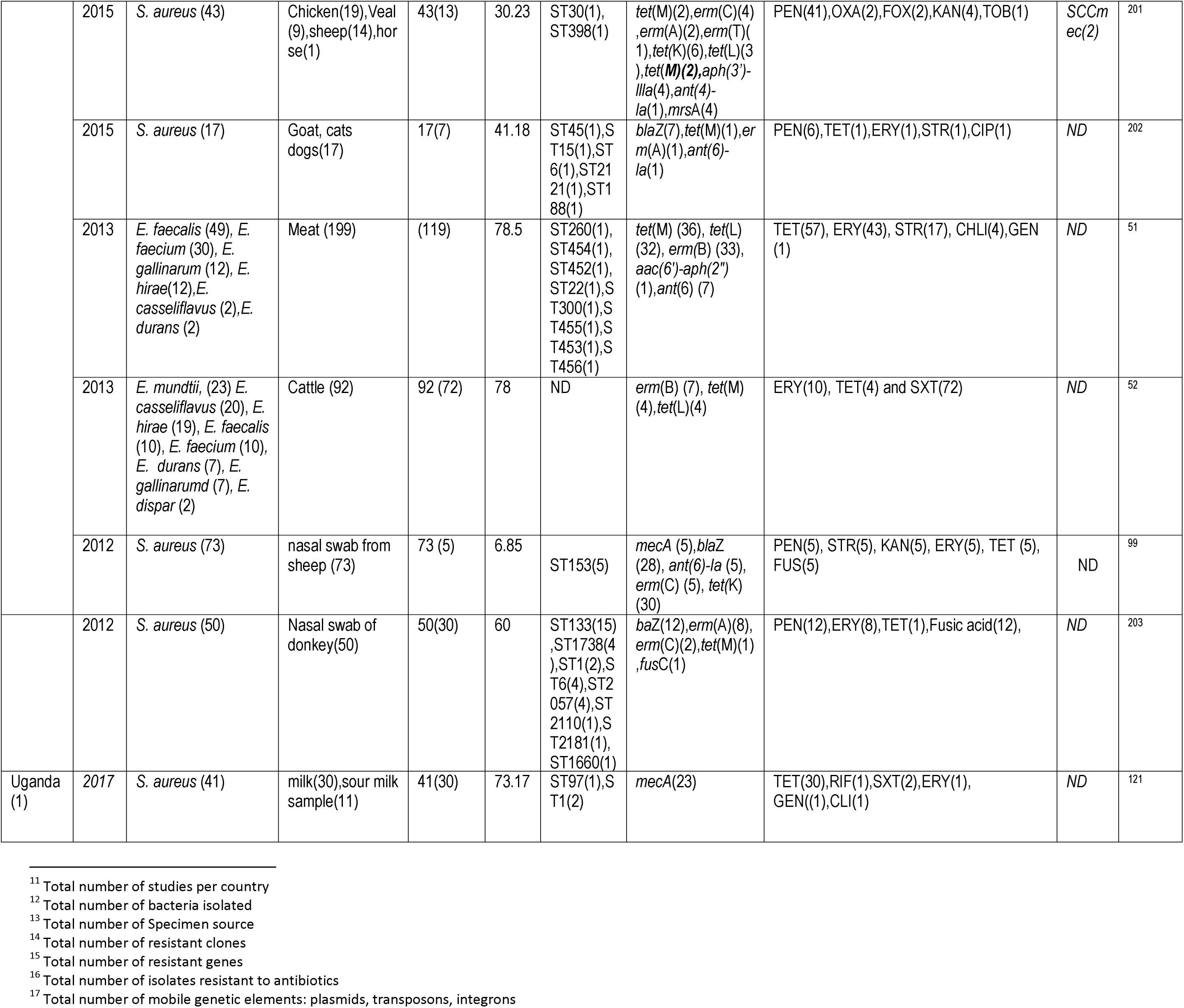
Geographical distribution, species, clones, and resistance mechanisms of antibiotic-resistant Gram-positive bacteria isolated from animals in Africa from 2007-2018.

**Table 4.**
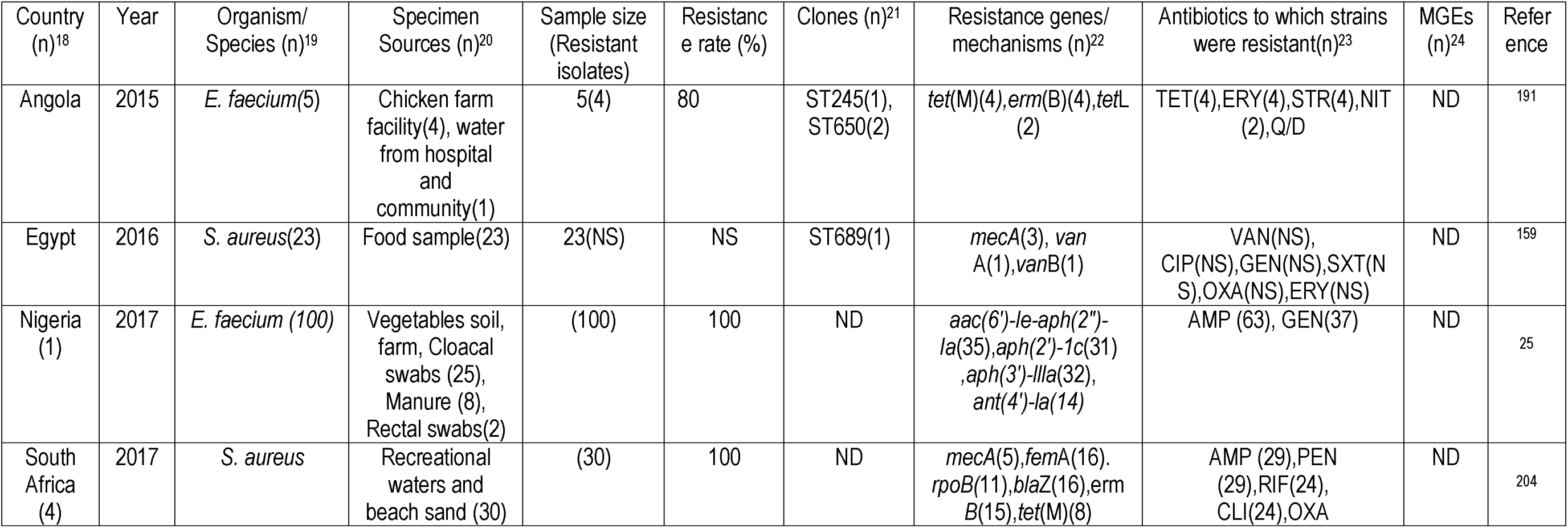
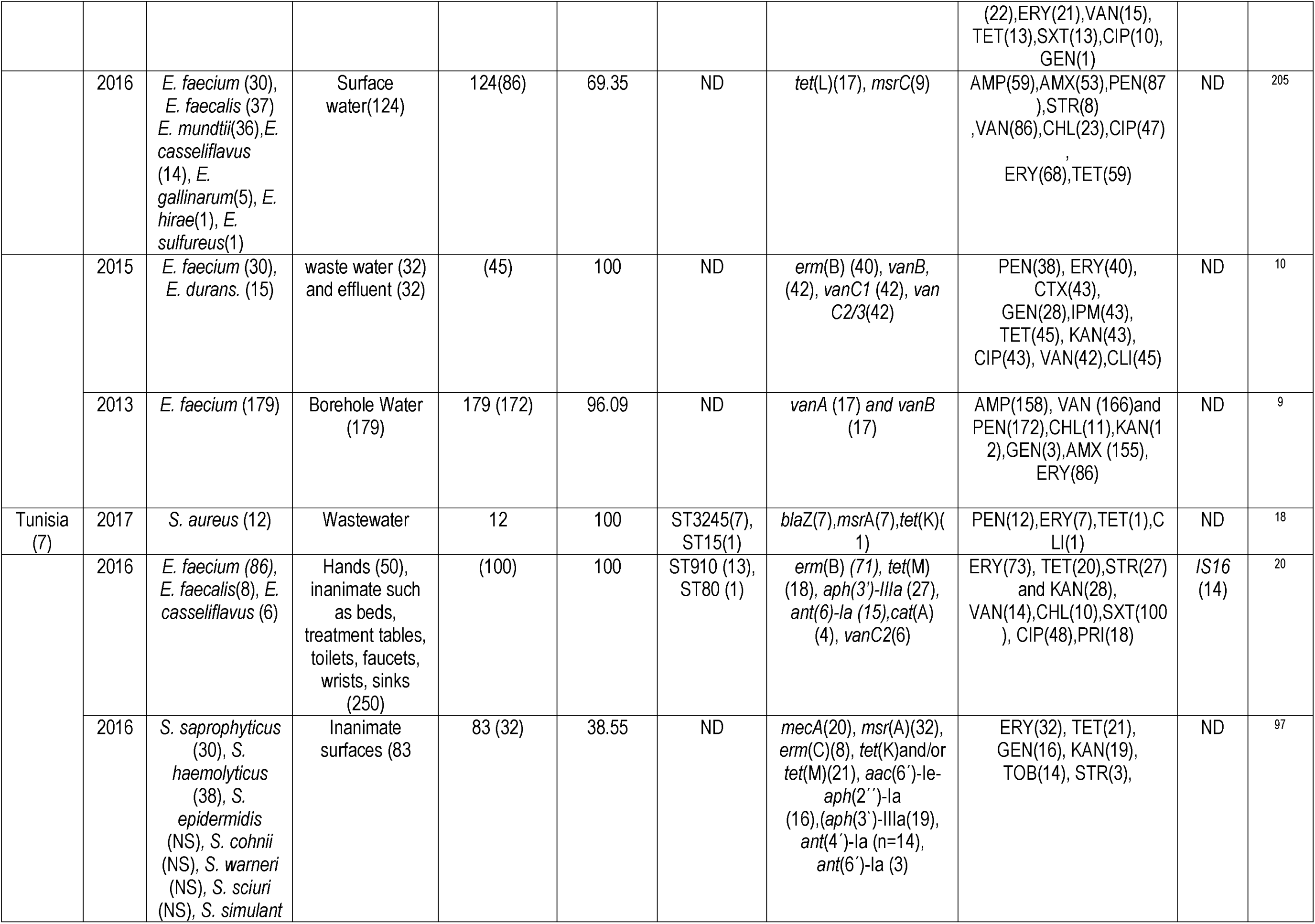
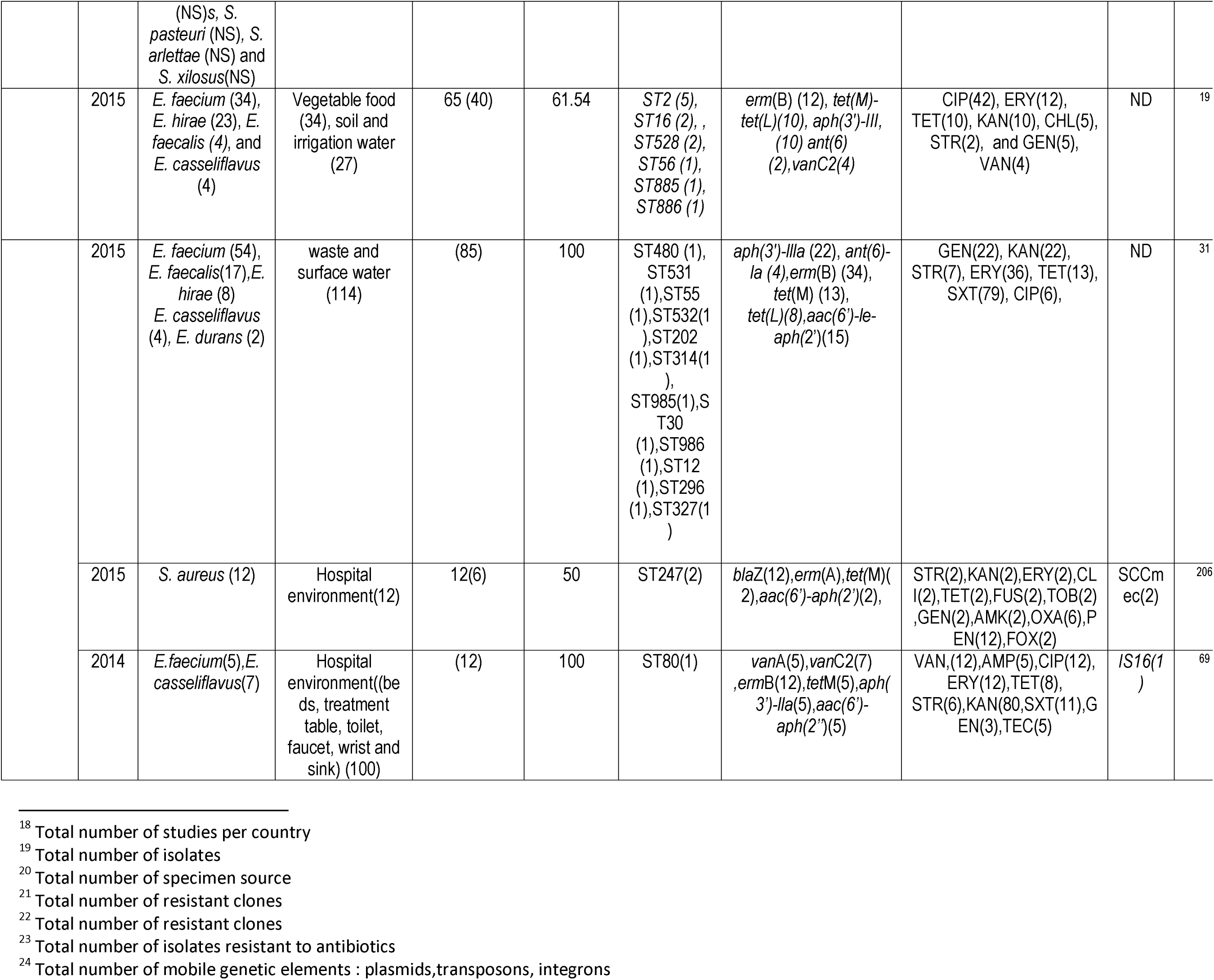
Geographical distribution, species, clones, and resistance mechanisms of antibiotic-resistant Gram-positive bacteria isolated from the environment in Africa from 2007-2017.

**Table 5.**
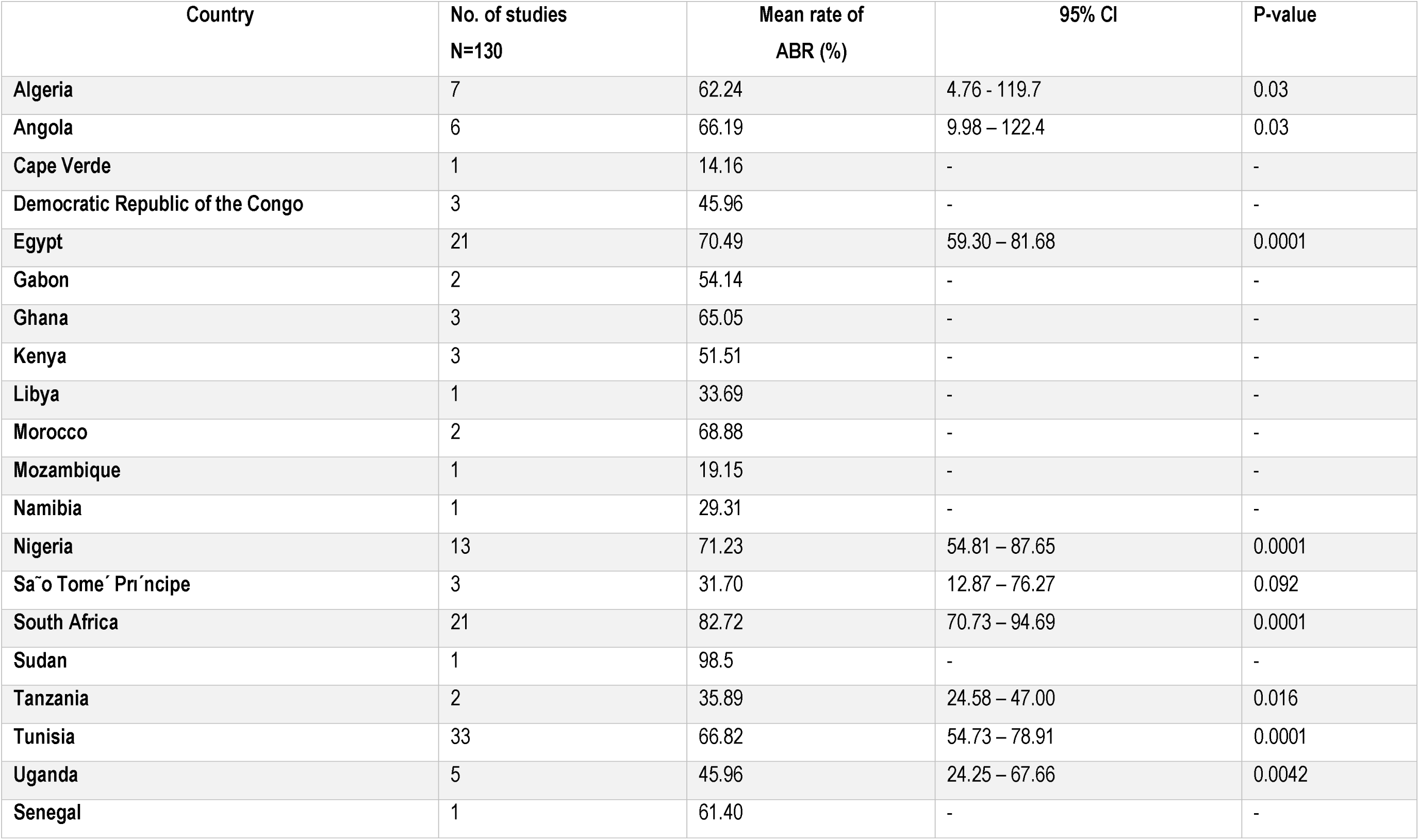
Mean antibiotic resistance rates per country in Africa.

| Country | No. of studies<br>N=130 | Mean rate of<br>ABR (%) | 95% CI | P-value |
| --- | --- | --- | --- | --- |
| Algeria | 7 | 62.24 | 4.76 - 119.7 | 0.03 |
| Angola | 6 | 66.19 | 9.98 – 122.4 | 0.03 |
| Cape Verde | 1 | 14.16 | - | - |
| Democratic Republic of the Congo | 3 | 45.96 | - | - |
| Egypt | 21 | 70.49 | 59.30 – 81.68 | 0.0001 |
| Gabon | 2 | 54.14 | - | - |
| Ghana | 3 | 65.05 | - | - |
| Kenya | 3 | 51.51 | - | - |
| Libya | 1 | 33.69 | - | - |
| Morocco | 2 | 68.88 | - | - |
| Mozambique | 1 | 19.15 | - | - |
| Namibia | 1 | 29.31 | - | - |
| Nigeria | 13 | 71.23 | 54.81 – 87.65 | 0.0001 |
| São Tome´ Pri´ncipe | 3 | 31.70 | 12.87 – 76.27 | 0.092 |
| South Africa | 21 | 82.72 | 70.73 – 94.69 | 0.0001 |
| Sudan | 1 | 98.5 | - | - |
| Tanzania | 2 | 35.89 | 24.58 – 47.00 | 0.016 |
| Tunisia | 33 | 66.82 | 54.73 – 78.91 | 0.0001 |
| Uganda | 5 | 45.96 | 24.25 – 67.66 | 0.0042 |
| Senegal | 1 | 61.40 | - | - |

**Table 6.**
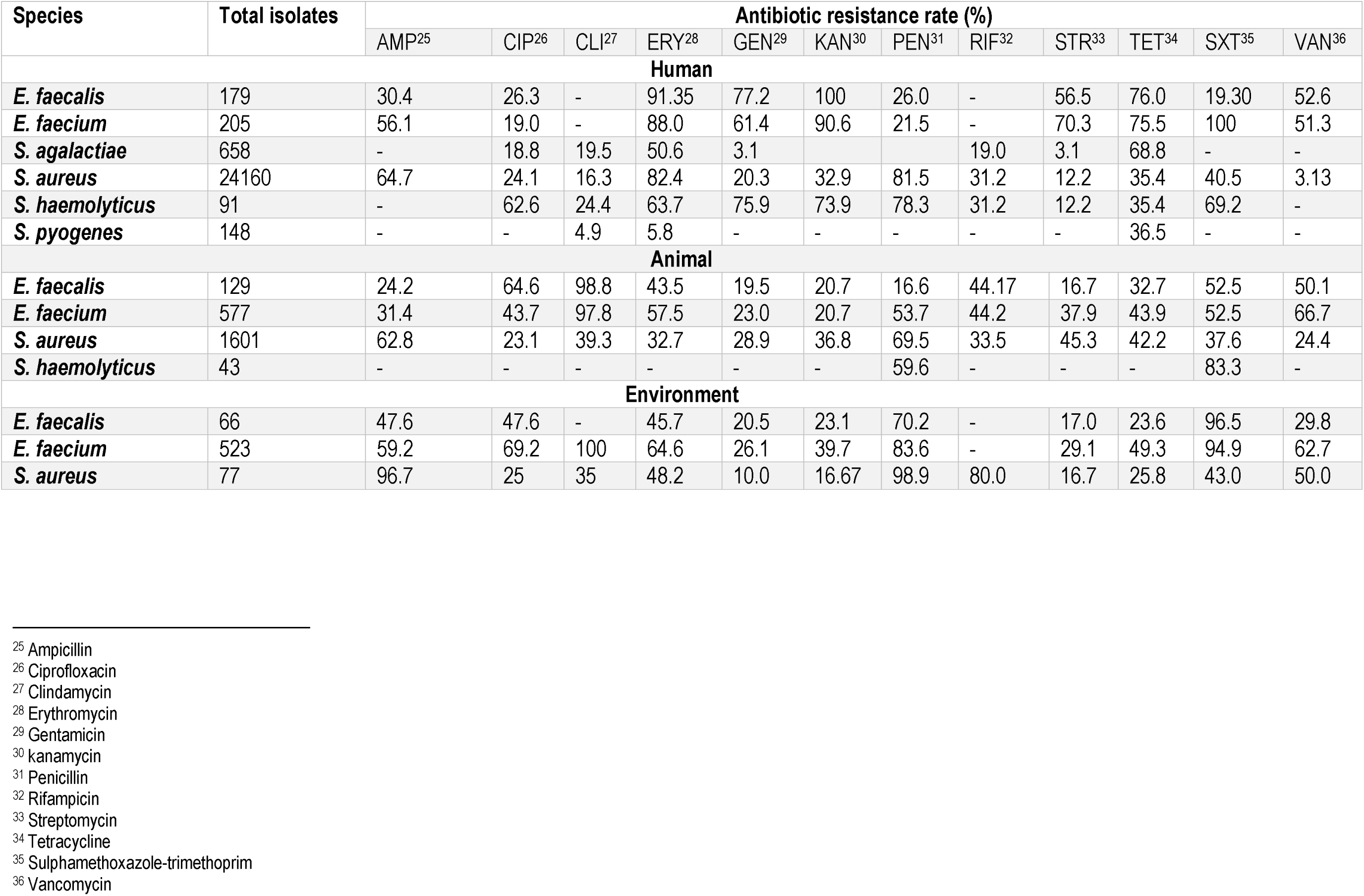
Antibiotic resistance rates of various Gram-positive bacterial species isolated from humans, animals and the environment in Africa between 2007 and 2018.

| Species | Total isolates | Antibiotic resistance rate (%) |  |  |  |  |  |  |  |  |  |  |  |
| --- | --- | --- | --- | --- | --- | --- | --- | --- | --- | --- | --- | --- | --- |
|  |  | AMP<sup>25</sup> | CIP<sup>26</sup> | CLI<sup>27</sup> | ERY<sup>28</sup> | GEN<sup>29</sup> | KAN<sup>30</sup> | PEN<sup>31</sup> | RIF<sup>32</sup> | STR<sup>33</sup> | TET<sup>34</sup> | SXT<sup>35</sup> | VAN<sup>36</sup> |
| Human |  |  |  |  |  |  |  |  |  |  |  |  |  |
| *E. faecalis* | 179 | 30.4 | 26.3 | - | 91.35 | 77.2 | 100 | 26.0 | - | 56.5 | 76.0 | 19.30 | 52.6 |
| *E. faecium* | 205 | 56.1 | 19.0 | - | 88.0 | 61.4 | 90.6 | 21.5 | - | 70.3 | 75.5 | 100 | 51.3 |
| *S. agalactiae* | 658 | - | 18.8 | 19.5 | 50.6 | 3.1 |  |  | 19.0 | 3.1 | 68.8 | - | - |
| *S. aureus* | 24160 | 64.7 | 24.1 | 16.3 | 82.4 | 20.3 | 32.9 | 81.5 | 31.2 | 12.2 | 35.4 | 40.5 | 3.13 |
| *S. haemolyticus* | 91 | - | 62.6 | 24.4 | 63.7 | 75.9 | 73.9 | 78.3 | 31.2 | 12.2 | 35.4 | 69.2 | - |
| *S. pyogenes* | 148 | - | - | 4.9 | 5.8 | - | - | - | - | - | 36.5 | - | - |
| Animal |  |  |  |  |  |  |  |  |  |  |  |  |  |
| *E. faecalis* | 129 | 24.2 | 64.6 | 98.8 | 43.5 | 19.5 | 20.7 | 16.6 | 44.17 | 16.7 | 32.7 | 52.5 | 50.1 |
| *E. faecium* | 577 | 31.4 | 43.7 | 97.8 | 57.5 | 23.0 | 20.7 | 53.7 | 44.2 | 37.9 | 43.9 | 52.5 | 66.7 |
| *S. aureus* | 1601 | 62.8 | 23.1 | 39.3 | 32.7 | 28.9 | 36.8 | 69.5 | 33.5 | 45.3 | 42.2 | 37.6 | 24.4 |
| *S. haemolyticus* | 43 | - | - | - | - | - | - | 59.6 | - | - | - | 83.3 | - |
| Environment |  |  |  |  |  |  |  |  |  |  |  |  |  |
| *E. faecalis* | 66 | 47.6 | 47.6 | - | 45.7 | 20.5 | 23.1 | 70.2 | - | 17.0 | 23.6 | 96.5 | 29.8 |
| *E. faecium* | 523 | 59.2 | 69.2 | 100 | 64.6 | 26.1 | 39.7 | 83.6 | - | 29.1 | 49.3 | 94.9 | 62.7 |
| *S. aureus* | 77 | 96.7 | 25 | 35 | 48.2 | 10.0 | 16.67 | 98.9 | 80.0 | 16.7 | 25.8 | 43.0 | 50.0 |

<sup>25</sup> Ampicillin <sup>26</sup> Ciprofloxacin <sup>27</sup> Clindamycin <sup>28</sup> Erythromycin <sup>29</sup> Gentamicin <sup>30</sup> kanamycin <sup>31</sup> Penicillin <sup>32</sup> Rifampicin <sup>33</sup> Streptomycin <sup>34</sup> Tetracycline <sup>35</sup> Sulphamethoxazole-trimethoprim <sup>36</sup> Vancomycin

Figure 1. PRISMA-adapted flow chart showing included and excluded articles. All search were conducted on PubMed, Web of Science and African Journals Online, and a final number of 130 articles were used for the quantitative analysis.

**Figure 2.**
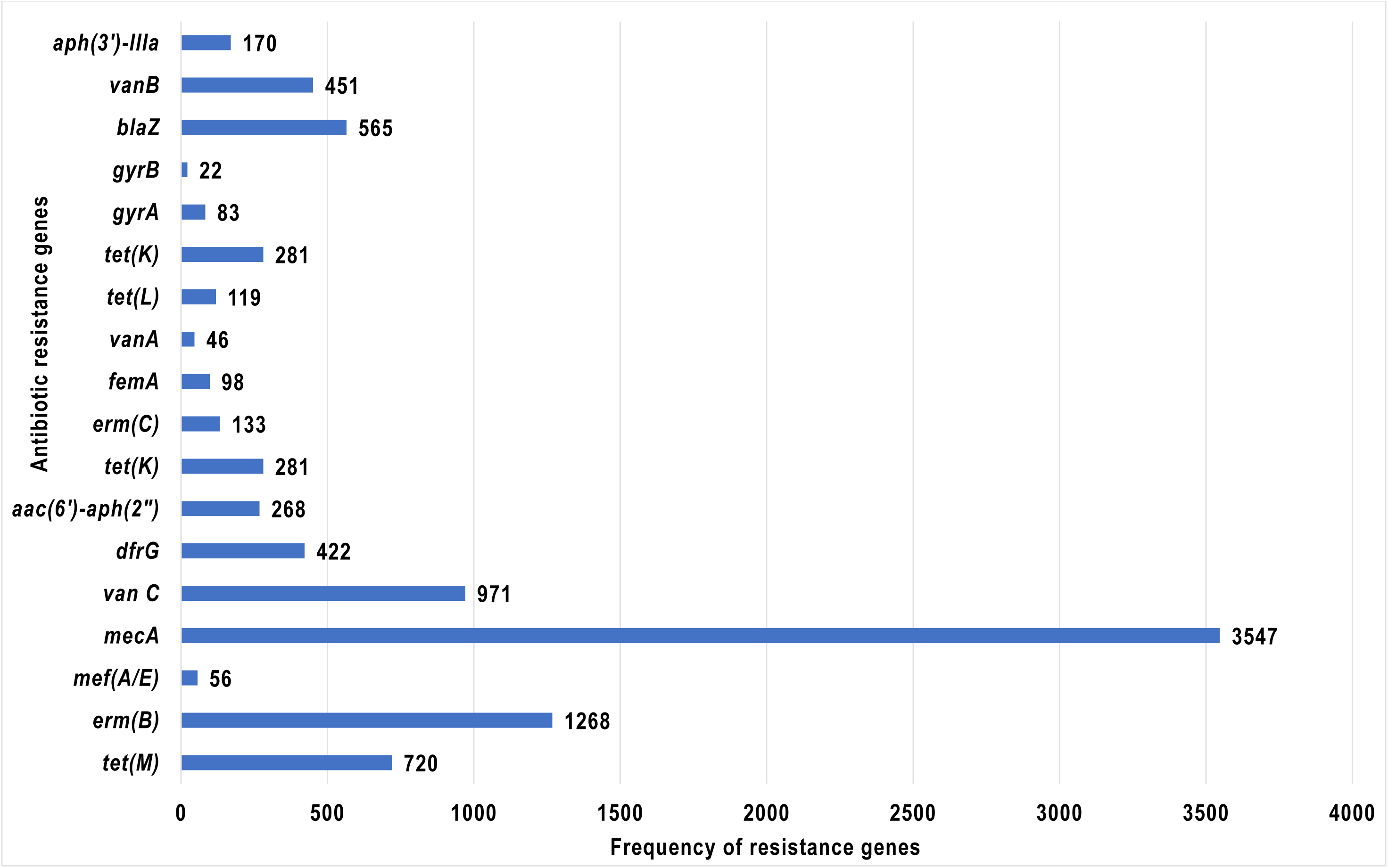
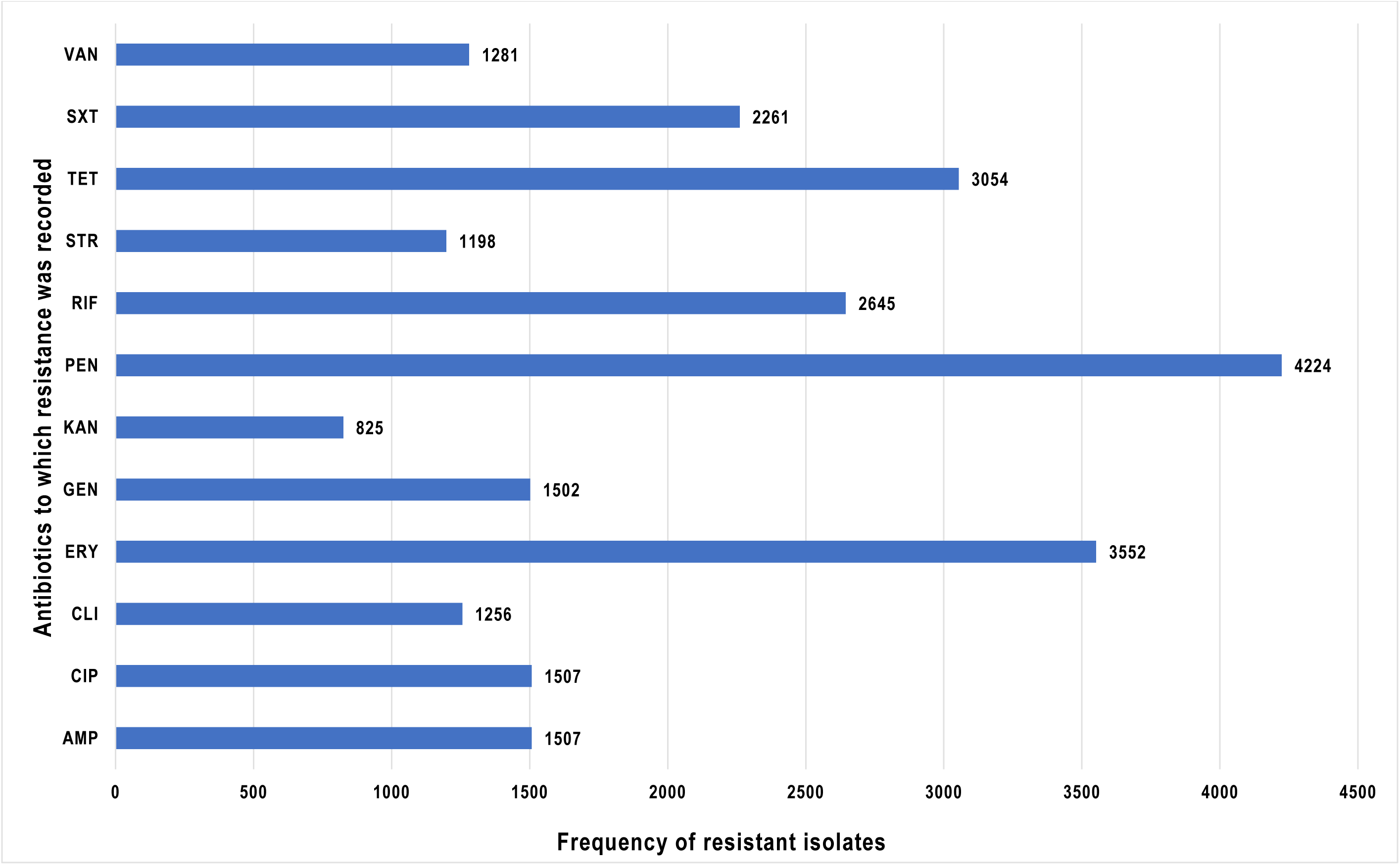
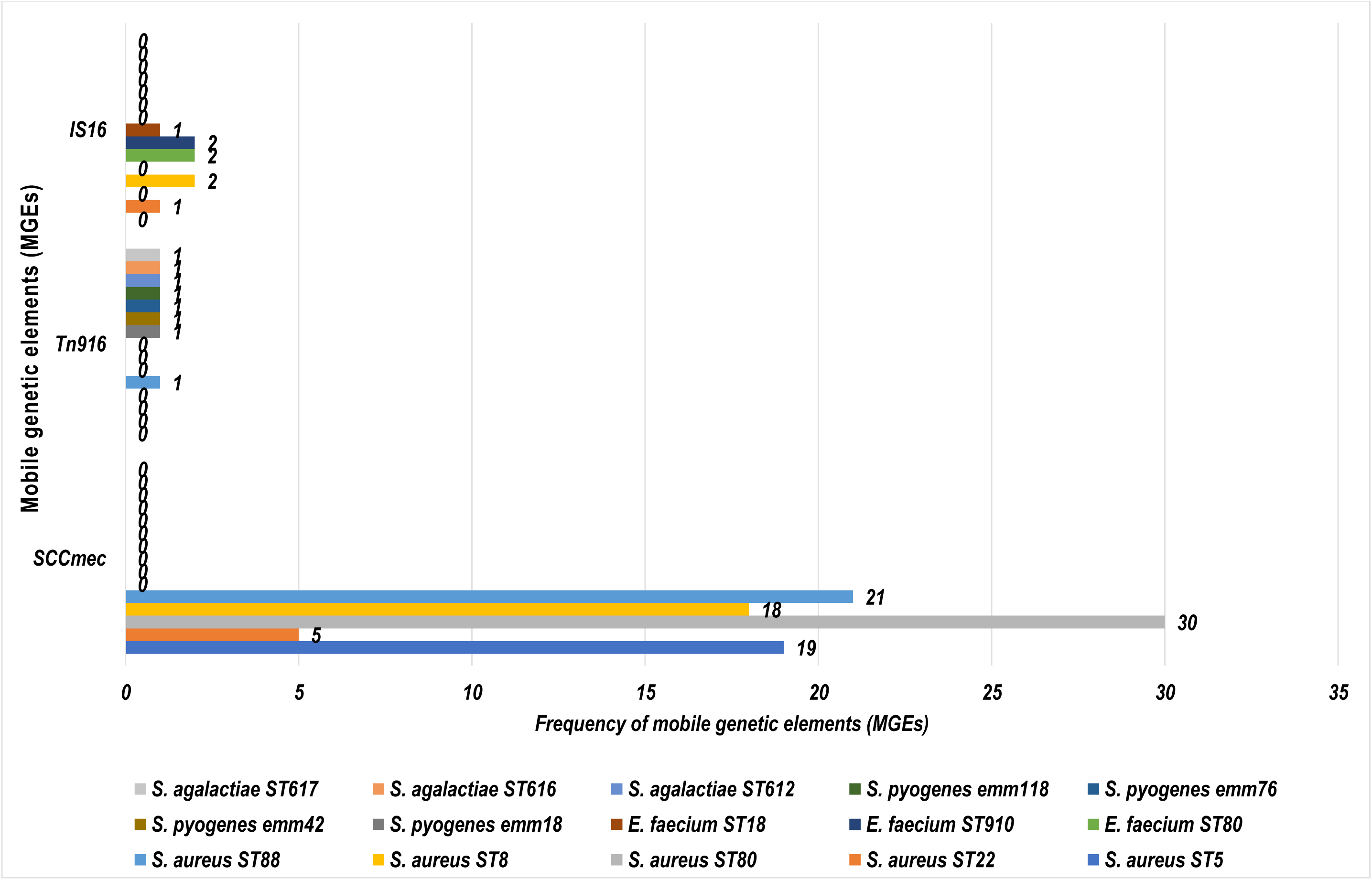
Frequency and distribution of resistance genes, antibiotics, and mobile genetic elements (MGEs) with recorded resistance in Gram-positive bacteria in Africa. 2ai) Shows the frequency of the various resistance genes found in the drug-resistant Gram-Positive bacterial strains. *mecA* and *erm*(B) were the most dominant resistance genes detected, followed by tet(M), *dfrG, vanB, vanC1* etc. 2aii) Shows the antibiotics to which the isolates were most resistant: erythromycin (ERY) was the least effective drug, followed by rifampicin (RIF), tetracycline (TET), penicillin (PEN), sulphamethoxazole/trimethoprim (SXT), ciprofloxacin (CIP), gentamicin (GEN), vancomycin (VAN), ampicillin (AMP), clindamycin (CLI), streptomycin (STR), chloramphenicol (CHL), and kanamycin (KAN). 2b) Shows the MGEs per resistant Gram-positive bacterial clones in Africa. The figure represents resistant clones and the different MGEs they carry. Each colour represent a particular resistant clone. *S. agalactiae* (ST612, ST616, ST617) and *S. pyogenes* (emm18, *emm42, emm76,* emm118), *E. faecium* (ST18, ST80, ST910) and *S. aureus* (ST5, ST22,ST35) were associated with *Tn916,* IS16 and *SCCmec* respectively.

**Figure 3.**
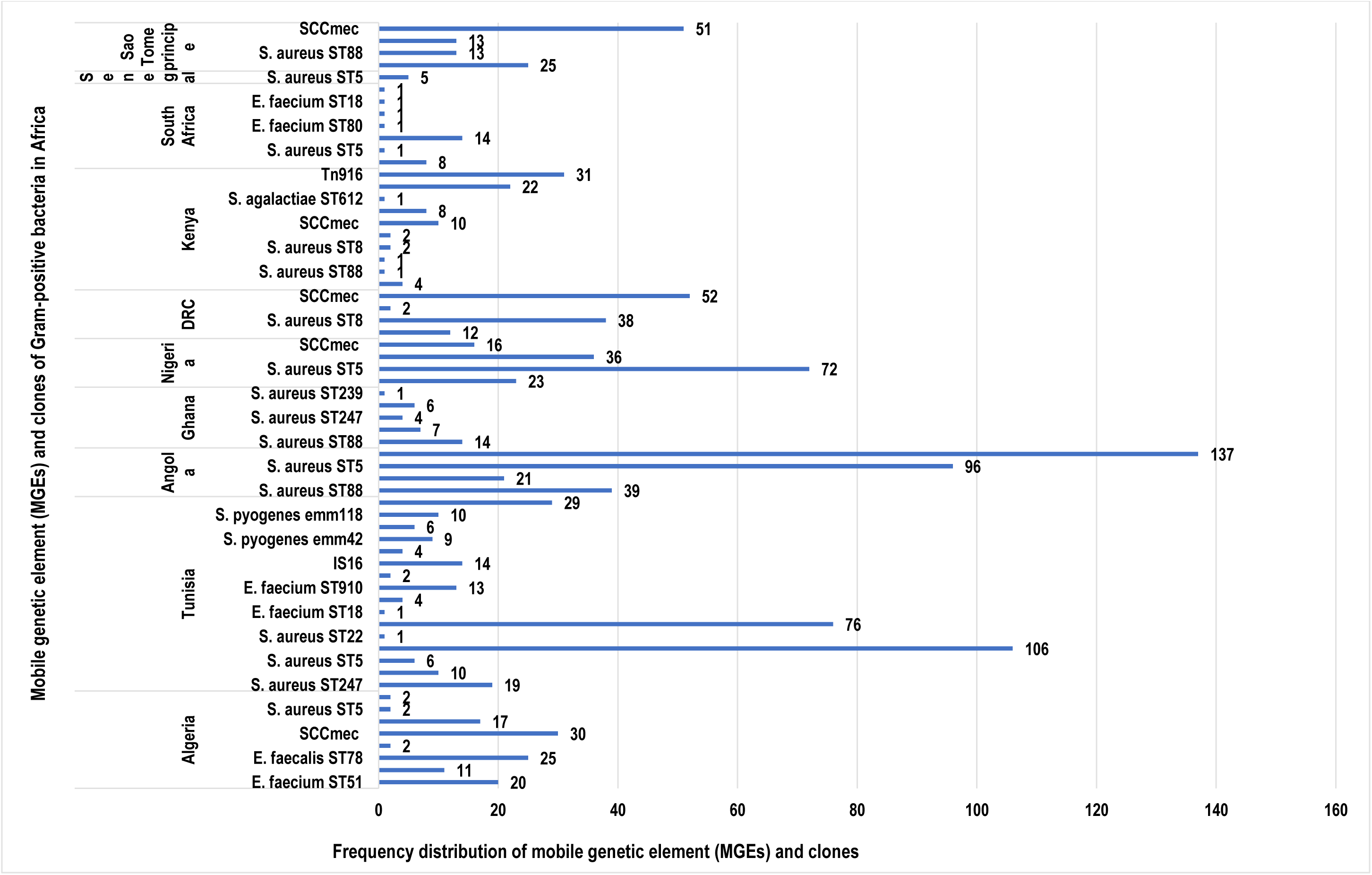
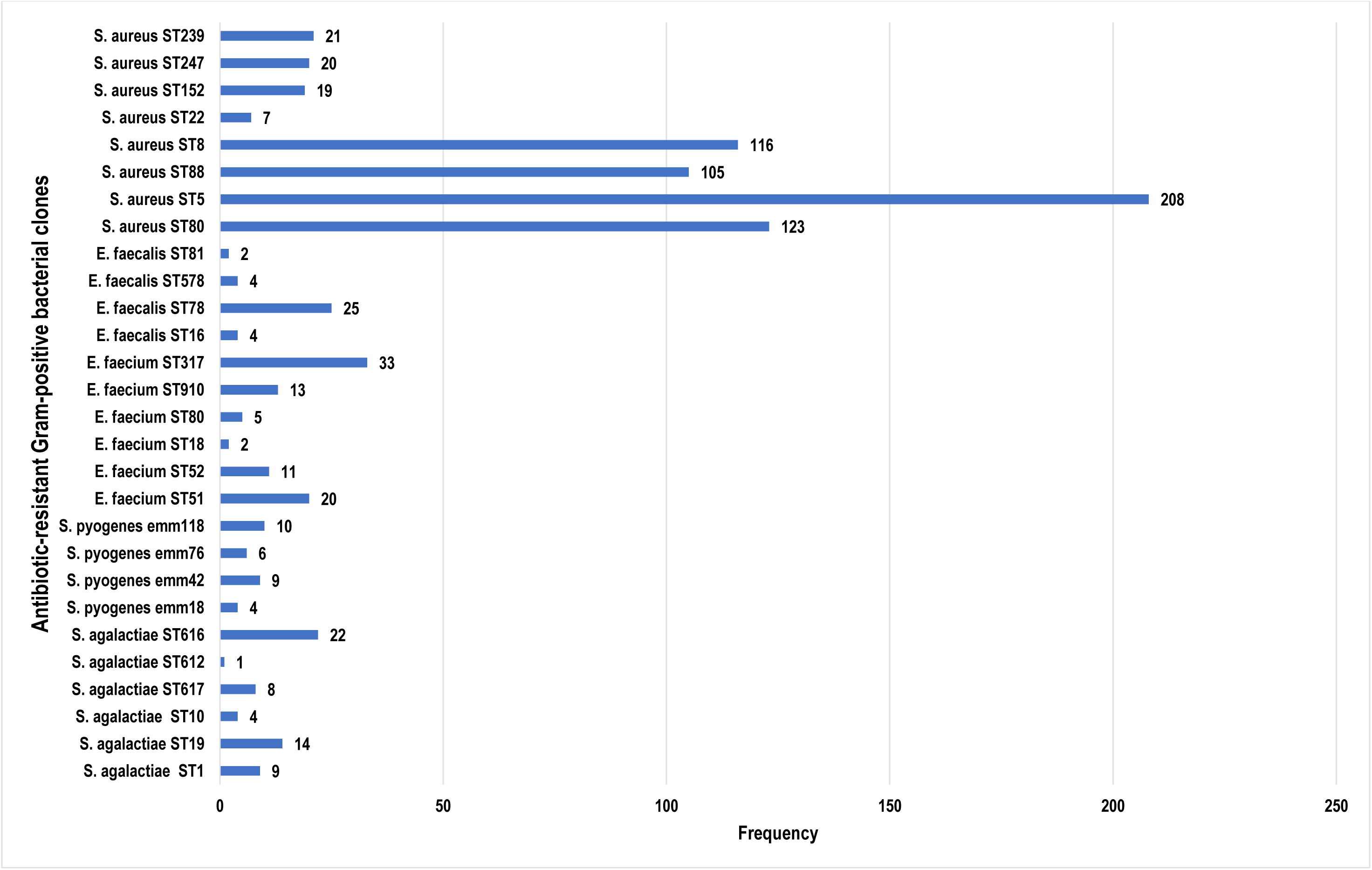
Frequency distribution of resistant Gram-positive bacterial species, clones and mobile genetic elements (MGEs) per country in Africa. 3a) Shows the distribution frequencies of the resistant species, clones and MGEs per country in Africa whilst 3b) shows the total frequency per clone in Africa. It is obvious that *S. aureus* ST5 is predominant in Tunisia, the DRC and Senegal whilst ST22 is highly prevalent in Algeria. *SCCmec* was the commonest MGE in most of the countries except in Tunisia where IS16 and *Tn916* were higher in prevalence. *S. aureus* ST8 and ST80 were the most common clones reported, followed by *E. faecium* ST317.

## Supplementary data 1

List of excluded articles on the basis of only phenotypic (antibiotic sensitivity) tests.

## Supplementary data 2

Raw data and analysis of extracted information from included articles.

## References

1. World Health Organization. Antimicrobial resistance—global report on surveillance. Geneva, Switzerland. WHO 2014.

2. Frean J, Perovic O, Fensham V, McCarthy K, von Gottberg A de GL et al. External quality assessment of national public health laboratories in Africa, 2002-2009. Bull WHO 2012; 90: 191-199A. Available at: doi:10.2471/BLT.11.091876.

3. Osei Sekyere J. Current State of Resistance to Antibiotics of Last-Resort in South Africa: a Review From a Public Health Perspective. Front Public Heal 2016; 4: 209. Available at: https://doi.org/10.3389/fpubh.2016.00209.

4. Williams PC, Isaacs D BJ. Antimicrobial resistance among children in sub-Saharan Africa. Lancet Infect Dis 2017; 18(2): Pages e33-e44. Available at: https://doi.org/10.1016/S1473-3099(17)30467-X.

5. Liu Y-Y, Wang Y, Walsh TR. Yi LX, Zhang R, Spencer J et al. Emergence of plasmidmediated colistin resistance mechanism MCR-1 in animals and human beings in China: a microbiological and molecular biological study. Lancet Infect Dis 2016; 16: 161-8. Available at: doi:10.1016/S1473-3099 (15)00424-7.

6. World Health Organization. WHO’s first global report on antibiotic resistance reveals serious, worldwide threat to public health. In Antimicrobial resistance–global surveillance report. Virtual Press Conference 2014 Apr. WHO 2014; (Vol. 30). Available at: http://www.who.int/mediacentre/news/releases/2014/amr-report/en/.

7. Witte W. ‘Antibiotic resistance in gram-positive bacteria: epidemiological aspects.’ J Antimicrob Chemother 1999; 44, su: 1-9. Available at:https://doi.org/10.1093/jac/44.suppl_1.1.

8. Seale AC, Mwaniki M, Newton CR BJ. Maternal and early onset neonatal bacterial sepsis: burden and strategies for prevention in sub-Saharan Africa. Lancet Infect Dis 2009; 9(7): 428-38. Available at: https://doi.org/10.1016/S1473-3099(09)70172-0.

9. Ateba CN, Lekoma KP KD. ‘Detection of vanA and vanB genes in vancomycin-resistant enterococci (VRE) from groundwater using multiplex PCR analysis.’ J Water Health 2013; 11.4: 684-691.

10. Iweriebor, Benson Chuks, Sisipho Gaqavu, Larry Chikwelu Obi, Uchechukwu U. Nwodo and AIO. ‘Antibiotic susceptibilities of Enterococcus species isolated from hospital and domestic wastewater effluents in Alice, eastern Cape Province of South Africa.’ 4231-4246. Int J Environ Res Public Health 12, no. 4. Available at: https://doi.org/10.3390/ijerph120404231.

11. Akanbi OE, Njom HA, Fri J, Otigbu AC CA. Antimicrobial Susceptibility of Staphylococcus aureus Isolated from Recreational Waters and Beach Sand in Eastern Cape Province of South Africa. Int J Environ Res public Heal 2017; 1;14(9): 1001. Available at:doi:10.3390/ijerph14091001.

12. Al-ashmawy MA, Sallam KI, Abd-elghany SM. Prevalence, Molecular Characterization, and Antimicrobial Susceptibility of Methicillin-Resistant Staphylococcus aureus Isolated from Milk and Dairy Products. FOODBORNE Pathog Dis 2016; 13: 156-62. Available at: https://doi.org/10.1089/fpd.2015.2038.

13. Iweriebor BC, Obi LC OA. Virulence and antimicrobial resistance factors of Enterococcus spp. isolated from fecal samples from piggery farms in Eastern Cape, South Africa. BMC Microbiol 2015; 15(1): 136. Available at: https://doi.org/10.1186/s12866-015-0468-7.

14. Katakweba AA, Møller KS, Muumba J, Muhairwa AP, Damborg P, Rosenkrantz JT, Minga UM, Mtambo MM OJ. Antimicrobial resistance in faecal samples from buffalo, wildebeest and zebra grazing together with and without cattle in Tanzania. J Appl Microbiol 2015; 118(4): 966-75. Available at: https://doi.org/10.1111/jam.12738%0A.

15. Osei Sekyere J. Types and selling practices of antibiotics in veterinary shops in ashanti region, GHANA. Int J Food, Agric Vet Sci 2014; 4(2): 87-96. Available at:http://www.cibtech.org/jfav.htm.

16. Osei Sekyere, John and FA. ‘Prevalence of Multidrug Resistance among Salmonella enterica Serovar Typhimurium Isolated from Pig Faeces in Ashanti Region, Ghana.’ Int J Antibiot 2015. Available at: http://dx.doi.org/10.1155/2015/898790.

17. Osei Sekyere J. ‘Antibiotic types and handling practices in disease management among pig farms in Ashanti Region, Ghana.’ J Vet Med 2014. Available at:http://dx.doi.org/10.1155/2014/531952.

18. Said MB, Abbassi MS, Gómez P, Ruiz-Ripa L, Sghaier S, Ibrahim C, Torres C HA. Staphylococcus aureus isolated from wastewater treatment plants in Tunisia: occurrence of human and animal associated lineages. J water Heal 2017; 15(4): 638-43.

19. Ben Said L, Klibi N, Dziri R, Borgo F, Boudabous A, Ben Slama K TC. ‘Prevalence, antimicrobial resistance and genetic lineages of Enterococcus spp. from vegetable food, soil and irrigation water in farm environments in Tunisia.’ J Sci Food Agric 2016; 96(5): 1627-33. Available at: https://doi.org/10.1002/jsfa.7264%0A.

20. Dziri R, Lozano C, Said LB, Bellaaj R, Boudabous A, Slama KB, Torres C KN. Multidrug-resistant enterococci in the hospital environment: detection of novel vancomycin-resistant E. faecium clone ST910. J Infect Dev Countries 2016; 10(08): 799-806. Available at: doi: https://doi.org/10.3855/jidc.8014.

21. Akindolire MA, Babalola OO AC. ‘Detection of antibiotic resistant Staphylococcus aureus from milk: A public health implication.’ Int J Environ Res Public Health 2015; 12, no. 9: 10254-10275. Available at: https://doi.org/10.3390/ijerph120910254.

22. Ben Said L, Dziri R, Sassi N, Lozano C, Ben Slama K, Ouzari I, Torres C KN. Species distribution, antibiotic resistance and virulence traits in canine and feline enterococci in Tunisia. Acta Vet Hungarica 2017; 65(2): 173-84. Available at: https://doi.org/10.1556/004.2017.018.

23. El-Hamid MI BM. ‘Comparative phenotypic and genotypic discrimination of methicillin resistant and susceptible Staphylococcus aureus in Egypt.’ Cell Mol Biol 2015; 61, no. 4: 101-112. Available at: http://www.cellmolbiol.com.

24. Mariem BJ, Ito T, Zhang M, et al. Molecular characterization of methicillin-resistant Panton-valentine leukocidin positive staphylococcus aureus clones disseminating in Tunisian hospitals and in the community. BMC 2013; 12: 2. Available at: https://doi.org/10.1186/1471-2180-13-2.

25. Ngbede EO, Raji MA, Kwanashie CN, Kwaga JK, Adikwu AA, Maurice NA AA. Characterization of high level ampicillin-and aminoglycoside-resistant enterococci isolated from non-hospital sources. J Med Microbiol 2017; 10;66(7): 1027-32. Available at: doi: 10.1099/jmm.0.000518.

26. Iweriebor BC, Obi LC OA. Virulence and antimicrobial resistance factors of Enterococcus spp. isolated from fecal samples from piggery farms in Eastern Cape, South Africa. BMC Microbiol 2015; 15: 136. Available at: https://doi.org/10.1186/s12866-015-0468-7.

27. Oumokhtar B, Elazhari M, Timinouni M, Bendahhou K, Bennani B, Mahmoud M, El Ouali Lalami A, Berrada S, Arrayhani M SHT. Staphylococcus aureus nasal carriage in a Moroccan dialysis center and isolates characterization. Hemodial Int 2013; 2: 542-7. Available at: https://doi.org/10.1111/j.1542-4758.2012.00759.x.

28. El Bayomi RM, Ahmed HA, Awadallah MA, Mohsen RA, Abd El-Ghafar AE AM. Occurrence, Virulence Factors, Antimicrobial Resistance, and Genotyping of Staphylococcus aureus Strains Isolated from Chicken Products and Humans. VECTOR-BORNE ZOONOTIC Dis 2016; 16: 157-64. Available at: https://doi.org/10.1089/vbz.2015.1891.

29. Hashem RA, Yassin AS, Zedan HH, Amin MA. Fluoroquinolone resistant mechanisms in methicillin-resistant Staphylococcus aureus clinical isolates in Cairo, Egypt. J Infect Dev Ctries 2013; 7, no. 11: 796-803. Available at: https://doi.org/10.3855/jidc.3105.

30. Lochan H, Moodley C, Rip D, Bamford C, Hendricks M, Davidson A EB. ‘Emergence of vancomycin-resistant Enterococcus at a tertiary paediatric hospital in South Africa.’ SAMJ South African Med J 2016; 106, no. 6: 562-566. Available at: 10.7196/SAMJ.2016.v106i6.10858.

31. Said LB, Klibi N, Lozano C, Dziri R, Slama KB, Boudabous A TC. ‘Diversity of enterococcal species and characterization of high-level aminoglycoside resistant enterococci of samples of wastewater and surface water in Tunisia.’. Sci Total Environ 2015; 530: 11-7. Available at: https://doi.org/10.1016/j.scitotenv.2015.05.091.

32. Shabayek S, Abdalla S. Macrolide- and tetracycline-resistance determinants of colonizing group B streptococcus in women in Egypt. J Med Microbiol 2014; 63: 1324-7. Available at: doi: 10.1099/jmm.0.077057-0.

33. Mbelle NM, Maningi NE, Tshisevhe V, Modipane L, Amoako DG, Osei Sekyere J. Draft Genome Sequence of a Clinical Enterococcus faecium Sequence Type 18 Strain from South Africa. Genome Announc 2017; 5: e01381-17. Available at:https://doi.org/10.1128/genomeA.01381-17.

34. Malachowa N DF. ‘Mobile genetic elements of Staphylococcus aureus.’ Cell Mol life Sci 2010; 67.18: 3057-3071. Available at: DOI 10.1007/s00018-010-0389-4.

35. Roberts AP MP. ‘Tn916-like genetic elements: a diverse group of modular mobile elements conferring antibiotic resistance.’ FEMS Microbiol Rev 2011; 35, no. 5: 856-871. Available at: https://doi.org/10.1111/j.1574-6976.2011.00283.x.

36. Frost LS, Leplae R, Summers AO TA. ‘Mobile genetic elements: the agents of open source evolution.’ Nat Rev Microbiol 2005; 3, no. 9: 722-732. Available at: doi:10.1038/nrmicro1235.

37. Thomas CM NK. ‘Mechanisms of, and barriers to, horizontal gene transfer between bacteria.’ (2005):. Nat Rev Microbiol 2005; 3, no. 9:711-21. Available at:doi: 10.1038/nrmicro1234.

38. Smillie C, Garcillán-Barcia MP, Francia MV, Rocha EP de la CF. ‘Mobility of plasmids.’ Microbiol Mol Biol Rev 2010; 74, no. 3: 434-452.

39. Bernabé KJ, Langendorf C, Ford N, Ronat JB MR. Antimicrobial resistance in West Africa: a systematic review and meta-analysis. Int J Antimicrob agents 2017; 50(5):629-39. Available at: https://doi.org/10.1016/j.ijantimicag.2017.07.002.

40. Tadesse BT, Ashley EA, Ongarello S, et al. Antimicrobial resistance in Africa□: asystematic review. BMC Infect Dis 2017; 17:616: 1-17. Available at:https://doi.org/10.1186/s12879-017-2713-1.

41. Workneh M, Katz MJ, Lamorde M, Cosgrove SE MY. Antimicrobial Resistance of Sterile Site Infections in Sub-Saharan Africa: A Systematic Review. Open forum Infect Dis: ofx209). US: Oxford University Press. Available at: https://doi.org/10.1093/ofid/ofx209.

42. Schaumburg F, Alabi AS, Peters G BK. New epidemiology of Staphylococcus aureus infection in Africa. Clinical. Microbiol Infect 2014; 20(7): 589-96. Available at: https://doi.org/10.1111/1469-0691.12690%0A.

43. Nejad SB, Allegranzi B, Syed SB, Ellis B PD. Health-care-associated infection in Africa: asystematic review. Bull World Heal Organ 2011; 89(10): 757-65. Available at:doi:10.2471/BLT.11.088179.

44. Abdulgader SM, Shittu AO, Nicol MP KM. Molecular epidemiology of Methicillin-resistant Staphylococcus aureus in Africa: a systematic review. Front Microbiol 2015; 6: 348. Available at: https://doi.org/10.3389/fmicb.2015.00348.

45. Bergal A, Loucif L, Benouareth DE, Bentorki AA, Abat C RJ. Molecular epidemiology and distribution of serotypes, genotypes, and antibiotic resistance genes of Streptococcus agalactiae clinical isolates from Guelma, Algeria and Marseille, France. European Journal of Clinical Microbiology & Infectious Disea. Eur J Clin Microbiol Infect Dis 2015; 34, no. 12: 2339-48. Available at: DOI 10.1007/s10096-015-2487-6.

46. Djoudi F, Benallaoua S, Aleo A, Touati A, Challal M, Bonura C MC. Descriptive Epidemiology of Nasal Carriage of Staphylococcus aureus and Methicillin-Resistant Staphylococcus aureus Among Patients Admitted at Two Healthcare Facilities in Algeria. Microb Drug Resist 2014; 21(2): 218-23. Available at: https://doi.org/10.1089/mdr.2014.0156.

47. Ouchenane Z, Agabou A, Smati F, Rolain JM RD. Spread of methicillin-resistant Staphylococcus aureus between the community and the hospitals in Asian countries: an ANSORP study. Journal of antimicrobial chemotherapy. Pathol Biol 2013; 61: 280-1. Available at: https://doi.org/10.1016/j.patbio.2013.05.006.

48. Djahmi N, Boutet-Dubois A, Nedjai S, Dekhil M, Sotto A LJ. Molecular epidemiology of Enterococcus sp. isolated in a university hospital in Algeria. Scand J Infect Dis 2012; 2011: 656-62. Available at: https://doi.org/10.3109/00365548.2012.673232.

49. Conceição T, Coelho C, de Lencastre H A-SM. Frequent occurrence of oxacillin-susceptible mecA -positive Staphylococcus aureus (OS-MRSA) strains in two African countries ‘. J Antimicrob Chemother 2015; 2015; 70: 3200-4. Available athttps://doi.org/10.1093/jac/dkv261.

50. Elhani D, Gharsa H, Kalai D, Lozano C, Gómez P, Boutheina J, Aouni M, Barguellil F, Torres C SK. ‘Clonal lineages detected amongst tetracycline-resistant meticillin-resistant Staphylococcus aureus isolates of a Tunisian hospital, with detection of lineage ST398.’ J Med Microbiol 2015; 64, no. 6: 623-629. Available at: doi: 10.1099/jmm.0.000066.

51. Klibi N, Said LB, Jouini A, Slama KB, López M, Sallem RB, Boudabous A TC. ‘Species distribution, antibiotic resistance and virulence traits in enterococci from meat in Tunisia.’ (2013):. Meat Sci 2013; 93, no. 3:675-80.Available at: http://doi.org/10.1016j.meatsci.2012.11.020.

52. Klibi N, Lagha AB, Slama KB, Boudabous A TC. ‘Faecal enterococci from camels in Tunisia: species, antibiotic resistance and virulent genes.’ Vet Rec 2013; 172,: 213-213. Available at: doi: 10.1136/vr. 100910.

53. Abdel moein KA. El Hariri M SA. Methicillin-Resistant Staphylococcus aureus_: An Emerging Pathogen of Pets in Egypt with a Public Health Burden. Transbound Emerg Dis 2012; 59: 331-5. Available at: https://doi.org/10.1111/j.1865-1682.2011.01273.x.

54. Fowoyo PT, Ogunbanwo ST. Antimicrobial resistance in coagulase negative staphylococci from Nigerian traditional fermented foods. Ann Clin Microbiol Antimicrob 2017; 16:4: 1-7. Available at: https://doi.org/10.1186/s12941-017-0181-5.

55. Nurjadi D, Olalekan AO, Layer F, Shittu AO, Alabi A, Ghebremedhin B, Schaumburg F, Hofmann-Eifler J, Van Genderen PJ, Caumes E FR. Emergence of trimethoprim resistance gene dfrG in Staphylococcus aureus causing human infection and colonization in sub-Saharan Africa and its import to Europe. J Antimicrob Chemother 2014; 27: 2361-8. Available at: https://doi.org/10.1093/jac/dku174.

56. O’Malley SM, Emele FE, Nwaokorie FO, Idika N, Umeizudike AK, Emeka-Nwabunnia I,Hanson BM, Nair R, Wardyn SE ST. Molecular typing of antibiotic-resistant Staphylococcus aureus in Nigeria. J Infect Public Health 2015; 8: 187-93. Available at:http://dx.doi.org/10.1016/jjiph.2014.08.001.

57. Shittu A, Oyedara O, Abegunrin F, Okon K, Raji A, Taiwo S, Ogunsola F, Onyedibe K EG. Characterization of methicillin-susceptible and -resistant staphylococci in the clinical setting: a multicentre study in Nigeria. BMC Infect Dis 2012; 12(1): 286. Available at: https://doi.org/10.1016/S0732-8893(98)00118-7.

58. Vitali LA, Petrelli D, Lamikanra A, Prenna M, Akinkunmi EO. Diversity of antibiotic resistance genes and staphylococcal cassette chromosome mec elements in faecal isolates of coagulase-negative staphylococci from Nigeria. BMC Microbiol 2014; 14:106. Available at: https://doi.org/10.1186/1471-2180-14-106.

59. van Rensburg MJ, Whitelaw AC EB. Genetic basis of rifampicin resistance in methicillin-resistant Staphylococcus aureus suggests clonal expansion in hospitals in Cape. BMC Microbiol 2012; 12: 46. Available at: https://doi.org/10.1186/1471-2180-12-46.

60. Perovic O, Iyaloo S, Kularatne R, Lowman W. Prevalence and Trends of Staphylococcus aureus Bacteraemia in Hospitalized Patients in South Africa, 2010 to 2012□: Laboratory-Based Surveillance Mapping of Antimicrobial Resistance and Molecular Epidemiology. PLoS One 2015: 1-14.

61. Adegoke AA, Okoh AI. Species diversity and antibiotic resistance properties of Staphylococcus of farm animal origin in Nkonkobe Municipality, South Africa. FoliaMicrobiol 2014; 59: 133-40. Available at: DOI 10.1007/s12223-013-0275-1.

62. WHO. Tackling Antibiotic Resistance from a Food Safety Perspective in Europe. WHO-Europe, Denmark 2011: 1-88. Available at: http://www.euro.who.int/…/e94889.pdf.

63. O’Neill J. ‘Tackling drug-resistant infections globally: final report and recommendations.’ Rev Antimicrob Resist 2016.

64. Cavaco LM AF. ‘Resistance in bacteria of the food chain: epidemiology and control strategies.’ Microb Drug Resist Futur Med Ltd, Brastislava, Slovakia 2013: 136-158. Available at: https://doi.org/10.2217/ebo.12.361

65. Boeck H De, Vandendriessche S, Hallin M, Batoko B, Alworonga J. Staphylococcus aureus nasal carriage among healthcare workers in Kisangani, the Democratic Republic of the Congo. Eur J Clin Microbiol Infect Dis 2015; 34(8): 1567-72. Available at: DOI 10.1007/s10096-015-2387-9.

66. Fall C, Seck A, Richard V, Ndour M, Sembene M, Laurent F BS. Epidemiology of Staphylococcus aureus in Pigs and Farmers in the Largest Farm in Dakar, Senegal. FOOD BORNE Pathog Dis 2012; 9: 962-5. Available at: https://doi.org/10.1089/fpd.2012.1197.

67. Stefani S, Chung DR, Lindsay JA, Friedrich AW, Kearns AM, Westh H MF. ‘Meticillin-resistant Staphylococcus aureus (MRSA): global epidemiology and harmonisation of typing methods.’ Int J Antimicrob Agents 2012; 39, no. 4: 273-282. Available at: https://doi.org/10.1016/j.ijantimicag.2011.09.030.

68. Song JH, Hsueh PR, Chung DR, Ko KS, Kang CI, Peck KR et al. Spread of methicillin-resistant Staphylococcus aureus between the community and the hospitals in Asian. J Antimicrob Chemother 2011 2011; 66: 1061-9. Available at: https://doi.org/10.1093/jac/dkr024.

69. Elhani D, Klibi N, Dziri R, Hassan MB, Mohamed SA, Said LB, Mahjoub A, Slama KB, Jemli B, Bellaj R BF. ‘vanA-containing E. faecium isolates of clonal complex CC17 in clinical and environmental samples in a Tunisian hospital.’ Diagn Microbiol Infect Dis 2014; 79, no. 1: 60-63. Available at: http://dx.doi.org/10.1016/j.diagmicrobio.2014.01.011.

70. Dziri R, Lozano C, Said LB, Bellaaj R, Boudabous A, Slama KB, Torres C KN. ‘Multidrug-resistant enterococci in the hospital environment: detection of novel vancomycin-resistant E. faecium clone ST910.’ J Infect Dev Ctries 2016; 10, no. 8: 799-806. Available at: doi: https://doi.org/10.3855/jidc.8014.

71. Udo EE, Pearman JW GW. Genetic analysis of community isolates of methicillin-resistant Staphylococcus aureus in Western Australia. J Hosp Infect 1993; 25: 97-108. Available at: https://doi.org/10.1016/0195-6701(93)90100-E%0A.

72. Salgado CD, Farr BM CD. Community-acquired methicillin-resistant Staphylococcus aureus: a meta-analysis of prevalence and risk factors. Clin Infect Dis 2003, 36 2003; 36: 131-139. Available at: https://doi.org/10.1086/345436.

73. Hiramatsu K, Okuma K, Ma XX, Yamamoto M, Hori S et al: New trends in Staphylococcus aureus infections: glycopeptide resistance in hospital and methicillin resistance in the community. Curr Opin Infect Dis 2002, 15 2002;15: 407-413. Available at:doi:10.1097/00132980-200201060-0000.

74. Chambers H. The changing epidemiology of Staphylococcus aureus? Emerg Infect Dis 2001; 7: 178-182.

75. Diekema DJ, Pfaller MA, Schmitz FJ, Smayevsky J, Bell J et al. Survey of infections due to Staphylococcus species: frequency of occurrence and antimicrobial susceptibility of isolates collected in the United States, Canada, Latin America, Europe, and the Western Pacific region for the SENTRY Antimicrobial Surveillanc. Clin Infect Dis 2001; 32 Suppl 2: S114-132. Available at: https://doi.org/10.1086/320184.

76. NNIS. National Nosocomial Infections Surveillance (NNIS) System Report, data summary from January 1992 through June 2004. Am J Infect Control 2004; 2: 470-485.

77. Katayama Y, Ito T HK. A new class of genetic element, staphylococcus cassette chromosome mec, encodes methicillin resistance in Staphylococcus aureus. Antimicrob agents Chemother 2000; 44(6): 1549-55. Available at: doi: 10.1128/AAC.44.6.1549-1555.2000.

78. Akinkunmi E LA. Species distribution and antibiotic resistance in coagulase-negative staphylococci colonizing the gastrointestinal tract of children in Ile-Ife, Nigeria. Trop J Pharm Res 2010; 9(1): 35-43. Available at: http://dx.doi.org/10.4314/tjpr.v9i1.52033.

79. Archer GL CM. Antimicrobial susceptibility of coagulase-negative staphylococci. Antimicrob Agents Chemother 1994; 38: 2231-2237.

80. Hashem YA, Yassin AS AM. Molecular characterization of Enterococcus spp. clinical isolates from Cairo, Egypt. Indian J Med Microbiol 2015; 33(5): p.80. Available at: doi: 10.4103/0255-0857.148836.

81. Hraoui M, Boubaker IB, Doloy A, Redjeb SB BA. Molecular Mechanisms of Tetracycline and Macrolide Resistance and emm Characterization of Streptococcus pyogenes Isolates in Tunisia. Microb Drug Resist 2011; 17(3): 377-82. Available at:https://doi.org/10.1089/mdr.2010.0160.

82. Rachdi M, Boubaker IB, Moalla S, et al. Phenotypic and genotypic characterization of macrolide resistant Streptococcus pneumoniae in Tunisia. Pathol Biol 2008; 56: 125-9. Available at: https://doi.org/10.1016/j.patbio.2007.05.005.

83. Wolter N, Gottberg A Von, Gouveia L De, Klugman KP. Molecular basis and clonal nature of increasing pneumococcal macrolide resistance in South Africa, 2000 – 2005. Int J Antimicrob Agents 2008; 32: 2000-5. Available at: https://doi.org/10.1016/j.ijantimicag.2008.01.013.

84. Hancock RE. Mechanisms of action of newer antibiotics for Gram-positive pathogens. Lancet Infect Dis 2005; 5: 209-18. Available at: https://doi.org/10.1016/S1473-3099(05)70051-7.

85. Hraoui M, Boubaker IB, Rachdi M, Slim A RS. Macrolide and tetracycline resistance in clinical strains of Streptococcus agalactiae isolated in Tunisia. J Med Microbiol 2017; 61: 1109-13. Available at: doi: 10.1099/jmm.0.037853-0.

86. Fischer A, Liljander A, Kaspar H, Muriuki C, Fuxelius HH, Bongcam-Rudloff E, de Villiers EP, Huber CA, Frey J, Daubenberger C BR. Camel Streptococcus agalactiae populations are associated with specific disease complexes and acquired the tetracycline resistance gene tetM via a Tn 916 -like element Camel Streptococcus agalactiae populations are associated with specific disease complex. Vet Res 2013, 2013; 44:86. Available at: https://doi.org/10.1186/1297-9716-44-86.

87. Omuse G, Zyl KN Van, Hoek K, Abdulgader S, Kariuki S, Whitelaw A. Molecular characterization of Staphylococcus aureus isolates from various healthcare institutions in Nairobi, Kenya: a cross sectional study. Ann Clin Microbiol Antimicrob 2016; 15:51: 1-9. Available at: https://doi.org/10.1186/s12941-016-0171-z.

88. Stokes HW GM. ‘Gene flow, mobile genetic elements and the recruitment of antibiotic resistance genes into Gram-negative pathogens.’ FEMS Microbiol Rev 2011; 35, no. 5: 790-819. Available at: https://doi.org/10.1111/j.1574-6976.2011.00273.x.

89. Gogarten JP TJ. ‘Horizontal gene transfer, genome innovation and evolution.’ Nat Rev Microbiol 2005; 3, no. 9: 679-687. Available at: doi:10.1038/nrmicro1204.

90. Bouchami O, Hassen AB, De Lencastre H MM. High prevalence of mec complex C and ccrC is independent of SCC mec type V in Staphylococcus haemolyticus. Eur J Clin Microbiol Infect Dis 2012; 31: 605-14. Available at: 10.1007/s10096-011-1354-3.

91. Khalil W, Hashwa F, Shihabi A TS. Methicillin-resistant Staphylococcus aureus ST80-IV clone in children from Jordan. Diagn Microbiol Infect Dis 2012; 73: 228-230. Available at:https://doi.org/10.1016/j.diagmicrobio.2012.03.012%0AGet

92. Udo EE SE. The dissemination of ST80-SCCmec-IV community-associated methicillin resistant Staphylococcus aureus clone in Kuwait hospitals. Ann Clin Microbiol Antimicrob 2010; 9: 1–7. Available at: https://doi.org/10.1186/1476-0711-9-31.

93. Ahmed EF, Gad GF, Abdalla AM, Hasaneen AM AS. Prevalence of Methicillin Resistant Staphylococcus aureus among Egyptian Patients after Surgical interventions. Surg Infect (Larchmt) 2014; 15: 404-11. Available at: https://doi.org/10.1089/sur.2013.212.

94. Osman K, Alvarez-Ordóñez A, Ruiz L, Badr J, ElHofy F, Al-Maary KS, Moussa IM, Hessain AM, Orabi A, Saad A EM. Antimicrobial resistance and virulence characterization of Staphylococcus aureus and coagulase-negative staphylococci from imported beef meat. Ann Clin Microbiol Antimicrob 2017; 16(1): 35. Available at: https://doi.org/10.1186/s12941-017-0210-4.

95. Perveen I, Majid A, Knawal S, Naz I, Sehar S, Ahmed S RM. Biochemical characters and antibiotic susceptibility of S. aureus isolates. Asian Pac J Trop Biomed 2011; 1: 212-216. Available at: doi: 10.1016/S2221-1691(11)60029-4.

96. Perveen, I., Majid, A., Knawal, S., Naz, I., Sehar, S. A. S. and Raza MA. Prevalence and antimicrobial susceptibility pattern of methicillin-resistant Staphylococcus aureus and coagulasenegative staphylococci in Rawalpindi, Pakistan. Br J Med Med Res 2013; 3: 198-209. Available at: doi: 10.9734/%0ABJMMR/2013/2109.

97. Dziri R, Klibi N, Lozano C, et al. High prevalence of Staphylococcus haemolyticus and Staphylococcus sapro-phyticus in environmental samples of a Tunisian hospital. Diagn Microbiol Infect Dis 2016; 85.2: 136-140. Available at:http://dx.doi.org/10.1016/j.diagmicrobio.2016.03.006.

98. Stoll BJ, Hansen N, Fanaroff AA, Wright LL, Carlo WA ER, Lemons JA, Donovan EF,Stark AR, Tyson JE, Oh W BC, Korones SB, Shankaran S, Laptook AR, Stevenson DK, Papile LA P. WK. Late-onset sepsis in very low birth weight neonates: the experience of the NICHD Neonatal Research Network. J Pediatr 2002; 110: 285-291. Available at: doi:10.1542/peds.110.2.285.

99. Gharsa H, Slama KB, Lozano C, Gómez-Sanz E, Klibi N, Sallem RB, Gómez P, Zarazaga M, Boudabous A TC. Prevalence, antibiotic resistance, virulence traits and genetic lineages of Staphylococcus aureus in healthy sheep in Tunisia. Veterinary microbiol. Vet Microbiol 2012; 156(3): pp.367-l373. Available at: https://doi.org/10.1016/j.vetmic.2011.11.009.

100. Cui S, Li J, Hu C, Jin S, Li F, Guo Y, Ran LU MY. Isolation and characterization of methicillin-resistant Staphylococcus aureus from swine and workers in China. J Antimicrob Chemother 2009; 64: 680-683. Available at: https://doi.org/10.1093/jac/dkp275.

101. Jiménez JN, Vélez LA, Mediavilla JR, Ocampo AM, Vanegas JM, Rodríguez EA, Kreiswirth BN CM. Livestock-associated methicillin-susceptible Staphylococcus aureus ST398 infection in woman, Colombia. Emerg Infect Dis 2011; 17: 1970-1971. Available at: doi:10.3201/eid1710.110638.

102. Khanna T, Friendship R, Dewey C WJ. Methicillin resistant Staphylococcus aureus colonization in pigs and pig farmers. Vet Microbiol 2008; 128: 298-303. Available at: https://doi.org/10.1016/j.vetmic.2007.10.006.

103. Smith TC, Male MJ, Harper AL, Kroeger JS, Tinkler GP, Moritz ED, Capuano AW, Herwaldt LA DD. Methicillin-resistant Staphylococcus aureus (MRSA) strain ST398 is present in midwestern U.S. swine and swine workers. PLoS One 2009; 4,: e4258.

104. Maalej SM, Malbruny B, Leclercq R HA. Emergence of Staphylococcus aureus strains resistant to pristinamycin in Sfax (Tunisia). Pathol Biol 2012; 60: e71-4. Available at: http://dx.doi.org/10.1016/j.patbio.2011.10.012.

105. Bouchami O, Hassen AB, De Lencastre H MM. High prevalence of mec complex C and ccrC is independent of SCCmec type V in Staphylococcus haemolyticus. Eur J Clin Microbiol Infect Dis 2012; 31(4): 605-614. Available at: DOI 10.1007/s10096-011-1354-3.

106. Tabe Y, Nakamura A, Oguri T IJ. Molecular characterization of epidemic multiresistant Staphylococcus haemolyticus isolates. Diagn Microbiol Infect Dis 1998; 32: 177-183. Available at:https://doi.org/10.1016/S0732-8893(98)00118-7.

107. Santos Sanches I, Mato R, de Lencastre H, Tomasz A C, Collaborators C and the I. Patterns of multidrug resistance among methicillin-resistant hospital isolates of coagulase-positive and coagulase-negative staphylococci collected in the international multicenter study RESIST in 1997 and 1998. Microb Drug Resist 2000; 6: 199-211. Available at: https://doi.org/10.1089/mdr.2000.6.199.

108. Al-haddad OH, Zorgani A GK. Nasal Carriage of Multi-Drug Resistant Panton-Valentine Leucocidin-Positive Methicillin-Resistant Staphylococcus aureus in Children in Tripoli-Libya. 2014; 90: 724-7. Available at: doi: https://doi.org/10.4269/ajtmh.13-0715

109. Egyir B, Guardabassi L, Sørum M, Nielsen SS, Kolekang A, Frimpong E, Addo KK, Newman MJ LA. Molecular epidemiology and antimicrobial susceptibility of clinical Staphylococcus aureus from healthcare institutions in Ghana. PLoS One, 2014; 9(2): p.e89716.

110. Egyir B, Guardabassi L, Monecke S, Kwasi K, Jemima M, Rhod A. Resistance Methicillin-resistant Staphylococcus aureus strains from Ghana include USA300. J Glob Antimicrob Resist 2015; 3: 26-30. Available at: http://dx.doi.org/10.1016/jjgar.2014.11.006.

111. Bouchami O, Ben Hassen A, de Lencastre H MM. High prevalence of mec complex C and ccrC is independent of SCCmec type V in Staphylococcus haemolyticus. Eur J Clin Microbiol Infect Dis 2012; 31(4): 605-614. Available at: DOI 10.1007/s10096-011-1354-3.

112. Shittu A, Oyedara O, Abegunrin F, Okon K, Raji A, Taiwo S, Ogunsola F, Onyedibe K EG. ‘Characterization of methicillin-susceptible and-resistant staphylococci in the clinical setting: a multicentre study in Nigeria.’ BMC Infect Dis 12, no. 1: 286. Available at:https://doi.org/10.1186/1471-2334-12-286.

113. Naber KG, Schito G, Botto H, Palou J MT. Surveillance Study in Europe and Brazil on Clinical Aspects and Antimicrobial Resistance Epidemiology in Females with Cystitis (ARESC): Implications for Empiric Therapy. Eur Urol 2008; 54: 1164-1178. Available at: https://doi.org/10.1016/j.eururo.2008.05.010.

114. Conceição T, Coelho C, Silva IS, de Lencastre H A-SM. Staphylococcus aureus in former Portuguese colonies from Africa and the Far East: missing data to help fill the world map. Clin Microbiol Infect 2015; 21(9): 842-e1. Available at: https://doi.org/10.1016/j.cmi.2015.05.010.

115. Jensen SO LB. Genetics of antimicrobial resistance in Staphylococcus aureus. Futur Microbiol 2009; 4: 565-82. Available at: https://doi.org/10.2217/fmb.09.30.

116. Frey KM, Lombardo MN, Wright DL AA. Towards the understanding of resistance mechanisms in clinically isolated trimethoprim-resistant, methicillin-resistant Staphylococcus aureus dihydrofolate reductase. J Struct Biol 2010; 170: 93-7. Available at:https://doi.org/10.1016/jjsb.2009.12.011.

117. Kadlec K, Fessler AT, Hauschild T SS. Novel and uncommon antimicrobial resistance genes in livestock-associated methicillinresistant Staphylococcus aureus. Clin Microbiol Infect 2012; 18: 745-55. Available at: https://doi.org/10.1111/j.1469-0691.2012.03842.x.

118. Badri M, Ehrlich R, Wood R MG. Initiating co-trimoxazole prophylaxis in HIV-infected patients in Africa: an evaluation of the provisional WHO/UNAIDS recommendations.. AIDs 2001; 15(9): 1143-8. Available at: doi:10.1097/00002030-200106150-00009.

119. Phaku P, Lebughe M, Strauß L, et al. Unveiling the molecular basis of antimicrobial resistance in Staphylococcus aureus from the Democratic Republic of the Congo using whole genome sequencing. Clin Microbiol Infect 2016. Available at: http://dx.doi.org/10.1016/jxmi.2016.04.009.

120. Vandendriessche S, De Boeck H, Deplano A, Phoba MF, Lunguya O, Falay D, Dauly N, Verhaegen J, Denis O JJ. Characterisation of Staphylococcus aureus isolates from bloodstream infections, Democratic Republic of the Congo. Eur J Clin Microbiol Infect Dis 2017; 36(7): 1163-71. Available at: DOI 10.1007/s10096-017-2904-0.

121. Asiimwe BB, Baldan R, Trovato A CD. Prevalence and molecular characteristics of Staphylococcus aureus, including methicillin resistant strains, isolated from bulk can milk and raw milk products in pastoral communities of South-West Uganda. BMC Infect Dis 2017; 17(1): 422. Available at: https://doi.org/10.1186/s12879-017-2524-4.

122. Seni J, Bwanga F, Najjuka CF, Makobore P, Okee M, Mshana SE, Kidenya BR, Joloba ML KD. Molecular Characterization of Staphylococcus aureus from Patients with Surgical Site Infections at Mulago Hospital in Kampala, Uganda. PLoS One 2013; 8: 1-7.

123. Conceiçao T, Coelho C, Santos-Silva I, de Lencastre H A-SM. Epidemiology of methicillin-resistant and-susceptible Staphylococcus aureus in Luanda, Angola: first description of the spread of the MRSA ST5-IVa clone in the African continent. Microb Drug Resist 2014; 20(5): 441-9. Available at: https://doi.org/10.1089/mdr.2014.0007.

124. Conceição T, Coelho C, de Lencastre H A-SM. Frequent occurrence of oxacillin-susceptible mecA-positive Staphylococcus aureus (OS-MRSA) strains in two African countries. J Antimicrob Chemother 2015; 70(12): 3200-4. Available at: https://doi.org/10.1093/jac/dkv261.

125. Hososaka Y, Hanaki H EH et al. Characterization of oxacillinsusceptible mecA-positive Staphylococcus aureus: a new type of MRSA. J Infect Chemother 2007; 13: 79-86. Available at:https://doi.org/10.1007/s10156-006-0502-7.

126. Petinaki E, Kontos F MA. Emergence of two oxacillin-susceptible mecA-positive Staphylococcus aureus clones in a Greek hospital. J Antimicrob Chemother 2002; 50: 1090-1. Available at: https://doi.org/10.1093/jac/dkf235.

127. Pu W, Su Y LJ et al. High incidence of oxacillin-susceptible mecA-positive Staphylococcus aureus (OS-MRSA) associated with bovine mastitis in China. PLoS One 2014; 9: e88134.

128. Corrente M, Normanno G, Martella V, Bellacicco AL, Quaglia NC, Dambrosio A, Buonavoglia D, D’Abramo M BC. Comparison of methods for the detection of methicillin resistance in Staphylococcus aureus isolates from food products. Lett Appl Microbiol 2007; 45: 535-9. Availableat:https://doi.org/10.1111/j.1472-765X.2007.02226.x%0A.

129. Meeren BT, Millard PS, Scacchetti M, Hermans MH, Hilbink M, Concelho TB, Ferro JJ WP. Emergence of methicillin resistance and Panton-Valentine leukocidin positivity in hospital- and community-acquired Staphylococcus aureus infections in Beira, Mozambique. Trop Med Int Heal Vol 2013; 0. Available at: https://doi.org/10.1111/tmi.12221%0A.

130. De Angelis G, Cipriani M, Cauda R TE. Treatment of skin and soft tissue infections due to community-associated methicillin-resistant Staphylococcus aureus in Europe: the role of trimethoprim-sulfamethoxazole. Clin Infect Dis author reply 2 2011; 52: 1471-2. Available at: https://doi.org/10.1093/cid/cir247.

131. Chua K, Laurent F CG et al. Antimicrobial resistance: not community-associated methicillin-resistant Staphylococcus aureus (CA-MRSA)! A clinician’s guide to community MRSA—its evolving antimicrobial resistance and implications for therapy. Clin Infect Dis 2011; 52: 99-114. Available at: https://doi.org/10.1093/cid/ciq067.

132. Marais E, Aithma N, Perovic O, Oosthuysen WF, Musenge E DA. Antimicrobial susceptibility of methicillin-resistant Staphylococcus aureus isolates from South Africa.. SAMJ 2009; 99: 170-3.

133. Shittu AO LJ. Antimicrobial susceptibility patterns and characterization of clinical isolates of Staphylococcus aureus in KwaZulu-Natal province, South Africa. BMC Infect Dis 2006; 6:125. Available at: https://doi.org/10.1186/1471-2334-6-125.

134. Wichelhaus TA, Schafer V, Brade V BB. Molecular characterization of rpoB mutations conferring cross-resistance to rifamycins on methicillin-resistant Staphylococcus aureus. Antimicrob Agents Chemother 1999; 43: 2813-2816.

135. Perovic O, Singh-Moodley A, Govender NP, Kularatne R, Whitelaw A, Chibabhai V, Naicker P, Mbelle N, Lekalakala R, Quan V SC. A small proportion of community-associated methicillin-resistant Staphylococcus aureus bacteraemia, compared to healthcare-associated cases, in two South African provinces. Eur J Clin Microbiol Infect Dis 2017; 36(12): 25191-32. Available at: DOI 10.1007/s10096-017-3096-3.

136. Amoako DG, Bester LA, Somboro AM, Baijnath S, Govind CN, Essack SY. Plasmid mediated resistance and virulence mechanisms in the private health sector in KwaZulu-Natal, South Africa: An investigation of methicillin resistant Staphylococcus aureus (MRSA) clinical isolates col. Int J Infect Dis 2016; 46: 38-41.Available at:https://doi.org/10.1016/j.ijid.2016.03.019.

137. Sit PS, Teh CS, Idris N, Sam IC, Syed Omar SF SH, Thong KL, Kamarulzaman A PS. Prevalence of methicillin-resistant Staphylococcus aureus (MRSA) infection and the molecular characteristics of MRSA bacteraemia over a two-year period in a tertiary teaching hospital in Malaysia. BMC Infect Dis 2017; 17: 274. Available at:https://doi.org/10.1186/s12879-017-2384-y.

138. Antiabong JF, Kock MM, Bellea NM EM. Diversity of Multidrug Efflux Genes and Phenotypic Evaluation of the In vitro Resistance Dynamics of Clinical Staphylococcus Aureus Isolates Using Methicillin; a Model β-lactam. open Microbiol journal 2017; 11: 132-41. Available at: doi: 10.2174/1874285801711010132.

139. Conceiçao T, Santos Silva I, de Lencastre H A-SM. Staphylococcus aureus nasal carriage among patients and health care workers in Sao Tome and Principe. Microb Drug Resist 2014; 20(1): 57-66. Available at: https://doi.org/10.1089/mdr.2013.0136.

140. Sadowy E, Matynia B HW. Population structure, virulence factors and resistance determinants of invasive, noninvasive and colonizing Streptococcus agalactiae in Poland. J Antimicrob Chemother 2010; 65(9): 1907-1914.

141. Bohnsack JF, Whiting A, GottschalkM, DunnDM, Weiss R A. PH, Philips JB 3rd, Weisman LE, Rhoads GG LF. Population structure of invasive and colonizing strains of Streptococcus agalactiae from neonates of six U.S. Academic Centers from 1995 to 1999. J Clin Microbiol 2010; 46(4): 1285-1291.

142. Verani, J. R., McGee, L., Schrag SJ. Prevention of perinatal group B streptococcal disease— revised guidelines from CDC, 2010. MMWR Recomm Rep 2010; 59 (RR-10): 1-36.

143. Rachdi M, Boubaker IB, Hraoui M RS. High rates of macrolide resistance among clinical isolates of Streptococcus agalactiae in Tunisia. Arch Inst Pasteur Tunis 2010; 87(1/2): p.35.

144. Hraoui M, Boubaker IB, Doloy A, Redjeb SB BA. ‘Molecular mechanisms of tetracycline and macrolide resistance and emm characterization of Streptococcus pyogenes isolates in Tunisia.’ Microb Drug Resist 2011; 17, no. 3: 377-382. Available at:https://doi.org/10.1089/mdr.2010.0160.

145. Ksia S, Smaoui H, Hariga D KA. Biotypes and antimicrobial susceptibility of Streptococcus pyogenes strains isolated in children in Tunis. Bull la Société Pathol Exot 2010; 103(2): pp.69-74. Available at: http://www.springerlink.com/…/.

146. Fischer A, Liljander A, Kaspar H, Muriuki C, Fuxelius HH, Bongcam-Rudloff E, de Villiers EP, Huber CA, Frey J, Daubenberger C BR. ‘Camel Streptococcus agalactiae populations are associated with specific disease complexes and acquired the tetracycline resistance gene tetM via a Tn 916-like element.’ Vet Res 2013; 44, no. 1: 86. Available at: https://doi.org/10.1186/1297-9716-44-86.

147. Bolukaoto JY, Monyama CM, Chukwu MO, Lekala SM, Nchabeleng M, Maloba MR,Mavenyengwa RT, Lebelo SL, Monokoane ST, Tshepuwane C MS. Antibiotic resistance of Streptococcus agalactiae isolated from pregnant women in Garankuwa, South Africa. BMC Res Notes 2015: 6-12. Available at: https://doi.org/10.1186/s13104-015-1328-0.

148. Djahmi N, Boutet-Dubois A, Nedjai S, Dekhil M, Sotto A LJ. Molecular epidemiology of Enterococcus sp. isolated in a university hospital in Algeria. Scand J Infect Dis 2012; 44(9),.: pp.656-662. Available at: https://doi.org/10.3109/00365548.2012.673232.

149. Matsushima A, Takakura S, Yamamoto M, Matsumura Y, Shirano M, Nagao M, Ito Y, Iinuma Y, Shimizu T, Fujita N IS. Regional spread and control of vancomycin-resistant Enterococcus faecium and Enterococcus faecalis in Kyoto, Japan. Eur J Clin Microbiol Infect Dis 2011. Available at: DOI 10.1007/s10096-011-1412-x.

150. Fallico L, Boldrin C, Grossato A, Franchin E DCE, Tommasini T et al. Molecular epidemiology of Enterococcus faecium isolates from an Italian hospital. Infection 2011; 39: 127-33. Available at: DOI 10.1007/s15010-011-0086-8.

151. Hsieh YC, Lee WS, Ou TY HP. Clonal spread of CC17 vancomycin-resistant Enterococcus faecium with multilocus sequence type 78 (ST78) and a novel ST444 in Taiwan. Eur J Clin Microbiol Infect Dis 2010; 29: 25-30. Available at: DOI 10.1007/s10096-009-0810-9.

153. Iweriebor, Benson C., Larry C. Obi and AIO. ‘Virulence and antimicrobial resistance factors of Enterococcus spp. isolated from fecal samples from piggery farms in Eastern Cape, South Africa.’ BMC Microbiol 15, no. 1: 136. Available at: https://doi.org/10.1186/s12866-015-0468-7.

154. Djahmi N, Messad N, Nedjai S, Moussaoui A, Mazouz D, Richard JL, Sotto A LJ. Molecular epidemiology of Staphylococcus aureus strains isolated from inpatients with infected diabetic foot ulcers in an Algerian University Hospital. Clin Microbiol Infect 2013; 19(9):E398-404. Available at: https://doi.org/10.1111/1469-0691.12199.

155. Conceiçao T, Coelho C, Santos Silva I, de Lencastre H A-SM. Methicillin-resistant Staphylococcus aureus in the community in Luanda, Angola: blurred boundaries with the hospital setting. Microb Drug Resist 2016; 22: 22-7. Available at:https://doi.org/10.1089/mdr.2015.0236.

156. Abbassi MS, Znazen A, Mahjoubi F, Hammami A BA. Emergence of vancomycin-resistant Enterococcus faecium in Sfax: clinical features and molecular typing. Med Mal Infect 2007; 37: 240-1. Available at: doi: 10.1016/j.medmal.2006.09.011.

157. Abdelkareem MZ, Sayed M, Hassuna NA, Mahmoud MS AS. Multi-drug-resistant Enterococcus faecalis among Egyptian patients with urinary tract infection. J Chemother 2017; 29: 74-82. Available at: https://doi.org/10.1080/1120009X.2016.1182358.

158. Abouelfetouh A, Kassem M, Naguib M E-NM. Investigation and Treatment of Fusidic Acid Resistance Among Methicillin-Resistant Staphylococcal Isolates from Egypt. Microb Drug Resist 2017; 23: 8-17. Available at:https://doi.org/10.1089/mdr.2015.0336.

159. Bendary MM, Solyman SM, Azab MM, Mahmoud NF HA. Genetic diversity of multidrug resistant Staphylococcus aureus isolated from clinical and non clinical samples in Egypt. Cell Mol Biol 2016; 62: 55-61. Available at: doi: 10.14715/cmb/2016.62.10.9.

160. Bendary MM, Solyman SM, Azab MM, Mahmoud NF HA. Characterization of Methicillin Resistant Staphylococcus aureus isolated from human and animal samples in Egypt. Cell Mol Biol 2016; 62: 94-100. Available at: doi: 10.14715/cmb/2016.62.2.16.

161. El-Sharif A AH. Community-acquired methicillin-resistant Staphylococcus aureus (CA-MRSA) colonization and infection in intravenous and inhalational opiate drug abusers. Exp Biol Med 2008; 233: 874-80. Available at: https://doi.org/10.3181/0711-RM-294.

162. Okuda KV, Toepfner N, Alabi AS, Arnold B, Bélard S, Falke U, Menschner L, Monecke S, Ruppelt-Lorz A BR. Molecular epidemiology of Staphylococcus aureus from Lambarene, Gabon. Eur J Clin Microbiol Infect Dis 2016; 35: 1963-73. Available at: DOI 10.1007/s10096-016-2748-z.

163. Egyir B, Guardabassi L, Nielsen SS, Larsen J, Addo KK, Newman MJ LA. Prevalence of nasal carriage and diversity of Staphylococcus aureus among inpatients and hospital staff at Korle Bu Teaching Hospital, Ghana. J Glob Antimicrob Resist 2013; 1: 189-93. Available at: https://doi.org/10.1016/jjgar.2013.05.006.

164. Maina EK, Kiiyukia C, Wamae CN, Waiyaki PG KS. Characterization of methicillin-resistant Staphylococcus aureus from skin and soft tissue infections in patients in Nairobi, Kenya. Int J Infect Dis 2013; 17: e115-9. Available at:https://doi.org/10.1016/j.ijid.2012.09.006.

165. Al-haddad OH, Zorgani A GK. Nasal Carriage of Multi-Drug Resistant Panton-Valentine Leucocidin-Positive Methicillin-Resistant Staphylococcus aureus in Children in Tripoli-Libya. Am J Trop Med Hyg 2014; 90: 724-7. Available at: doi: https://doi.org/10.4269/ajtmh.13-0715.

166. Zriouil SB, Bekkali M ZK. Epidemiology of Staphylococcus aureus infections and nasal carriage at the Ibn Rochd University hospital center, Casablanca, Morocco. Brazilian J Infect Dis 2012 May 1;16(3)279-83 2012; 16: 279-83. Available at: https://doi.org/10.1016/S1413-8670(12)70324-0.

167. Ayepola OO, Olasupo NA, Egwari LO, Becker K. Molecular Characterization and Antimicrobial Susceptibility of Staphylococcus aureus Isolates from Clinical Infection and Asymptomatic Carriers in Southwest Nigeria. PLoS One 2015; 2304: 4-11. Available at: http://dx.doi.org/10.1371/journal.pone.0137531.

168. Kolawole DO, Adeyanju A, Schaumburg F, Akinyoola AL, Lawal OO, Amusa YB, Köck R BK. Characterization of colonizing Staphylococcus aureus isolated from surgical wards’ patients in a Nigerian university hospital. PLoS One 2013; 8(7): e68721.

169. Shittu AO, Okon K, Adesida S, Oyedara O, Witte W, Strommenger B, Layer F NU. Antibiotic resistance and molecular epidemiology of Staphylococcus aureus in Nigeria. BMC Microbiol 2011; 11: 92. Available at: https://doi.org/10.1186/1471-2180-11-92.

170. Okon KO, Basset P, Uba A, Lin J, Oyawoye B, Shittu AO BD. Cooccurrence of predominant Panton-Valentine leukocidin-positive sequence type (ST) 152 and multidrug-resistant ST 241 Staphylococcus aureus clones in Nigerian hospitals. J Clin Microbiol 47: 3000-3. Available at: doi: 10.1128/JCM.01119-09.

171. Ghebremedhin B, Olugbosi MO, Raji AM, Layer F, Bakare RA, König B KW. Emergence of a community-associated methicillin-resistant Staphylococcus aureus strain with a unique resistance profile in Southwest Nigeria. J Clin Microbiol 2009; 47: 2975-80. Available at: doi: 10.1128/JCM.00648-09.

172. Mbelle NM, Maningi NE, Tshisevhe V, Modipane L, Amoako DG SJ. First Report of a Whole-Genome Shotgun Sequence of a Clinical Enterococcus faecalis Sequence Type 6 Strain from South Africa. Genome Announc 2017; 5(50): e01382-17. Available at: doi: 10.1128/genomeA.01382-17.

173. Mbelle NM, Maningi NE, Tshisevhe V, Modipane L, Amoako DG SJ. Draft Genome Sequence of a Clinical Enterococcus faecium Sequence Type 18 Strain from South Africa. Genome Announc 2017; 5(48): e01381-17. Available at:https://doi.org/10.1128/genomeA.01381-17.

175. Shittu AO, Udo EE LJ. Phenotypic and molecular characterization of Staphylococcus aureus isolates expressing low-and high-level mupirocin resistance in Nigeria and South Africa. BMC Infect Dis 2009; 9: 10. Available at: https://doi.org/10.1186/1471-2334-9-10.

176. Shittu A, Lin J MD. Molecular identification and characterization of mannitol-negative methicillin-resistant Staphylococcus aureus. Diagnostic Microbiol Infect Dis 57(1): 93-5. Available at: https://doi.org/10.1016/j.diagmicrobio.2006.05.004.

177. Elhassan MM, Ozbak HA, Hemeg HA, Elmekki MA AL. Absence of the mecA gene in methicillin resistant Staphylococcus aureus isolated from different clinical specimens in shendi city, Sudan. BioMed Res Int 2015: 2015. Available at: http://dx.doi.org/10.1155/2015/895860.

179. Nejma MB, Mastouri M, Jrad BB NM. Characterization of ST80 Panton-Valentine leukocidin-positive community-acquired methicillin-resistant Staphylococcus aureus clone in Tunisia. Diagnostic Microbiol Infect Dis 2013; 77: 20-4.Available at:https://doi.org/10.1016/j.diagmicrobio.2008.02.010.

180. Bouchami O, Achour W, Mekni MA, Rolo J HA. Antibiotic resistance and molecular characterization of clinical isolates of methicillin-resistant coagulase-negative staphylococci isolated from bacteremic patients in oncohematology. Folia Microbiol 56: 122. Available at: DOI 10.1007/s12223-011-0017-1.

181. Slama KB, Gharsa H, Klibi N, Jouini A, Lozano C, Gómez-Sanz E, Zarazaga M, Boudabous A TC. Nasal carriage of Staphylococcus aureus in healthy humans with different levels of contact with animals in Tunisia: genetic lineages, methicillin resistance, and virulence factors. Eur J Clin Microbiol Infect Dis 2011; 30: 499-508. Available at: DOI 10.1007/s10096-010-1109-6.

182. Bouchami O, Achour W HA. ‘Prevalence of resistance phenotypes and genotypes to macrolide, lincosamide and streptogramin antibiotics in Gram-positive cocci isolated in Tunisian Bone Marrow Transplant Center.’ Pathol Biol 2011; 59, no. 4: 199-206. Available at: https://doi.org/10.1016/j.patbio.2009.03.010.

183. Bouchami O, Achour W HA. Typing of staphylococcal cassette chromosome mec encoding methicillin resistance in Staphylococcus aureus strains isolated at the bone marrow transplant centre of Tunisia. Curr Microbiol 2009; 59: 380-5. Available at: DOI 10.1007/s00284-009-9448-1.

184. Klibi N, Ben Slama K, Sáenz Y, Masmoudi A, Zanetti S, Sechi LA, Boudabous A TC. Detection of virulence factors in high-level gentamicin-resistant Enterococcus faecalis and Enterococcus faecium isolates from a Tunisian hospital. Can J Microbiol 2007; 53(3). Available at: https://doi.org/10.1139/W06-136.

185. Achour W, Bouchami O, Galopin S, Leclercq R BHA. Analysis of pristinamycin resistant Staphylococcus epidermidis isolates in the Tunisian Bone Marrow Transplant Center. Lett Appl Microbiol 46: 358-63. Available at: https://doi.org/10.1111/j.1472-765X.2007.02318.x.

186. Bouchami O, Achour W HA. Prevalence and mechanisms of macrolide resistance among Staphylococcus epidermidis isolates from neutropenic patients in Tunisia. Clin Microbiol Infect 2007; 13: 103-6.Available at: https://doi.org/10.1111/j.1469-0691.2006.01567.x.

187. Seni J, Bwanga F, Najjuka CF, Makobore P, Okee M, Mshana SE, Kidenya BR, Joloba ML KD. ‘Molecular characterization of Staphylococcus aureus from patients with surgical site infections at Mulago Hospital in Kampala, Uganda.’ PLoS One 2013; 8, no. 6: e66153.

188. Okee MS, Joloba ML, Okello M, Najjuka FC, Katabazi FA, Bwanga F, Nanteza A KD. Prevalence of virulence determinants in Staphylococcus epidermidis from ICU patients in Kampala, Uganda. J Infect Dev Countries 2011;6: 242-50. Available at:https://doi.org/10.3855/jidc.2007.

189. Kitara LD, Anywar AD, Acullu D, Odongo-Aginya E, Aloyo J FM. Antibiotic susceptibility of Staphylococcus aureus in suppurative lesions in Lacor Hospital, Uganda. African Heal Sci 11: 34-9. Available at: http://dx.doi.org/10.4314/ahs.v11i3.70068.

190. Ojulong J, Mwambu TP, Joloba M, Bwanga F K-MD. Relative prevalence of methicilline resistant Staphylococcus aureus and its susceptibility pattern in Mulago Hospital, Kampala, Uganda. Tanzania J Heal Res 2009; 11. Available at:http://dx.doi.org/10.4314/thrb.v11i3.47703.

191. Martins E, Novais C, Freitas AR, Dias AR, Ribeiro TG, Antunes P PL. Filling the map for antimicrobial resistance in sub-Saharan Africa: ampicillin-resistant Enterococcus from non-clinical sources in Angola. J Antimicrob Chemother 70: 2914-6.Available at:doi:10.1093/jac/dkv172.

192. Awad A, Ramadan H, Nasr S, Ateya A AS. Genetic Characterization Antimicrobial Resistance Patterns And Virulence Determinants of Staphylococcus aureus Isolated Form Bovine Mastitis. Pak J Biol Sci 2017; 20: 298-305. Available at: https://scialert.net/abstract/?doi=pjbs.2017.298.305.

193. Ammar AM, Attia AM, Abd EH, MI ES. IM and Abd El-Kader S. Genetic basis of resistance waves among methicillin resistant Staphylococcus aureus isolates recovered from milk and meat products in Egypt. Cell Mol Biol Noisy-le-Grand 2016; 621: 7-15.

194. Sallam KI, Abd-Elghany SM, Elhadidy M TT. Molecular characterization and antimicrobial resistance profile of methicillin-resistant Staphylococcus aureus in retail chicken. J food Prot 2015; 78: 1879-84. Available at: https://doi.org/10.4315/0362-028X.JFP-15-150.

195. Fischer A, Liljander A, Kaspar H, Muriuki C, Fuxelius HH, Bongcam-Rudloff E, de Villiers EP, Huber CA, Frey J, Daubenberger C BR. Camel Streptococcus agalactiae populations are associated with specific disease complexes and acquired the tetracycline resistance gene tetM via a Tn 916-like element. Vet Res 2013; 44(1): 86. Available at: https://doi.org/10.1186/1297-9716-44-86.

196. Ngbede EO, Raji MA, Kwanashie CN KJ. Antimicrobial resistance and virulence profile of enterococci isolated from poultry and cattle sources in Nigeria. Trop Anim Heal Prod 2017; 49(3): 451-8. Available at: DOI 10.1007/s11250-016-1212-5.

197. Chah KF, Gómez-Sanz E, Nwanta JA, Asadu B, Agbo IC, Lozano C, Zarazaga M TC. Methicillin-resistant coagulase-negative staphylococci from healthy dogs in Nsukka, Nigeria. Brazilian J Microbiol 2014; 45: 215-20. Available at: http://dx.doi.org/10.1590/S1517-83822014005000034.

198. Iweriebor BC, Obi LC OA. Macrolide, glycopeptide resistance and virulence genes in Enterococcus species isolates from dairy cattle. J Med Microbiol 2016; 65: 641-87.

199. Soyege AO, Ajayi AO, Ngwenya E, Basson AK OA. Vancomycin and Oxacillin CoResistance of Commensal Staphylococci. Jundishapur J Microbiol 2014; 7. Available at: doi: 10.5812/jjm.9310.

200. Freitas AR, Elghaieb H, León-Sampedro R, Abbassi MS, Novais C, Coque TM, Hassen A PL. Detection of optrA in the African continent (Tunisia) within a mosaic Enterococcus faecalis plasmid from urban wastewaters. J Antimicrob Chemother 2017; 72(12): 3245-51. Available at: https://doi.org/10.1093/jac/dkx321.

201. Chairat S, Gharsa H, Lozano C, Gómez-Sanz E, Gómez P, Zarazaga M, Boudabous A, Torres C BSK. Characterization of Staphylococcus aureus from raw meat samples in Tunisia: detection of clonal lineage ST398 from the African continent. Foodborne Pathog Dis 2015; 12: 686-92. Available at: https://doi.org/10.1089/fpd.2015.1958.

202. Gharsa H, Slama KB, Gómez-Sanz E, Lozano C, Zarazaga M, Messadi L, Boudabous A TC. Molecular characterization of Staphylococcus aureus from nasal samples of healthy farm animals and pets in Tunisia. Vector-Borne Zoonotic Dis 2015; 15: 109-15. Available at: https://doi.org/10.1089/vbz.2014.1655.

203. Gharsa H, Sallem RB, Slama KB, Gómez-Sanz E, Lozano C, Jouini A, Klibi N, Zarazaga M, Boudabous A TC. High diversity of genetic lineages and virulence genes in nasal Staphylococcus aureus isolates from donkeys destined to food consumption in Tunisia with predominance of the ruminant associated CC133 lineage. BMC Vet Res 2012; 8: 203. Available at: https://doi.org/10.1186/1746-6148-8-203.

204. Akanbi OE, Njom HA, Fri J, Otigbu AC CA. Antimicrobial Susceptibility of Staphylococcus aureus Isolated from Recreational Waters and Beach Sand in Eastern Cape Province of South Africa. Int J Environ Res public Heal 2017; 14(9): 1001. Available at: doi: 10.3390/ijerph14091001.

205. Molale LG BC. Antibiotic resistance, efflux pump genes and virulence determinants in Enterococcus spp. from surface water systems. Environ Sci Pollut Res 23: 21501-10. Available at: DOI 10.1007/s11356-016-7369-7.

206. Gharsa H, Dziri R, Klibi N, Chairat S, Lozano C, Torres C, Bellaaj R SK. Environmental Staphylococcus aureus contamination in a Tunisian hospital. J Chemother 2016; 28: 506-9. Available at: https://doi.org/10.1179/1973947815Y.0000000036.

